# Universal DNA methylation age across mammalian tissues

**DOI:** 10.1101/2021.01.18.426733

**Authors:** A.T. Lu, Z. Fei, A. Haghani, T.R. Robeck, J.A. Zoller, C.Z. Li, R. Lowe, Q. Yan, J. Zhang, H. Vu, J. Ablaeva, V.A. Acosta-Rodriguez, D.M. Adams, J. Almunia, A. Aloysius, R. Ardehali, A. Arneson, C.S. Baker, G. Banks, K. Belov, N.C. Bennett, P. Black, D.T. Blumstein, E.K. Bors, C.E. Breeze, R.T. Brooke, J.L. Brown, G. Carter, A. Caulton, J.M. Cavin, L. Chakrabarti, I. Chatzistamou, H. Chen, K. Cheng, P. Chiavellini, O.W. Choi, S. Clarke, L.N. Cooper, M.L. Cossette, J. Day, J. DeYoung, S. DiRocco, C. Dold, E.E. Ehmke, C.K. Emmons, S. Emmrich, E. Erbay, C. Erlacher-Reid, C.G. Faulkes, S.H. Ferguson, C.J. Finno, J.E. Flower, J.M. Gaillard, E. Garde, L. Gerber, V.N. Gladyshev, V. Gorbunova, R.G. Goya, M.J. Grant, C.B. Green, E.N. Hales, M.B. Hanson, D.W. Hart, M. Haulena, K. Herrick, A.N. Hogan, C.J. Hogg, T.A. Hore, T. Huang, J.C. Izpisua Belmonte, A.J. Jasinska, G. Jones, E. Jourdain, O. Kashpur, H. Katcher, E. Katsumata, V. Kaza, H. Kiaris, M.S. Kobor, P. Kordowitzki, W.R. Koski, M. Kruetzen, S.B. Kwon, B. Larison, S.G. Lee, M. Lehmann, J.F. Lemaitre, A.J. Levine, C. Li, X. Li, A.R. Lim, D.T.S. Lin, D.M. Lindemann, T.J. Little, N. Macoretta, D. Maddox, C.O. Matkin, J.A. Mattison, M. McClure, J. Mergl, J.J. Meudt, G.A. Montano, K. Mozhui, J. Munshi-South, A. Naderi, M. Nagy, P. Narayan, P.W. Nathanielsz, N.B. Nguyen, C. Niehrs, J.K. O’Brien, P. O’Tierney Ginn, D.T. Odom, A.G. Ophir, S. Osborn, E.A. Ostrander, K.M. Parsons, K.C. Paul, M. Pellegrini, K.J. Peters, A.B. Pedersen, J.L. Petersen, D.W. Pietersen, G.M. Pinho, J. Plassais, J.R. Poganik, N.A. Prado, P. Reddy, B. Rey, B.R. Ritz, J. Robbins, M. Rodriguez, J. Russell, E. Rydkina, L.L. Sailer, A.B. Salmon, A. Sanghavi, K.M. Schachtschneider, D. Schmitt, T. Schmitt, L. Schomacher, L.B. Schook, K.E. Sears, A.W. Seifert, A. Seluanov, A.B.A. Shafer, D. Shanmuganayagam, A.V. Shindyapina, M. Simmons, K. Singh, I. Sinha, J. Slone, R.G. Snell, E. Soltanmaohammadi, M.L. Spangler, M.C. Spriggs, L. Staggs, N. Stedman, K.J. Steinman, D.T. Stewart, V.J. Sugrue, B. Szladovits, J.S. Takahashi, M. Takasugi, E.C. Teeling, M.J. Thompson, B. Van Bonn, S.C. Vernes, D. Villar, H.V. Vinters, M.C. Wallingford, N. Wang, R.K. Wayne, G.S. Wilkinson, C.K. Williams, R.W. Williams, X.W. Yang, M. Yao, B.G. Young, B. Zhang, Z. Zhang, P. Zhao, Y. Zhao, W. Zhou, J. Zimmermann, J. Ernst, K. Raj, S. Horvath

## Abstract

Aging is often perceived as a degenerative process resulting from random accrual of cellular damage over time. Despite this, age can be accurately estimated by epigenetic clocks based on DNA methylation profiles from almost any tissue of the body. Since such pan-tissue epigenetic clocks have been successfully developed for several different species, we hypothesized that one can build pan-mammalian clocks that measure age in all mammalian species. To address this, we generated data using 11,754 methylation arrays, each profiling up to 36 thousand cytosines in highly-conserved stretches of DNA, from 59 tissue-types derived from 185 mammalian species. From these methylation profiles, we constructed three age predictors, each with a single mathematical formula, termed universal pan-mammalian clocks that are accurate in estimating the age (r>0.96) of any mammalian tissue. Deviations between epigenetic age and chronological age relate to mortality risk in humans, mutations that affect the somatotropic axis in mice, and caloric restriction. We characterized specific cytosines, whose methylation levels change with age across most mammalian species. These cytosines are greatly enriched in polycomb repressive complex 2-binding sites, are located in regions that gradually lose chromatin accessibility with age and are proximal to genes that play a role in mammalian development, cancer, human obesity, and human longevity. Collectively, these results support the notion that aging is indeed evolutionarily conserved and coupled to developmental processes across all mammalian species - a notion that was long-debated without the benefit of this new compelling evidence.

**SUMMARY:** This study identifies and characterizes evolutionarily conserved cytosines implicated in the aging process across mammals and establishes pan mammalian epigenetic clocks.

## INTRODUCTION

Aging is associated with multiple cellular changes that are often tissue-specific ^1^. Cytosine methylation, however, is unusual in this regard, as some of it changes in similar ways in different aging tissues. This allows for the development of pan-tissue aging clocks (multivariate age-estimators) that are applicable to all human tissues^2–4^.The subsequent development of similar pan-tissue clocks for mice and other species hint at the universality of the aging process^5–7^, and questions the comprehensiveness of the notion that aging is driven purely by random cellular damage accrued over time. To investigate this, we sought to: i) develop universal age-estimators that apply to all mammalian species and tissues (pan-mammalian clocks), ii) identify and characterize cytosines whose methylation levels change with age in all mammals. To these ends, we employed the mammalian methylation array, which we recently developed to profile methylation levels of up to 36,000 CpGs with flanking DNA sequences that are highly-conserved across the mammalian class^8^. We obtained such profiles from 11,754 samples from 59 tissue types, derived from 185 mammalian species, representing 19 taxonomic orders (**Data S1.1–1.4, Supplementary information, notes 1&2**) and ranging in age from prenatal to 139 years old (bowhead whale).

## RESULTS

### Universal pan-mammalian epigenetic clocks

In separate articles, we described the application of the mammalian methylation array to individual mammalian species ^9–17^. These studies already demonstrate that one can build dual species epigenetic age estimators (e.g. human-naked mole rat clocks), which we refer to as third generation epigenetic clocks ^9–16^, in contrast to first and second generation clocks that measure human age ^4,18,19^ and mortality risk ^20,21^, respectively. However, it is not yet known whether one can develop a mathematical formula to estimate age in all mammalian species. Here we present three such pan-mammalian age-estimators.

The first, *basic clock* (*Clock 1*), directly regresses log-transformed chronological age on DNA methylation levels of all available mammals. Although such a clock can directly estimate the age of any mammal, its usefulness could be further increased if its output was adjusted for differences in maximum lifespan of each species as well, as this would allow biologically meaningful comparisons to be made between species with very different lifespans. To this end, we developed a second universal clock that defines individual age relative to the maximum lifespan of its species; generating *relative age* estimates between 0 and 1. Since the accuracy of this *universal relative age clock* (*Clock 2*) could be compromised in species for which knowledge of maximum lifespan is inaccurate, we developed a third universal clock, which substitutes species maximum lifespan for species age at sexual maturity and gestation time because these traits are more easily established. Age at sexual maturity and gestation time explain more than 69% of the variation in maximum lifespan on the log scale (**Data S2**). This third clock is referred to as the *universal log-linear transformed age clock* (*Clock 3*). The non-linear mathematical functions underlying the age transformations of both Clock 2 and Clock 3 reflect the fact that epigenetic clocks tick faster during development, an observation that led to the establishment of the first pan-tissue clock for humans ^4^ (**Figures S1a-b & de**).

### Performance of universal epigenetic clocks across species

We employed two different strategies for evaluating the accuracy of the clocks. First, the leave-one-fraction-out (LOFO) cross-validation analysis divided the data set into 10 fractions, each of which contained the same proportions of species and tissue types, and a different fraction is left out for validation at each iteration of analysis. Second, the leave-one-species-out analysis (LOSO) was similarly cross-validated but with the omission of a species at each iteration instead. The final prediction models of the three clocks are composed of fewer than 1,000 CpGs each, as listed in **Data S3.1–3.3**. Of those, 401 common genes are proximal to CpGs that are constituents of both Clock 2 and Clock 3, (**Data S3.5**).

According to LOFO cross-validation, the epigenetic clocks are remarkably accurate estimators of chronological age across all mammalian species (r~0.96–0.98), with a median error of <1 year and a median relative error of <3.3% (**Figures1a,c, & 2, Figure S2a,Table S2, Data S4.1–4.3**). The clocks are also applicable to marsupials (r=0.908, med.MAE <0.8 year for Clock 2, **Figure 1b**) even though fewer CpGs on the mammalian array map to marsupials^8^. While our study of monotremes was underpowered (n=15), we nevertheless obtained very encouraging results (e.g., r=0.85 for Clock 2 **Data S4.1**). Employing the LOSO cross-validation, the performance of the clocks achieved age correlations up to r=0.941 (**Table S2**) which indicates that these clocks will apply to species that were not part of the training set. However, for some species, such as bowhead whales, the epigenetic age as predicted by the basic clock accords poorly with chronological age (**Figure S2b**).

**Figure 1.**
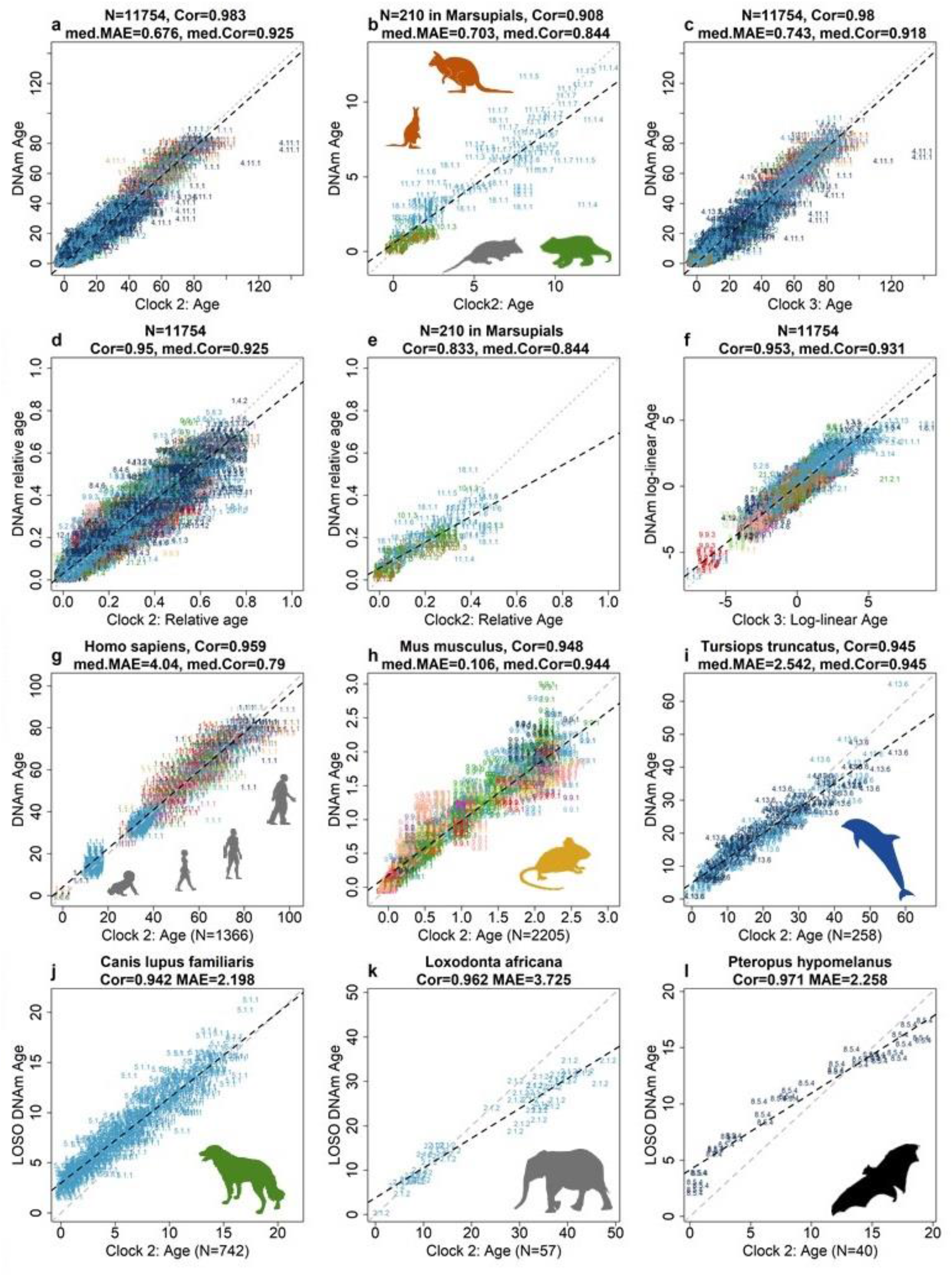
Universal clocks for transformed age across mammals. The figure displays relative age estimates of universal clock 2 (Clock 2), and log-linear transformed age of universal clock 3 (Clock3). Relative age estimates incorporates maximum lifespan, and assumes values between 0 and 1. Log-linear age is formulated with age at sexual maturity and gestational time. **a–i**, Age estimated via leave-one-fraction-out (LOFO) cross-validation for Clock 2 and Clock 3. **j–l**, age estimated via leave-one-species-out (LOSO) cross-validation for Clock 2. The DNAm estimates of age (y axes) of (**a–b**) and (**c**) are transformations of relative age (Clock 2) or log-linear age (Clock 3) into units of years. **b,e**, Only marsupials (eight species). Each panel reports Pearson correlation coefficient. Median correlation (med.Cor) and median of median absolute error (med.MAE) are calculated across species (**a–f**) or across species-tissue (**g–l**). All correlation P-values are highly significant (P<1.0×10^-22^). Each sample is labeled by mammalian species index and indicated by tissue color (**Data S1.3–1.4**).

**Figure 2.**
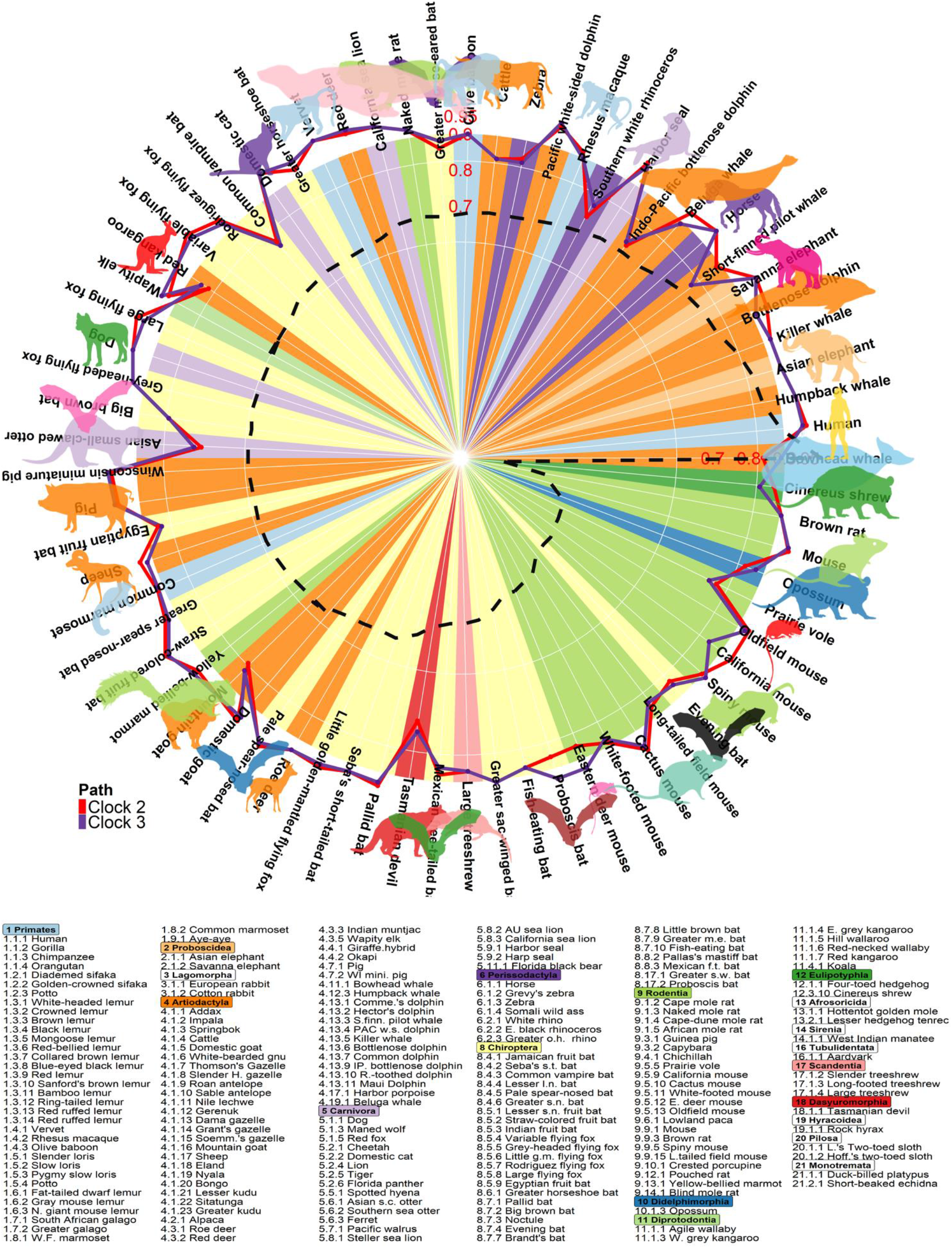
Accuracy of universal clocks are independent of species lifespan. The circle plot displays Pearson correlation between age and DNAm Age estimated by universal clock 2 (Clock 2) and 3 (Clock 3) for various species. Of the 185 species, correlation analysis was performed on 69 of them (with n ≥15 in a single tissue), across 12 taxonomic orders. We took log transformation of maximum lifespans of species and divided them by log (211), which is the maximum lifespan of bowhead whales. Values of the resulting ratios ranged from 0.12 (cinerus shrew) to one (bowhead whales). These ratios are displayed in descending order, in the circle plot marked by dark grey dash line, starting with bowhead whale (1), human (0.90) and ending with cinerus shrew (0.12), in counter clockwise direction. In the background, circumferences with increasing radii represent increasing correlation levels up to 0.9. These correlations between age and DNAm Age were estimated by Clock 2 (red path line) and Clock 3 (purple path line) for each species. Colors within the circle represent taxonomic order of the corresponding species, as listed below the circle. The median of correlation across species is 0.926 for Clock 2 and 0.918 for Clock 3. Straw-colored fruit bats exhibit the highest correlation (r=0.985) based on Clock 2 and Wisconsin miniature pigs have the second highest correlation (r=0.984) based on Clock 3. A majority of species with their circle lines located outside the background indicate that their correlation estimates are greater than 0.9. The text at the bottom list the 185 species under their corresponding taxonomic order. Each taxonomic order is marked by the same color matching with the circle plot. The numbers after the first and second decimal points enumerate the taxonomic family and species, respectively.

For the basic Clock 1, the mean discrepancy between LOSO DNAmAge and chronological age (Delta.Age) is negatively correlated with species maximum lifespan (r=-0.84, *P*=1.0×10^-19^) and species age at sexual maturity (r= −0.75, *P*=7.9×10^-14^, **Figure S2c–d**). Here, the strengths of Clock 2 and 3 come to fore as they adjust for these species characteristics during their construction (**Figure S1**). Clocks 2 and 3 achieve a correlation of r ≥ 0.95 between DNAm transformed age and observed transformed age, respectively (**Figure 1d,f**). Both of these clocks present comparably accurate LOFO estimates in numerous tissue types in 70 species (**Figures 1**, **Data S4.2)**. For example, Clock 2 leads to high age correlations in humans (LOFO estimate of r=0.959, 20 tissues), mice (r=0.948, 26 tissues), and bottlenose dolphins (r=0.945, 2 tissues). **Figure 2** displays the circle plots for the age correlation estimates in different species sorted by maximum lifespan.

We find that the age correlation resulting from clocks 2 and 3 are not related to maximum lifespan (denoted by the dash line in the circle of **Figure 2**). While the clocks accurately predicted the age for one mysticete species, the humpback whale, and all other mammalian species, the ages of bowhead samples were sometimes underestimated (mammalian species index 4.11.1 in **Figure 1a,c**). This could reflect an overestimation of the age of the older bowhead whales by the method of aspartic acid racemization rate. Clocks 2 and 3 are similarly accurate with LOSO age-estimates between evolutionarily distant species (**Data S5.2**) including dogs (n=742 blood samples from 93 dog breeds, r=0.942, MAE <2.28 years), African elephants (r=0.962, MAE<4 years), and flying foxes (r=0.971, MAE<2.3 years) (**Figure 1j–l**). Such high predictive accuracy of LOSO analyses demonstrates that these universal clocks tap into age-related mechanisms that are highly conserved across mammals and would therefore be applicable to any mammalian species, including those that are not part of the training data (**Data S5.1–5.2**). The three universal clocks performed just as well in 114 species, for which there were fewer than 15 samples per species (r~0.90, MAE~1.2 years, **Figure S3a–c**), showing very strong correlation between estimated and actual relative age (r=0.91, **Figure S3d**).

### Tissue specific pan-mammalian clocks

The universal pan-mammalian clocks were constructed on the basis of many tissue types, i.e. they are pan-tissue clocks. We also developed analogous clocks based solely on blood (Universal BloodClock 2 and Universal BloodClock 3) or skin (Universal SkinClock 2 and Universal SkinClock 3), which are the most accessible tissues across all species. These tissue-specific clocks are expected to be more accurate than the pantissue clocks when analyzing the respective tissues. Both blood and skin clocks exhibit high age correlations (r~0.98–0.99 for blood and r~0.95–0.97 for skin, **Figure S4c,g**).

### Pan-mammalian universal clocks across tissues

The markedly distinct epigenomic landscape across tissue types ^22,23^, raised the question of the performance of these clocks in different tissues. We assessed the tissue-specific accuracy of Clock 2 for relative age (r=0.95, **Figure 1d**) across 33 distinct tissue types and observed the median correlation to be 0.91 and median MAE for relative age to be 0.027 (**Data S4.3**). There was consistently high age-correlation in the brain regions: whole brain (r=0.991), cerebellum (r=0.963), cortex (r=0.957), hippocampus (r=0.954), and striatum (r=0.935, **Figures S5a,d,f,g&i** and **Data S4.3**) and organs: spleen (r=0.982), liver (r=0.963) and kidney (r=0.963, **Figures S5b,c&e**). Blood and skin also exhibited similarly high estimates of relative age correlations across different species: blood (r=0.952, MAE=0.022, 124 species) and skin (r=0.942, MAE=0.027, 92 species, **Figures S5h&k**). Below we demonstrate several potential applications of our novel universal clocks on human mortality study, human cellular reprogramming experiments and a variety of murine anti-aging studies.

### Human mortality risk

Retrospective epidemiological studies have shown that human epigenetic clocks predict time-to-death and human mortality risk even after adjusting for chronological age, sex, and other risk factors^21,24,25^. To test whether pan-mammalian methylation clocks can also do so, we used two large human retrospective cohort studies: Framingham heart study offspring cohort (FHS, n=2,544) and Women’s Health Initiative (WHI, n=2107). Our metaanalysis of Cox regression models shows that both pan-mammalian clocks 2 and 3 are predictive of human mortality risk even after adjusting for age. One year of epigenetic age acceleration was associated with a significant hazard ratio (HR) for all-cause mortality (HR=1.03 and P=6.0×10^-19^ for Clock 2 and HR=1.03, P=5.3×10^-11^, **Figure 3a—b**).

**Figure 3.**
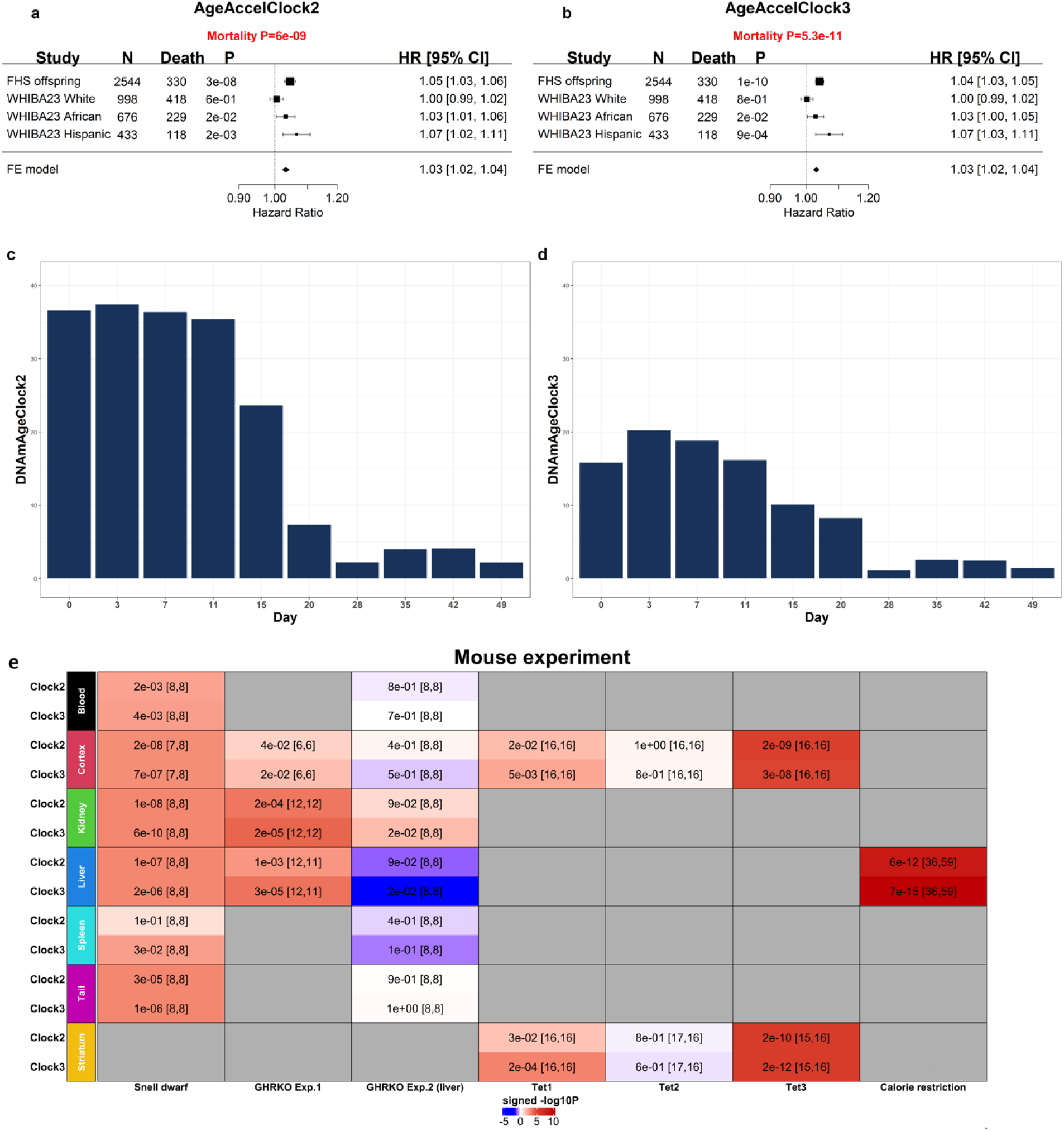
Applications of universal pan-mammalian clocks in human cohorts, reprogramming experiment, and murine anti-aging studies. (**a,b**) Meta-analysis forest plot combining hazard ratios calculated using Cox regression models for time-to-death based on epigenetic age acceleration (AgeAccel) measures of Clock 2 (AgeAccelClock2) and Clock 3 (AgeAccelClcok3), across different strata of human racial groups within two epidemiological cohorts (FHS and WHI). Each row reports a hazard ratio (for time-to-death) corresponding to a one-year increase in AgeAccel measures and a 95% confidence interval resulting from a Cox regression model. (**c,d**) DNAm age estimates of human dermal fibroblasts during OSKM-induced reprogramming. The y-axes are DNAm age estimates of Clock 2 and Clock 3 at day 0, 3, 7,…,42 and 49, respectively during reprogramming (Ohnuki et al., 2014). (**e**) Evaluations of mouse anti-age interventions: i) age-matched Snell dwarf mutation study: 48 normal and 47 Dwarf mice with ages approximately 0.52 (mean±SD of age =0.52±0.01 years), ii) age-matched whole body growth hormone receptor knock-out experiment 1 with 36 normal and 35 GHRKO mice (mean±SD of age is 0.65±0.06 years), iii) age-matched GHRKO experiment 2 with GHR knock-out in livers only with 48 normal and 48 GHRKO genotypes (mean±SD of age is 0.51±0.03 years old), iv) Tet gene knout-out study with all samples at age 0.5 years. Tet1: 32 controls and 32 Tet1 KO genotypes. Tet2: 33 controls and 32 Tet2 KO genotypes. Tet3: 31 control and 32 Tet3.KO genotypes, and v) calorie restriction (CR) study in livers: 59 in CR versus 36 control mice with all ages at 1.57 years old. The comparisons in the experiments ii) & iii) were based on the measures of epigenetic age acceleration, which were defined as raw residual resulting regressing DNAm age on chronological age. By definition, the resulting measure of epigenetic age acceleration (AgeAccel) is not correlated with chronological age. All the mice in experiment i) have approximately the same ages and in experiments iv) & v) have the same ages, respectively. There was no need to define a measure of epigenetic age acceleration in these experiments. The color gradient is based on the sign of t-test for controls versus experimental mice (Snell dwarf, GHRKO, CR and Tet3). A positive t-test indicates that the mice in the control group exhibit older DNAm age or higher age acceleration compared to the mice in the experimental group.

### OSKM-based reprogramming

Epigenetic clocks such as the pan-tissue clock for humans^4^ indicate that cellular reprogramming based on the Yamanaka factors (OSKM) leads to age reversal^4,26^. To assess whether the universal clocks exhibit the same age reversal pattern during reprogramming, we applied Clock 2 and Clock 3 to a previously published reprogramming dataset in human dermal fibroblasts (HDFs)(Ohnuki et al., 2014). Both clocks indicate age reversal following transduction with OSKM (**Figure 3c,d**). Universal Clock 2 showed decreased epigenetic age in partially reprogrammed cells after 11 days (**Figure 3c**).

### Transgenic mice for studying the somatotropic axis

Growth hormone, produced by somatotropic cells, stimulates the growth of all tissues of the body including bone. The somatotropic axis (growth hormone, IGF-1 levels, and their cognate receptors) occupies a central position in the study of aging and longevity^27^. Decreased growth hormone/IGF-1 signaling has been shown to extend longevity in a wide variety of species including mice^28^. A full body growth hormone receptor knockout (GHRKO) mouse currently holds the Methuselah Mouse Prize as the world’s longest-lived laboratory mouse, a title awarded by the Methuselah Foundation (http://reason.com/archives/2004/08/18/methuselah-mouse). To address the question whether lowered activity of the GH/IGF-1 pathway slows the universal pan-mammalian clocks, we analyzed 3 different mouse models: 1) Snell dwarf mice, which do not produce growth hormone, and as a consequence exhibit an increase lifespan^29,30^, 2) full-body growth hormone receptor knockout mice (GHRKO) which also exhibit an increase in lifespan ^31^, and 3) and liver-specific GHRKO mice which exhibit lowered serum IGF1-levels but not an increase in lifespan. Analyses with both Clocks 2 and 3 revealed that Snell dwarf mice do indeed exhibit noticeably slower epigenetic aging in several tissues (t-tests: cortex *P*=2.0×10^-8^, kidney *P*=6.0×10^-10^, liver *P*=1.0×10^-7^, tail *P*=1.0×10^-6^ and with less extent effects in blood *P*=2.0×10^-3^ and in spleen *P*=0.03, heatmap listed in **Figure 3e** and bar plots listed **in Figure S6**), compared to wild-type mice. The other long-lived mutant mice, full-body GHRKO mice, also exhibited considerably slower epigenetic aging in multiple tissues (liver *P*=3.0×10^-5^, kidney *P*=2.0×10^-5^, and cortex *P*=0.02, the second column in **Figure 3e** and **Figure S7**), compared to wild-type mice. Since growth hormone receptor signaling stimulates IGF-1 synthesis in the liver, it is a plausible hypothesis that the epigenetic age reversal in the dwarf mice is due to lower circulating levels of IGF-1. This hypothesis, however, is not supported by our epigenetic age measurements of liver-specific GHRKO mice which exhibit non-significant difference from the wild-type controls ^32^. Both Clocks 2 and 3 show that the liver-specific GHRKO mice are not epigenetically younger than the wild type mice (with the possible exception of liver or kidney exhibiting trend significance, the third column in **Figure 3e** and **Figure S8**).

### Caloric restriction in mice

Caloric restriction, which also slows the somatotrophic axis, is associated with prolonged lifespan in several mouse strains ^33,34^. Previous studies using mouse clocks have shown that CR reduces the rate of epigenetic aging in liver samples ^5,6,35^. Similarly, we find with these clocks, that CR results in a reduced rate of epigenetic aging in mouse liver samples (*P*=6.0×10^-12^ for Clock 2, *P*=7.0×10^-15^ for Clock 3, the last column **Figure 3e and Figure S9**).

### TET enzyme KO studies in mice

TET enzymes are instrumental in active DNA de-methylation. Since hydroxymethylation mediated by TET enzymes is prevalent in brain tissue, we applied the universal clocks to brain tissue samples from Tet1, Tet2, and Tet3 knockout (KO) mice. Analysis with our universal clocks revealed that Tet3 KO mice exhibit a reduced rate of epigenetic aging: cerebral cortex *P*=3.0×10^-9^ and striatum *P*=2.0×10^-12^ (the sixth column in **Figure 3e**). By contrast, significant epigenetic age reversal effects in brain tissue were relatively weak for Tet1 (cerebral cortex *P*=6.0×10^-3^ and striatum *P*=2.0×10^-4^, the fourth columns in **Figure 3e**) and barely could be observed forTet2 KO mice (*P*> 0.6, the fifth columns in **Figure 3e**). Bar plots with individual observations overlaid are depicted in **Figure S10**.

### Meta EWAS of age across species

As these universal clocks are established by penalized regression models, only CpGs with age-related methylation changes that are most predictive of age collectively, are selected to constitute the clock. This means that most other age-related CpGs are not retained in the final formulas, which are constructed based on minimizing squared loss with an elastic net penalty. To identify all age-related CpGs, we carried out two-stage meta-analysis across species and tissues in eutherians (98% of the samples) in epigenome-wide association study (EWAS). Cytosines that become increasingly methylated with age (i.e. positively correlated) were found to be more highly conserved across tissues and species (**Figure 4a**). From these, we identified 832 age-related CpGs at a significance threshold of α=10^-200^ across all eutherian species and tissues (**Figure 4a, Data S6.1**). Cytosines cg12841266 (*P*=1.4×10^-1001^) and cg11084334 (*P*=2.6×10^-891^), both located in exon 2 of LHFPL4 (Hg38), were the most significantly associated with age across all species, having a correlation ≥0.8 in 28 species (**Data S7**, three examples shown in **Figure 4b–d**). Another strongly age-correlated cytosine, cg09710440, resides in exon 1 of *LHFPL3* (P=5.0×10^-787^), a paralog of *LHFPL4* (**Figure 4a, Figure S11, Data S6.1–6.7**).

**Figure 4.**
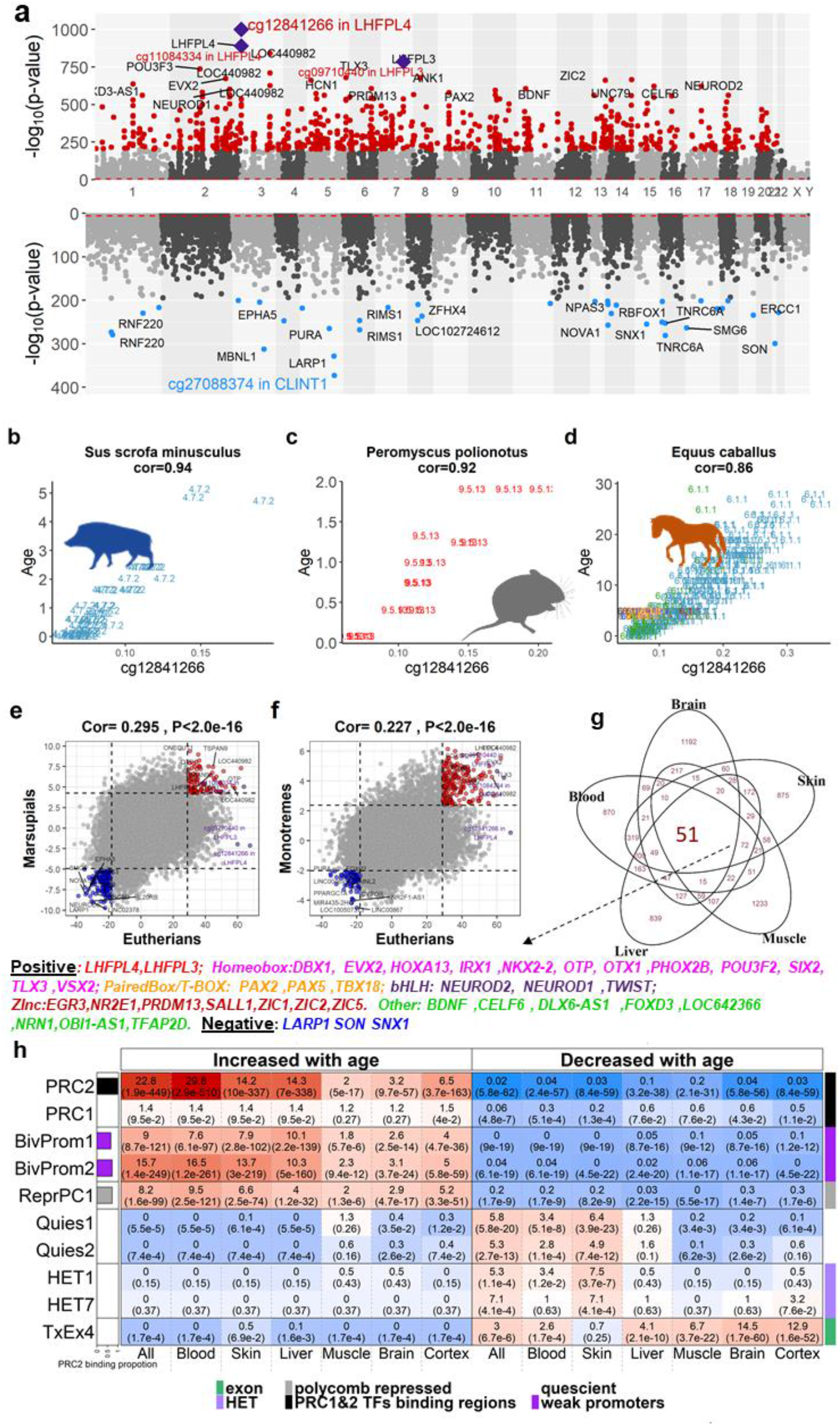
Meta-analysis of methylation change in function of chronological age across species and tissues. a–d, g–h correspond to EWAS of age in Eutherians. **a,**Meta-analysis P-value (−log base 10 transformed) of specific age-related CpGs (annotated with name of proximal genes) located on chromosomes (x-axis) based on human genome assembly 38 (hg38). The upper and lower panels of the Manhattan plot depict CpGs that respective gain or loss methylation with age. CpGs colored in red and blue exhibit highly significant (*P*<10^-200^) positive and negative age correlations respectively. The most significant CpG (cg12841266, *P*=1.41×10^-1001^) is located in exon 2 of the *LHFPL4* gene in humans and most other mammalian species, followed by cg11084334 (*P*=2.59×10^-891^). These two CpGs and cg097720 (*P*=4.97×10^-787^) located in the paralog gene *LHFPL3* are marked with purple diamonds. Scatter plots of cg12841266 (in *x*-axis) versus chronological age (years) in **b**, mini pigs (*Sus scrofa minusculus*), **c**, Oldfield mouse (*Peromyscus polionotus*) and **d**, horses (*Equus caballus*). Tissue samples are labeled by the mammalian species index and colored by tissue type as detailed in **Data S1.1—1.4**. **e**, correlation analysis between Z scores of EWAS of age in eutherians versus marsupials, **f**, correlation analysis between Z scores of EWAS of age in eutherians versus monotremes. Panels **g—h**: annotations of the top 1,000 CpGs with increased or decreased methylation with age that were identified in EWAS meta-analysis across all species and tissues (results in panel **a**), brain, cortex, blood, liver, muscle and skin tissues. **g**, the Venn diagram displays the overlap of age-associated CpGs across different organs, based on EWAS of the top 1,000 CpGs with positive/negative correlation. **g**, A total of 38 genes (35 positive age correlation and three negative age correlation) are proximal to the 51 age-associated CpGs common across all organs in the Venn diagram. The 35 positive genes are marked by different colors according to their protein family: 2 in LHPLF family, 12 in HomeoBox, 3 in PairesBox/T-Box, 3 in bHLH, 7 in zinc finger, and 8 in others. **h**, Selected results from universal chromatin state and polycomb-group (PRC) proteins annotation analysis. Odds ratios (P-value) are presented at each cell. The color gradient is based on −log10 (hypergeometric P-value) times sign of odds ratio >1. The complete results are listed in **Figure S3**.

As *LHFPL4* and *LHFPL3* are in human chromosomes 3 and 7 respectively, their consistent age-related gain of methylation is not due to physical proximity. It implies instead their common involvement in the aging process, even if their activities as nucleators of GABA receptors do not immediately conjure an obvious mechanism at this point in time. Methylation of *LHFPL4* cg12841266 was moderately to highly correlated with age of multiple mouse tissues in both developmental (r=0.58 and *P*=8.9×10^-11^) and post-developmental stages (r=0.5 and *P*=8.6×10^-122^), particularly in the brain (r=0.92 and *P*=6.95×10^-8^), muscle (r=0.89 and *P*=7.6×10^-7^), liver (r=0.81 and *P*=6.6×10^-132^), and blood (r=0.81 and *P*=9.8×10^-82^, **Figure S12**). To obtain a broad overview for the relationship of age association across different temporal domains, we repeated our two-stage meta-EWAS for young, middle and old age groups, respectively. Importantly, age-related methylation changes in young animals concur strongly with those observed in middle-aged or old animals, excluding the likelihood that the changes are those involved purely in the process of organismal development (**Figure 5a-c**). This point is further supported by visualizing the mean methylation levels (*β*) of age-related CpGs with respect to their distances from transcriptional start sites (TSS, **Figure 5d**). The remarkable similarity of the distribution patterns between all three age groups further demonstrates that these methylation changes are not due to organismal development. Other notable gene pairs that are proximal to the top 30 most significant age-related CpGs are *ZIC1* (human chromosome 3) and *ZIC2* (chromosome 13), *PAX2* (chromosome 10) and *PAX5* (chromosome 9) and *CELF6* (chromosome 15) and *CELF4* (chromosome 18, **Data S6.1**). As these gene pairs are all on different chromosomes, their concerted methylation change with age cannot be a result of mutual proximity, but more likely reflects their functional involvement in the aging process. Interestingly, each gene pair encodes proteins with similar activities, all of which are involved in the process of development.

**Figure 5.**
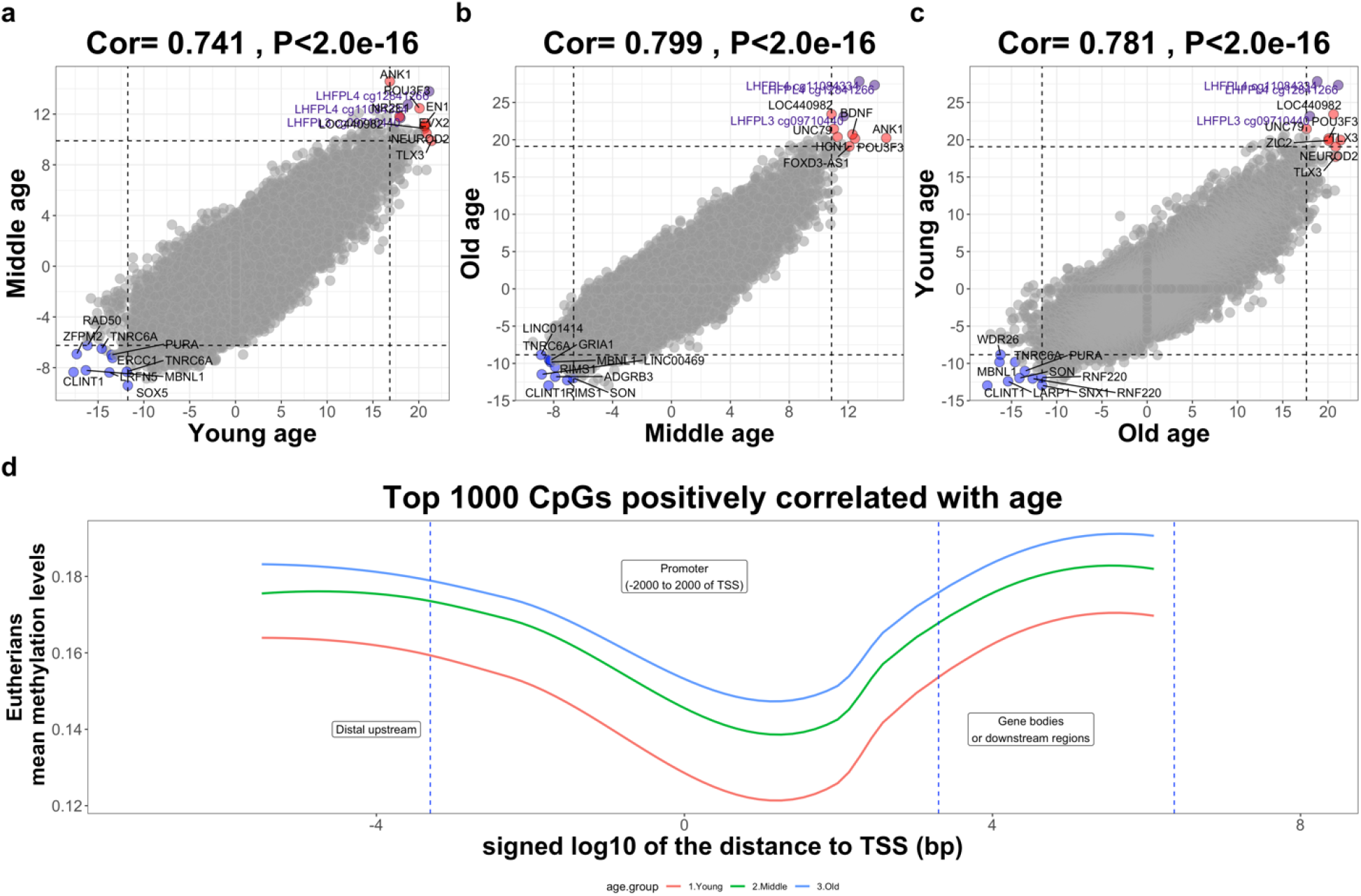
EWAS of age in three different age groups. Epigenome-wide association of age in three different age groups defined with respect to a species’ characteristics: average species age at sexual maturity (ASM) obtained from the anAge database (de Magalhaes et al., 2007). We defined the three age groups using intervals defined by multiples of ASM: young age is defined as age < 1.5* ASM, middle age is defined as age between 1.5 and 3.5 ASM, and old age is defined by age ≥3.5 ASM. Each axis reports a Z score from the meta-analysis EWAS of age across all mammalian species and tissues (Methods). Each dot corresponds to a CpG. Labels are provided for the top 10 hypermethylated/hypomethylated CpGs according to the product of Z scores in x and y-axis. CpGs that are located in LHFPL4/LHFPL3 genes are colored in purple. The Pearson correlation coefficient and corresponding nominal (unadjusted) two-sided correlation test P-value can be found in the title. **a**, EWAS of age in young versus EWAS in middle aged animals. **b**, EWAS of age in middle aged versus EWAS in old animals. **c**, EWAS of age in young versus EWAS in old animals. The high pairwise correlations indicate that conserved aging effects in mammals are largely preserved in different age groups. Many of the top CpGs for conserved aging effects in young mammals remain the top CpGs for conserved aging effects in old mammals. Specifically, we analyzed the mean methylation levels in eutherians across the three age groups. **d**,Mean methylation (y-axis) across the top 1000 CpGs positively correlated with age according to the EWAS across all mammalian tissue types (**Figure 1a**). The x-axis denotes the distance to the closest transcription start site (TSS) in log10 scale of bp. The positive TSS indicates the direction from 5’ to 3’ and the negative TSS indicates from the direction 3’ to 5’. The vertical phase is categorized into three regions: distal upstream ->promotor->gene bodies. The mean methylation levels are bounded by 0.2, reflecting that the CpGs beginning with lower methylation levels have higher propensity to increase with age.

### EWAS of age in marsupials and monotremes

We extended EWAS of age analysis to marsupials and monotremes. The top age-related CpGs for marsupial were also found to be proximal to genes involved in development. These include *GRIK2* (*P*=*8.8×10^-21^*, **Data S6.8**), which encodes a glutamate receptor associated with neurotransmitter activity and *ZIC4* (*P*=*2.7×10^-19^*), which encodes a Zinc finger protein. For EWAS of monotreme, the most significant CpG, cg22777952 (P=8.1×10^-10^, **Data S6.9**), is located in *FOXB1* (chromosome 5), which encodes a forkhead box protein. We only observed a moderate positive correlation with eutherian age-related methylation changes (r=0.295 in marsupials **Figure 4e** and r=0.227 in monotremes **Figure 4f**) which may reflect the relatively low number of samples from these groups. Despite these limitations, age effect on methylation of cg11084334 (but not cg12841266) in *LHFPL4* is nevertheless preserved in marsupials (P=4.8×10^-7^ in **Figure 4e**) and in monotremes (P=2.4×10^-5^ in **Figure 4f**).

### Meta-analysis of age-related CpGs across specific tissues

To gain a wider and deeper understanding of age-related CpGs within specific tissues across different species, we focused on 6 tissues/organs for which many species were available: brain (whole and cortex), blood, liver, skeletal muscle and skin. We performed an EWAS meta-analysis on 935 whole brains (18 species-brain tissue categories across 8 species), 391 cortices (6 species), 4,513 blood (56species), 1,063 liver (10 species), 354 muscle (5 species), and 2,363 skin (65 species, **Data S1.6—1.11**). Consistently across all tissues, there were more CpGs with positive correlations with age than negative ones (**Figure S11**) and most of them were located within CpG islands, which are known to become increasingly methylated with age (**Data S6.2–6.7**). While many of these cytosines were either specific to individual organs or shared between several organs, 51 potential universal age-related CpGs (48 positively and 3 negatively age-related CpGs) were shared among all five organs **(Figure 4g, Table S1)**. In total, 35 genes are proximal to the 48 CpGs and 3 genes are proximal to the 3 negative age-related CpGs. Interestingly, 20 of the 35 genes encode transcription factors, of which 11 encode homeobox (homolog) proteins, 7 encode zinc finger transcription factors and 2 encode paired box proteins, which are involved in developmental processes including embryonic development (**Table S1**). The relevance of this becomes evident below, where the chromatin state, function, and tissue-specific accessibility associated with the location of age-related CpGs are described.

### Analyses of chromatin states of DNA bearing age-related cytosines

To gain an overall perspective on the chromatin state in which age-related CpGs are located, we employed a detailed universal chromatin state map in which chromatin structure and their associated characteristics are annotated. This recently-available resource was constructed based on 1,032 experiments that mapped 32 types of chromatin marks in over 100 human cell and tissue types ^36^ (**Figure 4h, Figure S13, Data S8.2–8.9**). We overlaid the positions of the top 1,000 age-related CpGs onto this universal chromatin state map, and observed that all *positive* age-related CpGs showed strong enrichments in states that were previously shown to have a strong association with PRC2-binding sites (states BivProm1-2, ReprPC1)^36^. These CpGs localized to PRC2-binding sites, which are characterized by Eed, Ezh2, and Suz12 binding, as shown in the first row of **Figure 4h**. This was the case when all tissue types were analyzed collectively (odds ratio OR=22.8, hypergeometric *P*=1.9×10^-449^), and remained so when tissues were analyzed individually: blood (OR=29.8, *P*=2.9×10^-510^), liver (OR=14.3, *P*=7.3×10^-338^), skin (OR=14.3, *P*=7.0×10^-338^), cortex (OR=6.5, *P*=3.7×10^-163^), and brain (OR=3.2, *P*=9.7×10^-57^). Indeed, the vast majority of positive age-related CpGs were highly enriched in PRC2-binding sites: 808 of these PRC2-related CpGs correlated significantly with age in blood, 675 in liver and 672 in skin (**Data S8.1**).

In contrast, there was no significant enrichment of age-related cytosine in polycomb repressive complex 1 (PRC1)-binding sites in any of the tissue types (second row of **Figure 4h**). This accentuates the high specificity of enrichment of age-related CpG to PRC2-binding sites. PRC2 is a transcriptional repressor complex best known as a writer of H3K27 methylation, a chromatin modification associated with transcriptional repression^37^. Significantly, PRC2-mediated methylation of H3K27 is essential for the establishment of bivalent promoters, which simultaneously contain both H3K27me3 and H3K4me3 histones. As such, it is consistent that positive age-related CpGs are also found to be enriched in bivalent promoter states (rows 3 and 4 of **Figure 4h**). More specifically, there is greater enrichment of these CpGs in a bivalent promoter state that contains more H3K27me3 than H3K4me3 (state BivProm2), compared to states such as BivProm1, which contains equal amounts of these two histone types. The combination of these features are epitomized by the top EWAS hit *LHFPL4* cg12841266, which resides in a bivalent chromatin state (BivProm2), while also being located in a PRC2-binding region (Eed, Ezh2, and Suz12-binding sites (**Data S8.1**). These mammalian results echo those from *human* studies^38,39^, where tissue-independent age-related gain of methylation is characterized by cytosines that are located in PRC2-binding sites and bivalent chromatin domains.

Negative age-related CpGs (those that lose methylation levels with age) on the other hand, led to less significant overlaps with other chromatin states. When it comes to age-related loss of methylation it is important to distinguish proliferative tissues from non-proliferative tissue such as brain and muscle.

In highly proliferative tissues such as blood and skin, age-related loss of methylation is seen in CpGs located in genomic regions Quies1 and Quies2, which are characteristic of quiescent cells, as well as HET7 and HET1, which are heterochromatin regions. Both Quies and HET regions are marked by H3K9me3 (see **Data S8.2** and Vu&Ernst ^36^. Age-related loss of methylation in brain and muscle on the other hand, is observed in CpGs located in exon-associated transcription state TxEx4 (OR=14.5, P=1.7×10^-60^ in brain and OR=6.7, P=3.7×10^-22^ in muscle). In contrast, TxEx4 is not strongly enriched with age-related cytosines that lose methylation in blood (OR=2.6, P=1.7×10-4) or skin (OR=0.7, P=0.25). TxEx4 is most highly enriched for transcription termination sites, exons and promoters ^36^.

### Overlap with late-replicating domains

Our chromatin state analysis of age-related loss of methylation demonstrated that it is important to distinguish proliferating tissues (blood, skin) from non-proliferative tissues (brain, muscle). Therefore, we proceeded to look carefully at the relationship between DNA replication and methylation. Late-replicating domains of the genome have the propensity to become partially methylated: replication-related loss of methylation is particularly pronounced in cytosines known as solo-WCGW, which are CpGs flanked by A or T on either side ^40^. We overlayed the top 1,000 age-related CpGs (positive/negative) on the reported late-replicating domains, which are known also as partially methylated domains (PMDs)^40^. As previously reported for human tissues ^40^, we observed age-related loss of methylation in PMDs and solo-WCGW in mammalian tissues that proliferate, such as blood and skin (**Figure S14** and **Data S9**). Indeed, the top 1,000 negatively age-related CpGs in mammals exhibit highly significant overlap with common PMDs and solo-WCGW (Hg19): skin (OR=7.9, *P*=1.6×10^-90^), blood (OR=5.3, *P*=1.5×10^-50^), and all (OR=7.3, *P*=4.4×10^-81^, **Figure S14**). Importantly, this is not the case in non-proliferating tissues such as the brain where a different pattern emerges: CpGs that lose methylation with age are enriched in highly methylated domains (HMDs) as opposed to PMDs (OR=3.3, P-value=1.9×10^-74^). In contrast, CpGs that gain methylation with age exhibit substantially weaker overlap with both PMDs and HMDs. We made similar observations with late-replicating domains in the mouse genome (mm10, **Figure S14**). Overall, these results demonstrate that pan-mammalian CpGs that lose methylation with age are enriched in late-replicating regions of highly proliferative tissue.

### Functional enrichment analysis of age-related CpGs

We employed the Genomic Region Enrichment of Annotation Tool (GREAT) to annotate the function of *cis*-regulatory regions of age-related CpGs^41^. This will help to ascertain the biological processes and pathways that are potentially associated with the top 1,000 positive and the top 1000 negative age-related CpGs (**Figure 6**, **Data S10.1–10.17**). Analysis of positively-correlated CpGs across all tissues, revealed “nervous system development” to be a highly significant (*P*=1.3×10^-203^) Gene Ontology (GO) term. The same term could be found when individual tissues were analyzed specifically in blood (*P*=1.9×10^-224^), liver (*P*=2.6×10^-137^), muscle (*P*=3.4×10^-14^), skin (*P*=1.7×10^-145^), brain (*P*=6.4×10^-35^) and cortex (*P*=1.0×10^-78^). Other highly significant GO terms include “developmental process”, “regulation of RNA metabolic process”, “nucleic acid binding transcription factor (TF) activity”, “pattern specification” and “anatomical structure development” (**Figure 6**). Consistent with the above observation with the universal chromatin state map, GREAT analysis also revealed that a high proportion of the top 1,000 positive age-related CpGs reside in PRC2 target sites (GREAT *P*=8.3×10^-212^). This was also the case when individual core PRC2 subunits (SUZ12, EED, or EXZH2) were analyzed (**Figure 6**). It follows that these CpGs are also enriched in promoters with the H3K27me3 modification in embryonic stem cells: all tissue (*P*=1.9×10^-269^), blood (*P*=2.8×10^-285^), liver (*P*=3.4×10^-182^), muscle (*P*=9.0×10^-17^), skin (*P*=7.8×10^-202^), brain (*P*=1.4×10^-54^), and cortex (*P*=2.7×10^-115^) (**Figure 6**).

**Figure 6.**
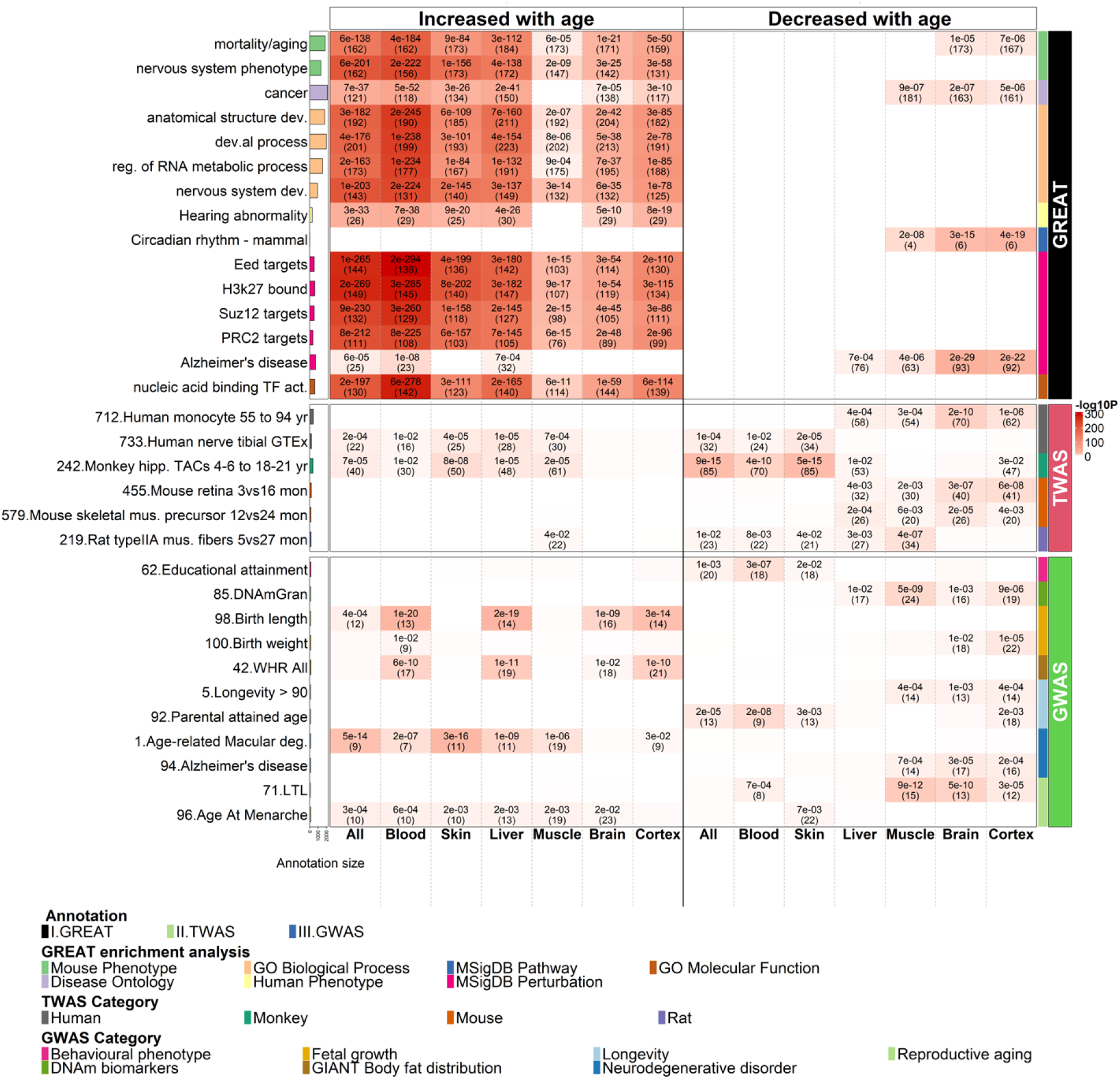
Biological pathways and functional gene sets enriched in age-related CpGs. Selected results from (I) genomic region-based GREAT functional enrichment (top panel), (II), gene-based EWAS-TWAS enrichment analysis (middle panel), and (III) genomic region-based EWAS-GWAS enrichment analysis (bottom panel). All the enrichment analyses were based on hypergeometric tests with background based on mammalian array. The bar plots in the first column report the total number of genes at each studied gene set adjusted based on the background. The left and right panels of the x-axis list the top 1,000 CpGs that increased or decreased with age from meta-EWAS of age across all, blood, skin, liver, muscle, brain and cerebral cortex tissues, respectively, respectively. On the right-side, the first column color band depicts the three types of enrichment analyses. The second column color band depicts (I) a total of six ontologies in GREAT analysis, (II) four species in our TWAS collections, and (III) a total of seven categories of human complex traits in GWAS as described in the legend. The heatmap color codes −log10 (hypergeometric P-value). Unadjusted hypergeometric P-values (number of overlapped genes) are reported in the heatmap provided (I) false discovery rate (FDR) <0.05, P<0.001 and the number of overlapped genes ≥3, (II) P<0.05 and (III) P<0.05. Abbreviations: act.=activity, deg.=degeneration, dev.=development(al), LTL= leukocyte telomere length, mus.=muscle, OPCs= oligodendrocyte precursor cells, TACs= transiently amplifying progenitor cells, WHR= waist to hip ratio.

These results corroborate our findings in the chromatin annotation analysis above which highlights the strong connection between PRC2 target sites and age-related gain of methylation. As PRC2 plays a critical role in development, these results reinforce the epigenetic link between development and aging. This link finds support in observations that mice that are developmentally compromised following ablation of growth hormone receptors (GHRKO) or the anterior pituitary gland (Snell mice) exhibit reduced rates of epigenetic aging in multiple tissues as measured by the universal epigenetic age clocks (**Figure 3e**).

While positive age-related CpGs (across all tissues) were enriched in 2,961 gene ontology/MSigDB terms at a false discovery rate of 0.05 (**Data S10.1**), negative age-related CpGs were enriched in only three. The dearth of significant enrichment for negative age-related CpGs is particularly pronounced for skin, blood, and liver for which a large number of species were available (**Data S10.9-12, 15 and 16)**. Interestingly, negative age-related CpGs in brain and muscle are enriched in genes that play a role in circadian rhythm (brain *P*=3.3×10^-15^, cerebral cortex *P*=4.0×10^-19^ and muscle *P*=2.3×10^-8^, **Figure 6**). They also significantly overlapped with Alzheimer’s disease-related gene sets (e.g., *P*=1.8×10^-29^ in brain and *P*=2.4×10^-22^ in cerebral cortex in **Figure 6**), and with gene sets related to mitochondrial function in brain, cerebral cortex, and muscle (e.g. *P*=3.6×10^-7^, **Data S10.2**).

Lastly, GREAT analysis showed that both positive and negative age-related CpGs were enriched with the gene sets related to mortality/aging, cancer (**Figure 6**) and targets of three Yamanaka factors: SOX2, MYC, and OCT4 (**Data S10.3**). Of the 341 genes proximal to positively age-related CpGs in all tissues, 162 are implicated in mortality/aging (*P*=6.3×10^-138^, **Figure 6**). Similar enrichments in mortality/aging genes are also seen in specific tissues: blood (*P*=3.8×10^-184^), liver (*P*=2.7×10^-112^), muscle (*P*=6.2×10^-5^), skin (*P*=9.1×10^-84^), brain tissues combined (*P*=1.2×10^-21^), and cerebral cortex (*P*=5.0×10^-50^). Enrichment for mortality/aging-related genes are also found in the top 1,000 negatively age-related CpGs in the brain (*P*=1.0×10^-5^) and cerebral cortex (*P*=7.5×10^-6^). Concerns that these highly significant enrichments may be a result of potential biases in the mammalian methylation array platform could be discounted after sensitivity analysis, as reported in **Supplementary Information, note 3**.

### Transcription factor binding

We used the CellBase ^42^ and ENCODE databases^43^ to annotate CpGs with binding sites identified through ChIP-seq, of a total of 68 TFs. The ChIP-seq analysis of these TFs was carried out in 17 different cell types depending on the target molecule. We assigned a CpG with a specific TF if it overlapped with the binding site of the latter in at least one cell type (in Hg19 assembly). The transcription factor enrichment analyses of the most significant age-related CpGs in mammals led to highly significant results for positively age-related CpGs. Among the top 1000 positively age-correlated CpGs across mammalian species, *REST* TF was found to be the most significant in all tissues (*P=*3.1×10^-54^), in blood only (*P*=2.7×10^-32^), skin (*P*=6.8×10^-59^), liver (*P*=1.5×10^-28^) and weakly in muscle (*P*=2.2×10^-3^), cortex (*P*=0.01) and brain (*P*=0.09, **Figure S15, Data S11**).The ChIP-seq analysis of the REST TF was done in five cell lines including human embryonic stem cell line (**Data S11.1**). Incidentally, the CpG cg12841266 on *LHFPL4* is also located in the binding regions of *REST. REST* is known for repressing neuronal genes in non-neuronal tissues, which could explain the weak enrichments in brain regions. Other significant TF enrichments include *TCF12* and *HDAC2*. Gene *TCF12*, involved in the initiation of neuronal differentiation, encodes a protein in the bHLH E-protein family. The binding regions overlap several of our top genes including another bHLH gene *NEUROD1* (**Table S1, Data S11**). We also found moderate enrichments with *CTCF, NANOG* and others. Among the top 1000 negatively age-related CpGs, we observed far fewer significant TF binding enrichment, of which JUN (c-jun) on chromosome 1 in blood (*P*=2.6×10^-9^) and brain (P=0.024, **Figure S15**) are notable.

### Age-related CpGs and age-related transcriptomic changes

We evaluated whether the top 1,000 positive and negative age-related CpGs are adjacent to genes whose mRNA levels correlate with age. We evaluated tissue-specific transcriptome-wide association studies (TWAS) of age in four mammalian species (human, monkey, mouse and rat) curated by two data bases: GenAge^44^ and Enrichr^45,46^. Our EWAS-TWAS overlap analysis (**Figure 6, Figure S16** and **Data S12**) reveals significant overlaps between age-related CpGs and age-related transcriptomic changes in several species, including human tibial nerve samples (collected by GTEx, **Figure 6**), normal monkey hippocampal samples (*P*≥=5.0×10^-15^) and several rat and mouse tissues (**Figure 6**). Overall, however, the overlap between EWAS and TWAS of age is weak and tissue-specific. This may reflect weak conservation of TWAS across tissue types and species.

### Age-related CpGs and GWAS of human traits

The proximal genes of the same top 1,000 positively and negatively age-related CpGs were compared with the top 2.5% of genes implicated by different human genome-wide association studies (GWAS) (**Supplementary Information, note 4**). The most significant enrichment was observed for positive age-related CpGs in genes associated with human length at birth (blood; *P*=1.0×10^-20^ and liver; *P*=2.0×10^-19^). Significant enrichments (defined here as nominal *P*<5.0×10^-4^) were also seen with genes linked to parental longevity (parental attained age; *P*=2.0×10^-8^ of negatively age-related CpGs in blood, **Figure 6, Figure S17, Data S13.1–13.7**), human longevity (*P*=4.0×10^-4^ with CpGs that lose methylation in cerebral cortex), Alzheimer’s disease (*P*=3.0×10^-5^ with negatively age-related CpGs in brain samples), leukocyte telomere length (*P*=9.0×10^-12^ and *P*=5×10^-10^ for negatively age-related CpGs in muscle and brain respectively), waist-to-hip ratio (*P*=6.0×10^-10^ and *P*=1.0×10^-11^ with positively age-related CpGs in blood and liver, respectively), and age at menarche (*P*=3.0×10^-4^ with positively age-related CpGs in all tissues). Overall, the overlap analysis with GWAS results demonstrates that pan-mammalian age-related CpGs are located near genes that play a role in human development (length at birth, menarche), obesity, and human longevity.

### Single cell ATAC seq analysis of human bone marrow

Low-methylated regions that are away from transcriptional start sites are associated with open chromatin, transcription factor-binding and enhancers ^43^. Therefore, our top positively age-related pan mammalian CpG sites (which begin as lowly methylated sites and gain methylation with age) could indicate a gradual loss of these open chromatin regions. To test this hypothesis, we investigated the association between the top 35 positively age-related CpGs (**Table S1**) and chromatin accessibility in single cells isolated from human bone marrow mononuclear cells (BMNCs). These single cell ATAC data are from a recent study ^47^ that used 10x Multiome technology to profile both ATAC and gene expression within the same cell across 10 donors of varying age. We overlapped the genomic regions of the top 35 CpGs (**Table S1**) with the called ATAC peaks within the BMNC dataset, resulting in 17 genes including *LHFPL4* **(Data S14, Figure 7a)**. Next, we calculated the percentage of cells in each individual containing the respective peak. Strikingly, we observed a strong and statistically significant negative correlation (**Figure 7b**) between age and the number of cells with the called ATAC peak overlapping cg12841266 on *LHFPL4,* confirming that with age (as this region gains methylation), the number of cells with open chromatin is reduced. Out of the 17 gene regions 16 showed a negative correlation with age, of which 7 were statistically significant (*P* < 0.05, **Figure 7a**) and this loss of accessibility with age was highly enriched for these hypermethylated sites (*P* < 0.001 based on permutation tests, **Figure 7b**). The seven significant genes (*LHFPL4, TLX3, ZIC2, PAX2, NR2E1, NEUROD1 and DLX6-AS1*) play a role in developmental processes (**Table S1**). As a side note, the other zinc finger gene, *ZIC5,* also exhibits a marginally significant negative correlation with age (r= −0.54 and *P*=0.07, **Data S14**). We did not observe any scATAC-seq signal in the region of cg09710440 on *LHFPL3,* plausibly owing to its close proximity to the transcriptional start site (232 bp).

**Figure 7.**
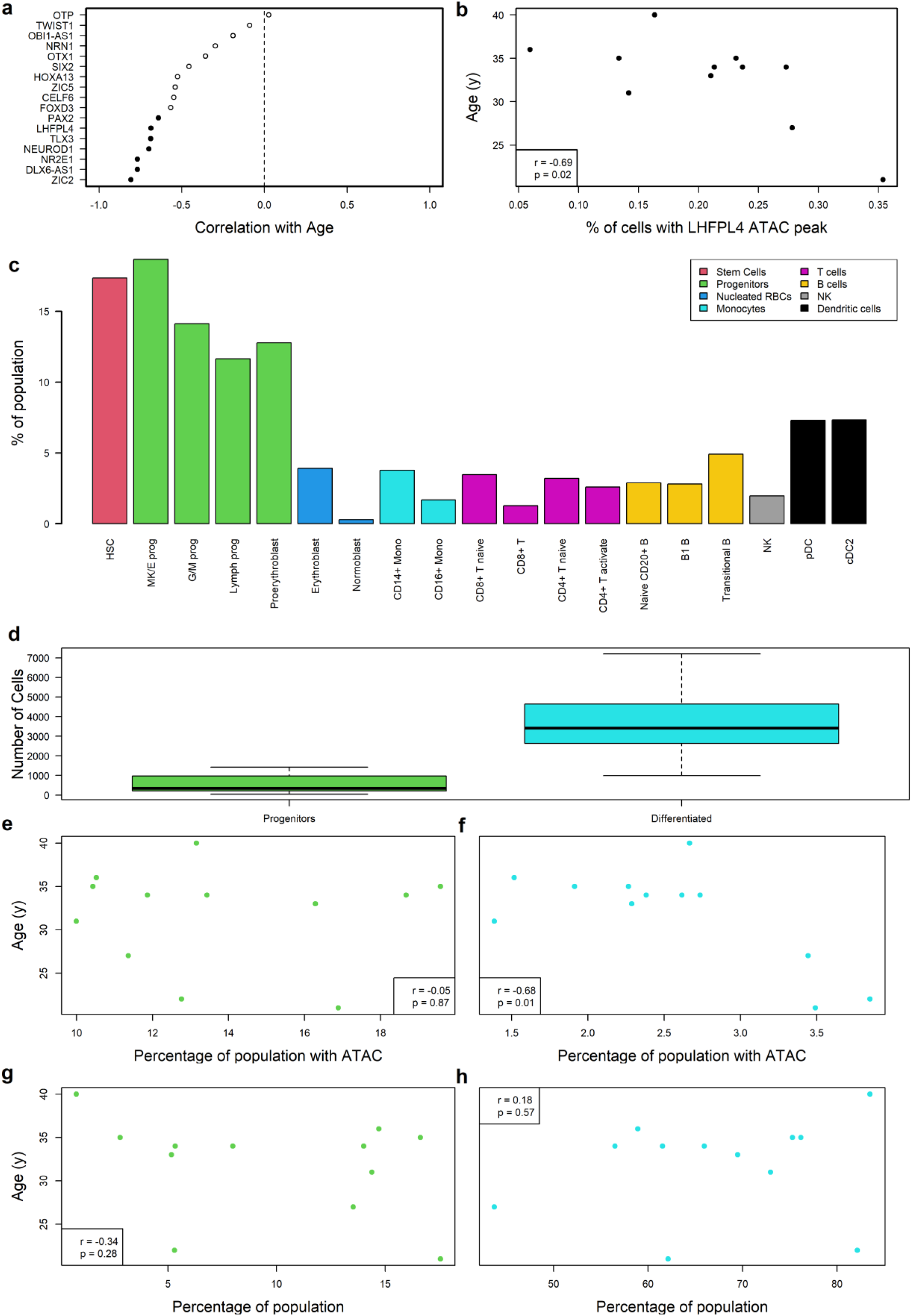
Single cell analysis of age-related changes in bone marrow. **a**, scATAC-seq results of 17 out of the 35 genes (listed in Table S1) that show at least a called ATAC peak in the region overlapping with our top CpGs with positive age correlation (listed in Table S1). The y-axis lists gene symbol. The x-axis reports the Pearson correlation between chronological age and the percentage of cells presenting an scATAC-seq signal in the respective CpG site (labelled by adjacent gene). The genes are ordered by correlation estimate (from most negative). A negative correlation estimate indicates the accessibility of the CpG site decreases with chronological age. Each dot presents a gene and marked by a solid shape provided P<0.05. **b**, scATAC-seq analysis results for *LHFPL4*. The y-axis depicts chronological age and the x-axis, the percentage of cells with an scATAC-seq signal (x-axis). **c**, The barplot presents the proportion of counts for each cell population that have at least one detected scATAC-seq signal among the seven genes with significant age correlation in panel a, arranged by stem-cell and progenitor populations on the left and differentiated cells on the right. **d**, number of cells identified as either a progenitor cell (left side includes hematopoietic stem cell [HSC] type) or a differentiated cell (right side). **e, f**, show the percentage of these two cell populations (e for progenitor and f for differentiated cell type) which contain at least one ATAC-seq signal, plotted against the age of each individual. **g, h** show the percentage of these two cell populations regardless of ATAC-seq signals (g for progenitor and h for differentiated cell type), plotted against the age of each individual.

We next tested whether these 7 statistically significant ATAC peaks marked a specific subset of cell types. Given the sparsity of single cell ATAC data, we calculated the proportion of each population containing at least one of these regions and found that stem cell/progenitor populations had a considerably higher proportion of open chromatin in these loci, compared to differentiated cells (mean 14.9% versus mean 2.9%, **Figure 7c**). Given that the open chromatin state of these regions is more enriched in progenitor cells, it is possible therefore, that the observed age-related reduction of these open chromatin states is due to the loss (e.g. death or differentiation) of progenitor cells in the tissue. To investigate this, we separated progenitor cells, including hematopoietic stem cells (HSC), from differentiated cells, and correlated the proportion of the cell type with age and the proportion containing an ATAC peak (**Figure 7d--h**). Interestingly, there was no loss of open chromatin (ATAC signal) with age in progenitor cells, although there was a negative correlation (r=-0.34, p=0.28) of the percentage of progenitors within each individual (**Figure 7e&g**). In contrast, we saw a significant loss of ATAC signal with age (**Figure 7f**), and no change in population of differentiated cells (**Figure 7h**). This suggests that these regions, which gain methylation and lose accessibility with age, are within differentiated cell population rather than the progenitor population. This rules out a passive loss of open chromatin state of these regions over time within the progenitor compartment and points instead to an age-related loss of an active process (epigenetic maintenance system) that maintains these states in differentiated cells.

## DISCUSSION

The universality of aging across all mammalian species has engendered speculations of its cause, with the predominant view that random damage to cells underlies this process. This idea, however, is challenged by the ability to accurately estimate ages of mammals by virtue of their DNA methylation profiles. Such precise alteration with age introduces the greater likelihood that aging may be driven, at least in part, by deterministic processes. The Mammalian Methylation Consortium investigated this question by generating an unprecedentedly large set of DNA methylation profiles from 348 species. Of those, 174 eutherian-, nine marsupial-, and two monotreme species with age information were used in this study. The results are clear. The methylation level of a set of CpGs found in some DNA sequences across mammals changes consistently with age, with the majority of them gaining methylation. This generality across mammalian species is accompanied by the impressive specificity with regards to the location of these age-related CpGs. Those that gain methylation are primarily in PRC2-binding sites and bivalent regulatory regions known as bivalent promoters 1 and 2. Both of these chromatin structures characterize regions that regulate expression of genes involved in the process of development ^37,48,49^, which is one of the most conserved biological processes that threads through all mammalian species. A typical example of such CpGs are those that are proximal to *LHFPL4* and *LHFPL3.* The currently understood activity of *LHFPL4* in synaptic clustering of GABA receptors does not immediately present an obvious connection to aging across all tissues, suggesting either that potentially greater functions of these proteins have yet to be realized or that the impact of clustering these receptors on the process of aging has yet to be understood. Despite this unknown, the specificity of their methylation change with age is not in doubt, given their locations on different human chromosomes, as is the case for several gene pairs including *LHFPL3/4, ZIC2/5, PAX2/5* and *CELF4/6.* The single cell ATAC-seq analysis of bone marrow mononuclear cells showed that CpGs with positive age correlations in most species are located in sites that lose chromatin accessibility with age in differentiated cells but not in progenitor cells. Such chromatin compaction is likely to be instigated by methylation of these sites ^50^, and their closure precludes the access of PRC2 complex to it targets. Hence methylation can prevent PRC2 from binding to its targets by virtue of structural incompatibility (methylated DNA sequence) and accessibility through chromatin compaction. In human bone marrow, we observed that (i) the chromatin states in the top age-related PRC2 targets are open in substantially more progenitor cells than differentiated cells, and (ii) with age, the percentage of progenitor cells with open age-related PRC2 targets remain constant, while the percentage of differentiated cells with open PRC2 targets decreased. Although this does not entirely rule out proliferation (or death) of specific cell populations within the differentiated compartment of the marrow, it does increase the likelihood that intracellular molecular alterations (age related closing of open chromatin in differentiated cells) are responsible for these age-related methylation changes.

When it comes to age-related loss of methylation it is important to distinguish proliferative tissues from non-proliferative tissues such as brain and muscle. In highly proliferative tissues, age-related loss of methylation is seen in CpGs located in genomic regions which are characteristic of quiescent cells, as well as heterochromatin regions. In proliferative tissues, negatively age related CpGs are often located in partially methylated domains (PMDs). Interestingly, PMDs are in late DNA replication regions. As methylation of replicated DNA is slow, and only completed very late in S and G2 phase, late replicated regions of the DNA are particularly disadvantaged in this regard. Indeed, progressive methylation loss in PMDs is exploited as mitotic clock, which also correlates very well with chronological age ^40^. As such, their identification as pan-mammalian negatively age-related CpGs is entirely consistent with studies observed in human cells. Interestingly, this late-replication effect on DNA methylation can be prevented by the binding of histone H3K36me3 to these regions^40^. This appears to be mediated by H3K36me3 recruitment of Dnmt3b to unmethylated and newly replicated DNA. Conversely, functional loss of NSD1, the enzyme that generates H3K36me3, leads to hypomethylation of DNA and accelerated epigenetic aging ^51,52^. Age-related loss of methylation in non-proliferative tissues (brain and muscle) on the other hand, is observed in CpGs located in an exon-associated transcription state (TxEx4), which is most highly enriched for transcription termination sites and is associated with the highest gene expression levels across many cell and tissue types ^36^.

Unlike CpGs that gain methylation with age, CpGs that lose methylation are typically not related to developmental genes. Instead, they are related to genes of circadian rhythm and mitochondria, the functions of which are progressively eroded with age. Finally, the *LARP1* gene, which is proximal to the highest ranked hypomethylated cytosine in liver and second across all tissues, encodes a protein that regulates translation of downstream targets of mTOR ^53^. mTOR has very well-documented links with aging and longevity ^54^, and is also linked to epigenetic aging ^55,56^. Overall, we provide collective evidence that methylation of the mammalian age-related CpGs that we identified are not merely stochastic marks accrued with age. They are instead, methylation changes that capture multiple facets of mammalian aging.

The deterministic features of these age-related changes on the mammalian epigenome make a compelling case that aging is not solely a consequence of random cellular damage accrued in time. It is instead a pseudo-programmed process that is also intimately associated with mammalian development that begins to unfold from conception. This is supported by, and consistent with, the finding that genes proximal to age-related CpGs were also identified by GWAS of human development features such as length at birth and age at menarche. A large body of literature including those by Williams in 1957 ^57,58^ has suggested a connection between growth/development and aging. More recently, several authors have suggested epigenetics to be the link between these two processes ^57,59–67^. This notion is further supported by recent demonstration of age reversal through transient expression of Yamanaka factors^68–73^, and the overlap between the targets of these factors and age-related CpGs that we identified here.

According to the pseudo-programmatic theory of aging, the process of aging is very much a consequence of the process of development, and the ticking of the epigenetic clock reflects the continuation of developmental processes ^57,67^. As predicted by the epigenetic clock theory of aging, the universal epigenetic clocks provide a continuous read-out of age from early development to old age in all mammals, as this feature underlies the continuous and largely deterministic process of aging from conception to tissue homeostasis ^61^. Consistent with this theory, the pan-mammalian methylation clocks are slowed by conditions that delay growth/development including Snell mice and fully body GHRKO mice. The successful construction of universal clocks is a compelling mathematical demonstration of the deterministic element in the process of aging that transcends species barriers within the mammalian class. Indeed, the centrality of PRC2, which is also present in non-mammalian classes, implies that the process of aging that is uncovered here is likely to be shared by vertebrates in general. Our human epidemiological studies and mouse interventional studies show that pan-mammalian clocks relate to human and mouse mortality risk, respectively. Our analysis of human reprogramming cells shows that the clocks can track different phases of reprogramming.

Overall, these results demonstrate that epigenetic aging is universal across all mammalian species and captures multiple processes and manifestations of age that have hitherto been thought to be unrelated to each other. The availability of these epigenetic clocks opens the path to uncovering the common age-related factors that link these apparently disparate processes and introduce a deep and comprehensive understanding of mammalian aging.

## METHODS

### Tissue samples

The tissue samples are described in the **Data S1.1-1.4** and related citations as listed in **Supplementary Information, notes 1&2**.

### Quality controls for establishing universal clocks

We generated two variables to guide the quality control (QC) of the study samples; the first being a variable indicating the confidence (0–100%) in the chronological age estimate of the sample. For example, a low confidence was assigned to samples from wild animals whose ages were estimated based on body length measurements. The epigenetic clocks were trained and evaluated on tissue samples from animals whose ages could be assessed with high confidence (less than 10 percent error). The second quality control variable was an indicator variable (yes/no) that flagged technical outliers or malignant (cancer) tissue. Since we were interested in “normal” aging patterns we excluded tissues from preclinical studies surrounding anti-aging or pro-aging interventions.

### Species characteristics

Species characteristics such as maximum lifespan (maximum observed age) and age at sexual maturity were obtained from an updated version of the Animal Aging and Longevity Database^74^ (AnAge, http://genomics.senescence.info/help.html#anage). To facilitate reproducibility, we have posted this modified/updated version of AnAge in **Data S1.13.**

### Three universal pan-mammalian clocks

We applied elastic net regression models to establish three universal mammalian clocks for estimating chronological age across all tissues (n=11,754 from 185 species) in eutherians (N=11439 from 176 species), marsupials (n=210 from 9 species) and monotremes (n=15 from 2 species). The three elastic net regression models corresponded to different outcome measures described in the following: 1) log transformed chronological age: *log*(*Age* + 2) where an offset of 2 years was added to avoid negative numbers in case of prenatal samples, 2) −*log*(−*log* (*RelativeAge*)) and 3) log-linear transformed age. DNAm age estimates of each clock were computed via the respective inverse transformation. Age transformations used for building universal clocks 2 and 3 incorporated three species characteristics: gestational time (*GestationT*), age at sexual maturity (*ASM*), and maximum lifespan (*MaxLifespan*). All of these species variables surrounding time are measured in units of years.

#### Loglog transformation of Relative Age for clock 2

Our measure of relative age leverages gestation time and maximum lifespan. We define relative age (*RelativeAge*) and apply the double logarithmic *Loglog* transformation:

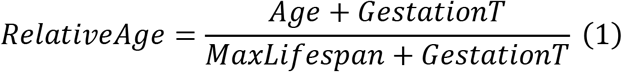

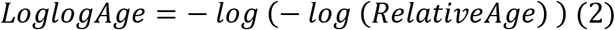

By definition, *RelativeAge* is between 0 to 1 and *LoglogAge* is positively correlated with age. The incorporation of gestation time is not essential. We simply include it to ensure that RelativeAge takes on positive values. We used the double logarithmic transformation to link relative age to the covariates (cytosines) for the following reasons. First, the transformation maps the unit interval to the real line. Second, this transformation ascribes more influence to exceptionally high and low age values (**Figure S1a-c**). Third, this transformation is widely used in the context of survival analysis. Fourth, this non-linear transformation worked better than the identity transformation.

The regression model underlying universal clock 2 predicts *LoglogAge*. To arrive at the DNA methylation based age estimate, one needs to apply the inverse transformation to *LoglogAge* based on the double exponential transformation:

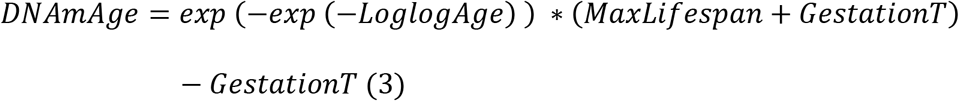

All species characteristics (e.g. MaxLifespan, gestational time) come from our updated version of AnAge. We were concerned that the uneven evidence surrounding the maximum age of different species could bias our analysis. While billions of people have been evaluated for estimating the maximum age of humans (122.5 years) or mice (4 years), the same cannot be said for any other species. To address this concern, we made the following assumption: the true maximum age is 30% higher than that reported in AnAge for all species except for humans and mice (Mus musculus). Therefore, we multiplied the reported maximum lifespan of non-human or non-mouse species by 1.3. Our predictive models turn out to be highly robust with respect to this assumption.

#### Transformation based on log-linear age for clock 3

Our measure of log-linear age leverages age at sexual maturity (ASM). The transformation has the following properties: it takes the logarithmic form when the chronological age is young and takes the linear form otherwise. It is continuously differentiable at the change.

First, we define a ratio of the age relative to ASM, termed as RelativeAdultAge, as the following:

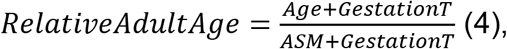

where the addition of *GestationT* ensures that the *RelativeAdultAge* is always positive. To model a faster rate of change during development, we used a log-linear transformation on *RelativeAdultAge* based on a function that generalizes the original transformation proposed for the human pan-tissue clock^4^:

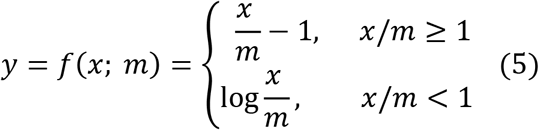

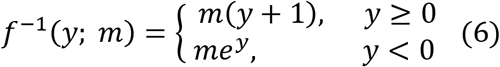

where x denotes RelativeAdultAge. This transformation ensures continuity and smoothness at the change point at *x* = *m*.

In the following, we describe the estimation of the parameter m. To ensure maximum value of *y* to be the same across all species, the parameter *m* should be proportional to the maximum of *x* for each species, i.e. the best value for m would be the oracle value 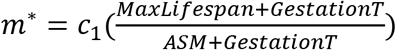 (**Figure S7d**).

The proportionality constant *c_1_* controls the distribution of *y*. We chose it so that *y* follows approximately a normal distribution with mean zero. Since we wanted to define Clock 3 without using *MaxLifespan*, we used the ratio 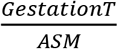 to approximate the oracle value *m*\* by fitting the following regression model with all the species available in our anAge database,

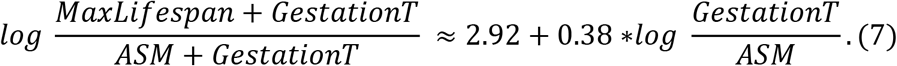

The two log variables in the formula 7 have moderate correlation (r=0.5). Subsequently, we defined 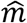 as follows

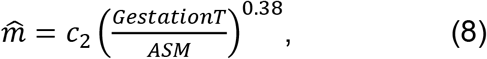

where *c*_2_ = *c*_1_*e*^2.92^. We chose *c*_2_ = 5.0, so that *LoglinarAge* (termed as y in equation (5)) follows approximately a normal distribution with mean 0 (median = 9.0×10^-4^, skewness = −0.02, **Figure S7f**).

Setting *x* = *RelativeAdultAge* in equation (5) results in

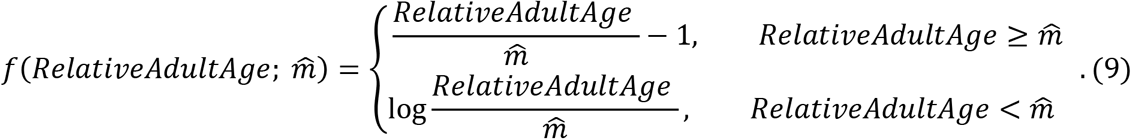

Universal Clock 3 predicts *LoglinearAge* (denoted as *y*). To arrive at an age estimate, we use both equation 4 and equation 6 to arrive at

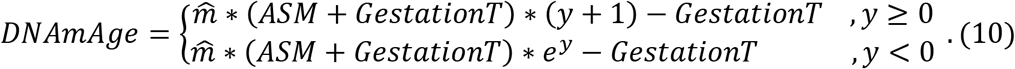

#### Statistics for performance of model prediction

To validate our model, we used DNAm age estimates from LOFO and LOSO analyses, respectively. At each type of estimate, we performed Pearson correlation coefficients and computed median absolute error (MAE) between DNAm based and observed variables across all samples. Correlation and MAE were also computed at species level, limited to the subgroup with n≥15 samples (within a species). We reported the medians for the correlation estimates (med.Cor) and the medians for the MAE estimates (med.MAE) across species. Analogously, we repeated the same analysis at species-tissue level, limited to the subgroup with at least 15 samples (within a species-tissue category).

For **Figure 4** we evaluated the difference (Delta.Age) between the LOSO estimate of DNAmAge and chronological age at half the maximum lifespan (*0.5*maxLifespan*). As expected, Delta.Age=LOSO DNAmAge-(0.5 * *MaxLifespan*) is negatively correlated with species maximum lifespan.

### Array Converter for Mammalian array

Since the human epidemiological cohort data were generated on a different genomic platform (Illumina 450K array), we developed an imputation scheme for converting measurements between the two platforms. The Array Converter from Human 450K Array to our Mammalian array was developed based on a study of n=141 human blood samples that were profiled using both the mammalian array and the Illumina 450k array. Next, we randomly split the data into training (80 percent) and test set (20 percent) ensuring that both data sets had a similar age distribution. In the training set, we fit penalized regressions (with elastic net penalty, alpha=0.5) for each mammalian CpG (dependent variable). To ensure that the predicted beta values lie between 0 and 1, we applied a logit transformation to each target mammalian CpG, 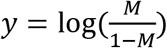, where *M* denotes the beta value. As covariates, we used the subset of CpGs shared between the two array platforms. We did not consider all available covariates/CpGs for a given target mammalian CpG. Rather, we focused on a subset of CpGs located in the genomic interval surrounding each target CpGs with bandwidth *w* upstream and downstream of the target CpG. We selected the bandwidth *w* = 60 *Mb* since it turned out to maximize the accuracy (R squared value) of array conversions in the test set. Finally, the sparse coefficient vectors fitted from the penalized regressions were stored as the array converters from 450K CpGs to the mammalian CpGs. For each mammalian CpG *M*, the imputation follows

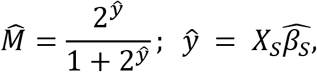

where the set *S* denotes the sparse subset of 450K CpGs corresponding to non-zero coefficient values. The accuracy of an imputed CpGs is guided by robust biweight midcorrelation (bicor) and Pearson correlation. We found that 87% CpGs in Clock 2 and 89% CpGs in Clock 3 exceeded the correlation threshold of 0.6. In calculating the Clock estimates, we replaced the methylation levels by 0.5 for the rest of CpGs that did not satisfy the threshold.

### Epigenetic age acceleration

To adjust for age, we defined epigenetic age acceleration (AgeAccel) as raw residual resulting from regressing DNAm age (from Universal Clocks 2 and 3) on chronological age. By definition, the resulting AgeAccel measure is not correlated with chronological age.

### Human epidemiological cohort studies of time-to-death

We applied our universal Clock 2 and Clock 3 on the 4,651 individuals from (a) the Framingham Heart Study, FHS offspring cohort (n=2,544 Caucasians, 54% females)^75^and (b) Women’s Health Initiative cohort^76,77^ (WHI, n=2107 postmenopausal women, **Supplementary Information**, **note 5**). Methylation levels were profiled in blood samples based on Illumina 450k arrays. In FHS, the mean (SD) chronological age at the time of the blood draw was 66.3 (8.9) years old. During follow-up, 330 individuals died. The mean (SD) follow-up time (used for assessing time-to-death due to all-cause mortality) was 7.8 (1.7) years. The WHI cohort is a national study that enrolled postmenopausal women aged 50-79 years. Our WHI data consists of three ethnic/racial groups: 47% European ancestry (Caucasians), 32% African Americans, and 20% Hispanic ancestry. All the three ethnic groups have marginally the same age distributions, with mean (SD) of 65.4 (7.1) years old. The mean (SD) of follow-up time was16.9 (4.6) years. During the follow-up, 765 women died. Our mortality analysis was peformed in the following. First, we applied our Array Converter algorithm to yield the imputed Mammalian arrays and to estimate DNAm ages based on our universal Clocks. Second, we computed AgeAccel for each cohort. Third, we applied Cox regression analysis for time-to-death (as a dependent variable) to assess the predictive ability of our universal Clocks for all-cause mortality. The analysis was adjusted for age at blood draw and adjusted for gender in FHS. We stratified the WHI cohort by ethnic/racial groups and combined a total of 4 results across FHS and WHI cohorts by fixed effect models weighted by inverse variance. The meta-analysis was performed in R *metafor* function.

### OSKM reprogramming cells in human dermal fibroblasts (HDFs)

We applied our universal Clock 2 and Clock 3 to a previously published reprogramming dataset (GSE54848)^78^. Briefly, human dermal fibroblasts (HDFs) were transfected with OSKM over a 49-day period, and successfully transformed cells were collected and analyzed for DNA methylation with Illumina 450k arrays. Similar to the applications on FHS and WHI cohorts, we applied our Array Converter algorithm to yield the imputed Mammalian arrays and to estimate DNAm ages based on our universal Clocks. The clocks were applied to a total of 27 samples across experiment day 0, 3, 7,11,15,20,28,35, 42 and 49, respectively.

### Murine anti-aging studies

None of the murine samples from the anti-aging studies were used in the training set of the universal clocks.

Universal Clocks 2 and 3 were evaluated in five mouse experiments (independent test data): (1) Snell dwarf mice (n=95), 2) growth hormone receptor knock-out experiment 1 (GHRKO, n=71 samples), (3) GHRKO experiment 2 (n=96 samples), (4) three Tet experiments: Tet1 KO (n=64), Tet2 KO (n=65) and Tet3 KO (n=63), and (5) calorie restriction (n=95). These mouse samples are truly independent test data (i.e. they were not used for training the universal clocks). These experiments were known to prolong the lifespan of mice. We simply performed t-tests to investigate if these conditions slowed epigenetic aging rates based on our universal clocks. As noted, the DNA methylation data in (1) Snell dwarf mice and (3) the GHRKO experiment 2 were profiled in an Illumina 320k customized array for mouse study. DNA methylation data for the other experiments were profiled in our mammalian array. As noted, our Clock2 (816 CpGs) and Clock3 (760 CpGs) contain a total of 1143 unique mammalian CpGs. Of the 1143 CpGs, 165 sites cannot be qualitatively calibrated in the custom 320k array. We simply used the median beta values from the 2205 mice samples that were used for training our universal clocks. Below, we briefly described the experiments. The details for each experiment are listed in **Supplementary Information, note 1 & note 6**.

#### Snell dwarf experiment (n=95)

We studied multiple tissue types from n=47 Snell dwarf^29,30^ and n=48 age matched wild type control mice which were about 6 monts old (mean±SD of age =0.52±0.01 years). We applied the mammalian methylation array to blood, cerebral cortex, liver, kidney, spleen and tail. For most tissue types we compared n=8 dwarf mice to n= 8 age matched wild type control mice with the exception of cerebral cortex where we compared 8 dwarf mice to 7 normal control mice.

#### Full body growth hormone receptor knock-out (n=71)

We studied several mouse tissue types from a growth hormone receptor knock-out (GHRKO, double knock-out in all tissues) study ^31^. We applied the mammalian methylation array to cerebral cortex (n=12), liver (n=23), kidney (n=24) and hippocampus (n=23). GHRKO mice and control mice were age matched: mean±SD [range] of age is 0.66±0.06 [0.57–0.79] years in the GHRKO group and 0.65±0.06 [0.58– 0.79] years in the wild-type control group. To adjust for age, we used the epigenetic age acceleration measure AgeAccel, and related it to genotype (GHRKO versus wild-type) with the non-parametric Kruskal Wallis test.

#### Liver-specific growth hormone receptor knock-out (n=96)

In the second GHRKO experiment, the growth hormone receptor was only knocked out in liver tissue^32^. Unlike the full body GHRKO mice, liver specific GHRKO mice do not exhibit increased lifespan. We applied the mammalian methylation array to blood, cortex, liver, kidney, spleen and tail (n=96 tissue samples). For each tissue type we compared n=8 liver-specific GHRKO mice with n=8 age matched control mice. The mean±SD [range] of age was 0.51±0.03 [0.48,0.59] years among the 96 samples.

#### Calorie restriction study (n=95)

We analyzed 95 male liver samples from a calorie restriction^79^ (CR) study: n=59 in the CR group versus n=36 in the control group. DNA methylation arrays were profiled in both groups. All mice have the same age (1.57 years).

#### *Tet genes* knock-out study

We analyzed three Tet experiments: *Tet1 KO* (*n*=64), *Tet2* KO (n=65) *andTet3* KO (n=63). Each experiment has approximately the same number of samples in *Tet* heterozygous KO and wild-type genotype at each tissue type. DNA methylation array was profiled in both genotypes at each tissue type: cerebral cortex and striatum. All mice have the same age (0.5 years). *Tet1* and *Tet2* knockout mice were purchased from the Jackson Laboratory (*Tet1* KO: Strain #:017358, B6; 129S4-*Tet1^tm1.1Jae^*/J; *Tet2* KO: Strain #:023359, B6(Cg)-*Tet2^tm1.2Rao^/J). Tet3* knockout mice were generated by Yang Lab using the CRISPR/Cas9 method with deletion of exon 4 at the Jackson Laboratory’s Customer Model Generation Core (Founder Strain# 407367, UCLA_GET3672_Tet3_KO_B6J). *Tet3* exon 4 was deleted using the following gRNAs: Up1 AGTATTATTTGGAGACAATT; Up2 TACCAAGAGCAAGTATTATT; Down1 AAATTCCCAAACGGAACCTC; Down2 ACCTGTCCAGAGGTTCCGTT). For each line, the heterozygous knockout mice were bred with C57BL/6J (JAX 000664) to set up the colony. All mice were maintained and bred under standard conditions consistent with National Institutes of Health guidelines and approved by the University of California, Los Angeles Institutional Animal Care and Use Committees. The cages were maintained on a 12:12 light/dark cycle, with food and water provided *ad libitum.* Mice were euthanized by cervical dislocation; cortical and striatal tissues were then quickly dissected out and flash frozen on dry ice until used for DNA extraction.

### Meta-analysis for EWAS of age

In our primary EWAS of age, we focused on the samples from eutherians (n=65 species) that each species has at least 15 samples from the same tissue type. In secondary analyses, we also studied aging effects in marsupials (n=4 marsupial species that had at least 10 same tissue type samples) and monotremes (only n=2 species).

Our meta-analysis for EWAS of age in eutherian species combined Pearson correlation test statistics across species-tissue strata that contained at least 15 samples each. The minimum sample size requirement resulted in 143 species-tissue strata from 65 eutherian species (**Data S1.5**). To counter the dependency patterns resulting from multiple tissues from the same species, the meta-analysis was carried out in two steps. First, we meta-analyzed the EWAS of different tissues for each species separately. These tissue specific summary statistics were combined within the same species to represent the EWAS results at species level. Second, we meta-analyzed the resulting 65 species EWAS results across species to arrive at the final meta-EWAS of age. In each meta-analysis step, we used unweighted Stouffer’s method as implemented in R. In more details, we gathered 68 blood samples from 27 distinct lemur species and 23 skin samples from 23 distinct lemur species, each species-tissue stratum with at most 3 samples. We therefore combined those 68 blood samples to perform blood EWAS in lemurs. Similarly, we combined the 23 skin samples for skin EWAS in lemurs. As listed in **Data 1.5**, the combined species in lemurs was denoted by Strepsirrhine in the column of “Species Latin Name”.

EWAS of age in marsupials was based on two-step meta-analysis where we relaxed the threshold of sample size in species-tissue category to n≥10 (**Data S1.12**). Due to a small sample size in monotremes (n=15), we combined all monotreme samples into a single data set.

#### Brain EWAS

We applied the two-step meta-analysis approach to the brain EWAS results based on more than 900 brain tissues (cerebellum, cortex, hippocampus, hypothalamus, striatum, subventricular zone, and whole brain) from eight species including human, vervet monkey, mice, olive baboon, brown rat, and pig species (**Data S1.6**).

#### EWAS of a single tissue

For the single brain region in cerebral cortex, we simply combined the tissue specific EWAS results across different species based on unweighted Stouffer’s method (**Data S1.7**). Similarly, we carried out the one-step meta-analysis EWAS of blood, liver, muscle, and skin (**Data S1.8—S1.11**). We combined blood and spleen samples into one category (blood) because unsupervised hierarchical clustering shows that the two sources of DNA are highly comparable. In an analogous manner, skin and ear samples were combined into one category (skin). Blood EWAS results were combined across six taxonomic orders (n=4,513) including 584 tissues from humans, 742 from dogs, 406 from mice, 36 from killer whales, 143 from bottlenose dolphins, 83 from Asian elephants. Skin EWAS results were combined across seven orders (n=2,363) including 79 from bowhead whales, 667 tissues from 19 bat species, 183 from killer whales, 107 from naked mole rats, 74 from humans. Liver EWAS results were combined across four orders (n=1,063) including 600 mice, 97 from humans, 48 from horses, etc. Muscle EWAS results were combined across three orders (n=354) including 24 from evening bats, 57 from humans, and 21 from naked mole rats, etc. Cerebral cortex EWAS results were combined across two orders (n=375) including, 46 from humans, 48 from vervet monkeys and 15 from naked mole rats.

#### Stratification by age groups

To assess whether the age related CpGs in young animals relate to those in old animals, we split the data into three age groups: young age (age<1.5* age at sexual maturity, ASM), middle age (age between 1.5 and 3.5 ASM), and old age group (age≥3.5 ASM). The threshold of sample size in species-tissue was relaxed to n≥10. The age correlations in each age group were meta-analyzed using the above-mentioned two-step metaanalysis approach.

### Polycomb repressive complex

Polycomb repressive complex annotations were defined based on the binding of at least two transcriptional factor members of polycomb repressor complex 1 (PRC1 with subgroups RING1, RNF2, BMI1) or 2 (PRC2 with subgroups EED, SUZ12, and EZH2) in 49 available ChipSeq datasets in ENCODE^43^.

We identified 640 and 5,287 CpGs on the array that were located in regions bound by PRC1 and PRC2, respectively. We performed a one-sided hypergeometric analysis to study both the enrichment (Odds ratios [OR] >1) and depletion (OR<1) patterns for our age-related markers based on the top 1,000 CpGs increased with age and the top 1,000 CpGs decreased with age from EWAS of age.

### Universal chromatin state analysis

To annotate our age-related CpGs based on chromatin states, we assigned a state for all our mammalian CpGs based on a recently published universal ChromHMM chromatin state annotation of the human genome^36^. The underlying hidden Markov model (HMM) was trained with over 1,000 datasets of 32 chromatin marks in more than 100 cell and tissue types. This model then produced a single chromatin state annotation per genomic position that is applicable across cell and tissue types, as opposed to producing an annotation that is specific to one cell or tissue type. A total of 100 distinct states were generated and categorized into 16 major groups according to the parameters of the model and external genome annotations^56^ (described in **Data S5.2**). We performed a one-sided hypergeometric analysis to study both the enrichment (Odds ratios [OR] >1) and depletion (OR<1) patterns for our age-related markers based on the top 1,000 CpGs with a positive correlation with age and the top 1,000 CpGs with a negative correlation with age across different eutherian species.

### Analysis of late-replicating domains

To annotate our age-related CpGs based on late-replicating regions, we used the annotations (Hg19 and mm10) conducted by Zhou et al. ^40^, which is available at https://zwdzwd.github.io/pmd. Based on PMD-HMD structure, each CpG can be assigned to three categories: common PMD, common HMD or neither. Briefly, DNA methylation data was profiled from tumor and adjacent normal tissues based on WGBS array at 1.5x coverage. PMD/HMD domains were first identified based on HMM stochastic models. The dichotomized status of PMD and HMD (in 100kb bin) were later defined using a Gaussian-Mixture model. The set of *“common* PMD” was defined using a more stringent threshold to distinguish from *“common* HMD”. We also annotated a CpG based on solo-WCGW region, which is prone to hypomethylation. For Hg19, 12527 CpGs (33.5%) are in common PMD, 13589 CpGs (30.9%) are in common HMD, 10087 CpGs (24.9%) are in neither group, and 1289 CpGs (3.4%) are missing for the category. For mm10, 2615 CpGs (0.09%) are in common PMD, 10593 CpGs (35.7%) are in common HMD, 15306 CpGs (51.6%) are in neither group, and 1123 CpGs (3.8%) are missing for the category. Similar to the annotation analysis based on universal chromatin state, we performed one-sided hypergeometric analysis to study the overlap of our age-related CpGs with a) common PMD/HMD structures and (b) solo-WCGW structures: genome-wide (solo-WCGW) and those in the common PMD regions (solo-WCGW common PMDs). Background in the hypergeometric analysis was adjusted accordingly based on the genome assembly used in an analysis.

### GREAT enrichment analysis

We applied the GREAT analysis software tool^41^ to the top 1,000 positively age-related and the top 1,000 negatively age-related CpGs from EWAS of age. GREAT implemented foreground/background hypergeometric tests over genomic regions where we input all 37,492 CpGs of the mammalian array as background and the genomic regions of the 1,000 CpGs as foreground. This yielded hypergeometric P-values not confounded by the number of CpGs within a gene. We performed the enrichment based on the settings (human genome assembly: hg19, Proximal: 5.0 kb upstream, 1.0 kb downstream, plus Distal: up to 50 kb) for about 76,290 gene sets associated with GO terms, MSigDB (including gene sets for upstream regulators), PANTHER, KEGG pathway, disease ontology, gene ontology, and human and mouse phenotypes. We report the gene sets with FDR<0.05, nominal hypergeometric *P*-values <0.001, and number of overlapping genes ≥3.

### EWAS-TWAS overlap analysis

Our EWAS-TWAS based overlap analysis related the gene sets found by our EWAS of age with the gene sets from our in-house TWAS database. To build the TWAS database, we collected published large-scale bulk or single-cell transcriptomic studies of chronological age. We also downloaded and integrated curated datasets of known aging-related genes from GenAge^44^ and Enrichr^45,46^ website (Aging_Perturbations_from_GEO, http://amp.pharm.mssm.edu/Enrichr/). The GenAge database includes genes related to aging in human and model organisms (Tacutu et al., 2018). Enrichr(Chen et al., 2013; Kuleshov et al., 2016) using a crowdsourcing method to extract gene expression signatures of aging from GEO. In total, our in-house TWAS database contains more than 700 aging-related gene sets across various species and tissue/cell types. For each TWAS gene set, we pruned genes present in our mammalian array. For each EWAS result, we studied the genomic regions from the top 1,000 CpGs which increased and decreased with age, respectively. CpGs were annotated to their proximal genes based on human genomic location. The gene assignments are highly orthologous across species^8^. To assess the overlap, we performed hypergeometric analysis and reported one-sided P-value for enrichment assessment. The number of background genes in the hypergeometric test was the total number of TWAS species-specific orthologous genes in our mammalian array. Different from the GREAT analysis conducted in genomic region, the hypergeometric analysis was performed at gene level to annotate TWAS results. In order to avoid the bias of the array, we conducted permutation tests using the following steps: 1) Randomly sample 1,000 CpGs among any on the array and conduct EWAS-TWAS on the permutation CpGs. For each TWAS gene set, a right-tail P-value is calculated. 2) Repeat Step 1 1,000 times to populate a list of P-values and model the resulting null distribution with a Gamma function. The cumulative distribution function (CDF) is also calculated. 3) Calculate the permutation P-value based on the CDF from step 2. In our study, we restricted the TWAS with permutation *P*<0.05, and highlighted the top 10 TWAS gene sets based on the unadjusted hypergeometric *P*-value. We only reported these TWAS results with ≥5 overlap genes.

### EWAS-GWAS overlap analysis

Our EWAS-GWAS overlap analysis related the gene sets found by our EWAS of age with the gene sets found by published large-scale GWAS of various phenotypes, across body fat distribution, lipid panel outcomes, metabolic outcomes, neurological diseases, six DNAm-based biomarkers, and other age-related traits (**Supplementary Information**, **note 4**). A total of 102 GWAS results were studied. The six DNAm biomarkers included four epigenetic age acceleration measures derived from 1) pantissue epigenetic age adjusted for age-related blood cell counts referred to as intrinsic epigenetic age acceleration (IEAA)^4,80^, 2) Hannum’s blood-based DNAm age^18^; 3) DNAmPhenoAge^20^; and 4) the mortality risk estimator DNAmGrimAge^21^, along with DNAm-based estimates of blood cell counts and plasminogen activator inhibitor 1 (PAI1) levels^21^. For each GWAS result, we used the MAGENTA software to calculate an overall GWAS *P*-value per gene, which is based on the most significant SNP association *P*-value within the gene boundary (± 50 kb) adjusted for gene size, number of SNPs per kb, linkage disequilibrium, and other potential confounders^81^. The MAGENTA analysis was performed in MATLAB (2017 version). We restricted the analysis to genomic regions of GWAS genes present on the mammalian array. For each EWAS result, we studied the genomic regions from the top 1,000 CpGs with positive and negative age correlations, respectively. To assess the overlap with a test trait, we selected the top 2.5 % genes for each GWAS trait and calculated one-sided hypergeometric *P*-values based on genomic regions. We report GWAS traits that led to a significant hypergeometric test (nominal *P*-value <5×10^-4^, Bonferroni corrected *P*<0.05) for any EWAS of age. The number of background genomic regions in the hypergeometric test was based on the overlap between all genes in the GWAS and all genomic regions represented by the mammalian array.

### Transcription factor binding

We used the CellBase database^42^ with the ENCODE^43^ transcription factor binding sites. A total of 186 TFs across 70 genes could be mapped to CpGs on our Mammalian array. The Chipseq analysis of these TFs was done in 17 different cell types depending on the target molecule (**Data S11.1**). We annotated a CpG with specific TF if it overlapped with the respective binding site in at least one cell type (in Hg19 assembly).

TFs with extreme annotation size (number of genes across binding regions < 5 or >2000) were removed, yielding 68 TFs remained in our analysis.

### Single Cell ATAC-seq analysis for the top age-related CpGs

Recent advances have enabled the sequencing of ATAC profiles within single cells enabling an assessment of the proportion of cells containing an open chromatin region ^47^. We cross referenced the top 35 CpGs with positive age correlation across mammalian tissues with publicly available single cell ATAC-Seq analysis (**Table S1**). We downloaded 10X Multiome count data in AnnData format as H5AD from GEO (https://www.ncbi.nlm.nih.gov/geo/query/acc.cgi?acc=GSE194122). The ATAC array data were managed using the Python package anndata. Hg38 ATAC peak locations were extracted from the metadata “.var” section using anndata. Peak locations were overlapped with probe locations using GenomicRanges used^82^ for the top 35 CpG sites. The overlapping peaks were then used to extract the processed counts for each cell using the anndata. The proportion of cells with each ATAC peak was calculated per individual and was correlated with the age of each sample. The cell type for each barcode was extracted from the observable object using the anndat. We subsequently computed the proportion of each cell type containing an ATAC peak in one of the 7 significantly correlated regions. Progenitor cells were grouped as HSC, MK/E prog, G/M prog, Lymph prog and Proerythroblast and differentiated cells as CD14+ Mono, CD16+ Mono, CD8+ T naive, CD8+ T, CD4+T naive, CD4+ T activate, Naive CD20+ B, B1 B, Transitional B and NK. Percentage of each of the progenitor and differentiated populations was calculated and the proportion of cells containing an ATAC peak in one of the 7 significantly correlated regions was calculated. To confirm enrichment for the hyper methylated sites showing decrease of chromatin accessibility with age, we randomly selected 1000 sets of 17 ATAC peaks and compared the mean correlation with age of the selected regions to the 1000 sampled sets of regions.

### URLs

AnAge, http://genomics.senescence.info/help.html#anage,

GREAT, http://great.stanford.edu/public/html/

Late-replicating domains: https://zwdzwd.github.io/pmd

UCSC genome browser: http://genome.ucsc.edu/index.html

## ACKNOWLEDGEMENTS and FUNDING

This work was mainly supported by the Paul G. Allen Frontiers Group (SH). Additional support was also provided by Open Philanthropy/Silicon Valley Fund (SH and Ken Raj). Julie A. Mattison and the NHP Core are supported by the Intramural Research Program, National Institute on Aging, NIH. Plains zebra sample collection was supported by National Geographic Society grant #8941-11. We acknowledge the Museum of Vertebrate Zoology and Chris J. Conroy from the University of California, Berkeley. We acknowledge Dr. Richard Miller and his lab (http://www.richmillerlab.com/long-lived-mutants) for providing various dwarf mice and controls (Snell dwarf mouse, GHRKO experiments).

The Framingham Heart Study is funded by National Institutes of Health contract N01-HC-25195 and HHSN268201500001I. The laboratory work for this investigation was funded by the Division of Intramural Research, National Heart, Lung, and Blood Institute, National Institutes of Health. The analytical component of this project was funded by the Division of Intramural Research, National Heart, Lung, and Blood Institute, and the Center for Information Technology, National Institutes of Health, Bethesda, MD.

The Women’s Health Initiative program is funded by the National Heart, Lung, and Blood Institute, National Institutes of Health, U.S. Department of Health and Human Services through contracts HHSN268201600018C, HHSN268201600001C, HHSN268201600002C, HHSN268201600003C, and HHSN268201600004C. The authors thank the WHI investigators and staff for their dedication, and the study participants for making the program possible. A full listing of WHI investigators can be found at: http://www.whi.org/researchers/Documents%20%20Write%20a%20Paper/WHI%20Investigator%20Long%20List.pdf. The views expressed in this manuscript are those of the authors and do not necessarily represent the views of funding bodies such as the National Heart, Lung, and Blood Institute; the National Institutes of Health; or the U.S. Department of Health and Human Services.

## Data availability

All data from the Mammalian Methylation Consortium will be posted on Gene Expression Omnibus. The list of the GSE (GSE174758, GSE184211, GSE184213, GSE184215, GSE184216, GSE184218, GSE184220, GSE184221, GSE184224, GSE190660, GSE190661, GSE190662, GSE190663, GSE190664, GSE174544, GSE190665,GSE174767,GSE184222,GSE184223,GSE174777,GSE174778,GSE1733 30,GSE164127,GSE147002,GSE147003,GSE147004) can be found in **Supplementary information note 2**. The mammalian methylation array is available through the non-profit Epigenetic Clock Development Foundation (https://clockfoundation.org/).

## R code for universal pan-mammalian clocks

The R codes for the three universal pan-mammalian clocks can be found in **Supplementary information note 7**.

## CONTRIBUTIONS

Ake T. Lu, Zhe Fei, Caesar Li, Joseph Zoller, SH developed the universal clocks. ATL, Amin Haghani, Robert Lowe, Qi Yan, Charles Breeze, Michael Thompson, Matteo Pellegrini, Ha Vu, Wanding Zhou, SH carried out additional bioinformatics analyses. Adriana Arneson, Jason Ernst, SH designed the mammalian methylation array. ATL, ZF, AH, RL, QY, Ken Raj, SH drafted the first version of the article. The remaining authors contributed tissues or DNA samples or helped with the data generation process.

All authors helped with editing the article and data interpretation. SH conceived the study and design.

## COMPETING INTERESTS

SH is a founder of the non-profit Epigenetic Clock Development Foundation which plans to license several patents from his employer UC Regents. These patents list SH, JE and AA as inventors. The other authors declare no conflicts of interest.

## CORRESPONDING AUTHOR

Correspondence to Steve Horvath

## Notes

### Summary of Updates

All sections updated with more species and samples. New sections added for analysis in human mortality, OSKM-based reprogramming, transgenic mice for studying the somatotropic axis, overlap with late-replicating domains, TF enrichment and single cell ATAC seq analysis of human bone marrow.

