## SupplementaryInfo for "Universal DNA methylation age across mammalian tissues"

**Supplementary Material**

for the article entitled “***Universal DNA methylation age across mammalian tissues***"

AT Lu *et al.*

#### Supplementary Figures

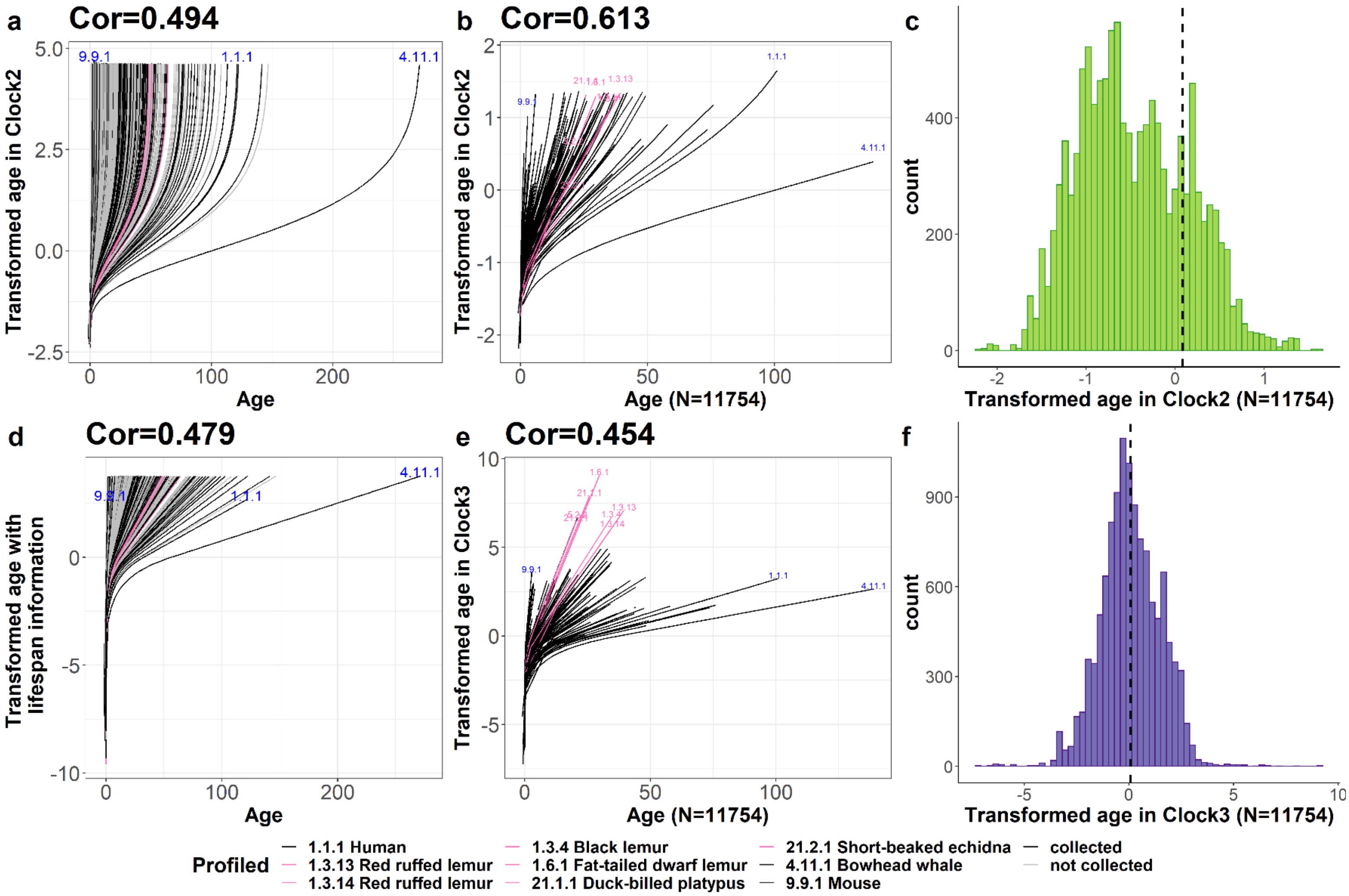


**Figure S1. Transformed age in universal clocks**. The plot displays transformed age in universal Clock 2 (**a--c**) and universal Clock 3 (**d-f**). (**a, b**) Loglog transformation of Relative Age (y-axis) versus age in universal Clock 2 and (**d, e**) log-linear age (y-axis) versus age in our universal Clock 3. Of the 969 mammalian species with available gestation time, age at sexual maturity and maximum lifespan in AnAge database, 339 species are available in our collection. We multiplied the reported maximum lifespan of non-human or non-mouse species by 1.3. Transformed ages were calculated for all the 969 species with imputed age ranging from gestation time through the modified maximum lifespan. The columns (**a,d**) display all the 969 species with the imputed ages. In panel d, we proposed the log-linear age with the parameter $\boldsymbol{m}$ formulated with maximum lifespan as the information is available for all species ( $\boldsymbol{m}^* = c_1 * \frac{MaxLifespan+GestationT}{ASM+GestationT}$ in Methods). Of the 339 species, 185 species with age information of high confidence and known tissue types were used in training universal clocks. The columns (**b, e**) empirically display these 185 species with the age variable (x-axis) based on the observed ages from all the samples in our collection (N=11,754). In panel **e**, we applied the log-linear age formulated without knowing maximum lifespan to train Clock 3 (formula (5) in Methods). Each line represents a species marked by gray for non-profiled and marked by black or pink for profiled species in our collection, as listed in the legend. Some species such as lemurs with relatively short gestation time in regressing $\boldsymbol{m}^*$ (formula (7) in Methods) exhibiting high log-linear ages in (**e**) are marked in pink. Each panel reports the Pearson correlation coefficient. (**c,f**) display the histograms of transformed ages based on the all the samples from the 187 species with vertical lines presenting at means.

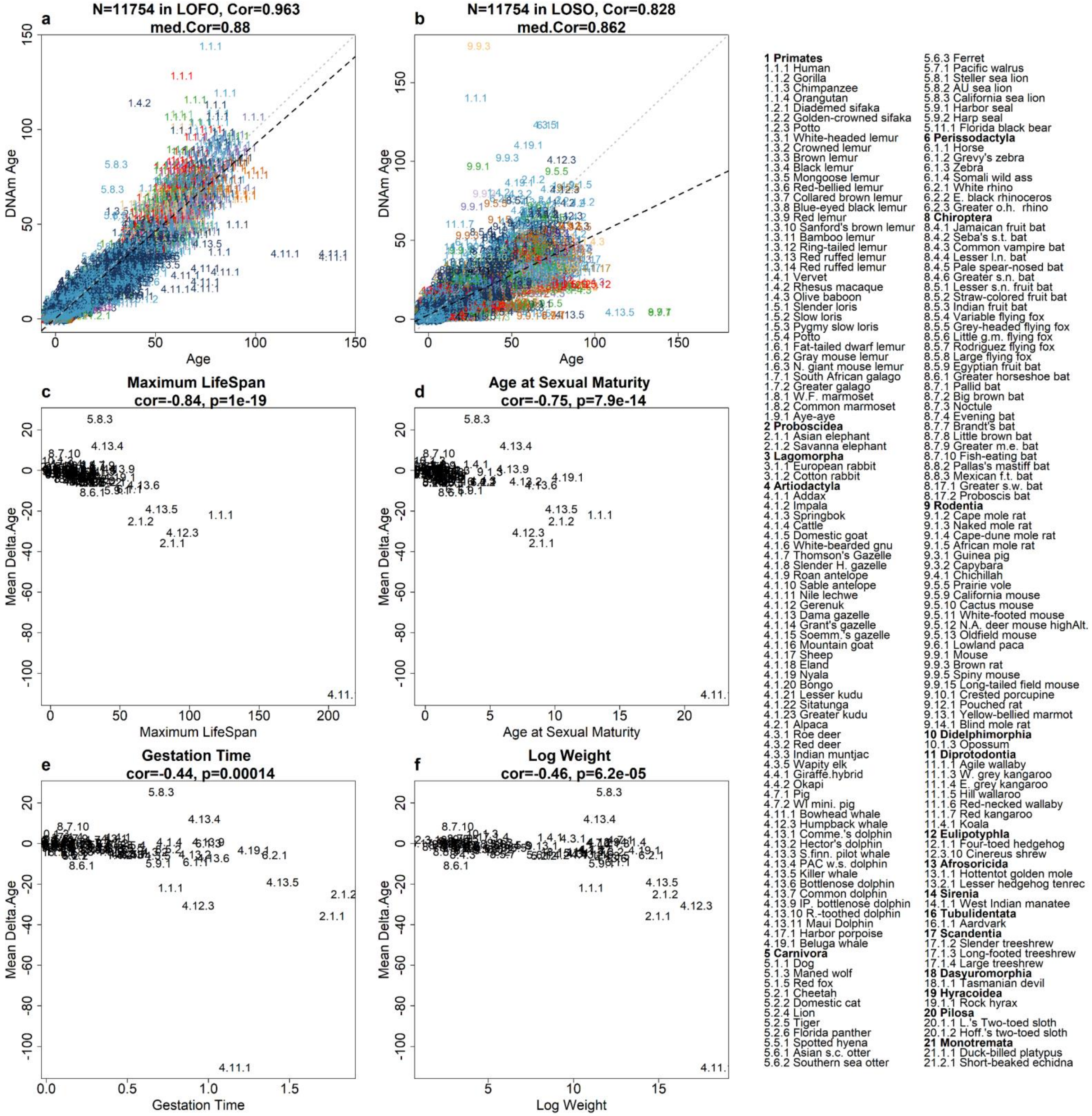

a
N=11754 in LOFO, Cor=0.963
med.Cor=0.88
DNAm Age
Age
b
N=11754 in LOSO, Cor=0.828
med.Cor=0.862
DNAm Age
Age
c
Maximum LifeSpan
cor=-0.84, p=1e-19
Mean Delta.Age
Maximum LifeSpan
d
Age at Sexual Maturity
cor=-0.75, p=7.9e-14
Mean Delta.Age
Age at Sexual Maturity
e
Gestation Time
cor=-0.44, p=0.00014
Mean Delta.Age
Gestation Time
f
Log Weight
cor=-0.46, p=6.2e-05
Mean Delta.Age
Log Weight
**1 Primates**
1.1.1 Human
1.1.2 Gorilla
1.1.3 Chimpanzee
1.1.4 Orangutan
1.2.1 Diademed sifaka
1.2.2 Golden-crowned sifaka
1.2.3 Potto
1.3.1 White-headed lemur
1.3.2 Crowned lemur
1.3.3 Brown lemur
1.3.4 Black lemur
1.3.5 Mongoose lemur
1.3.6 Red-bellied lemur
1.3.7 Collared brown lemur
1.3.8 Blue-eyed black lemur
1.3.9 Red lemur
1.3.10 Sanford's brown lemur
1.3.11 Bamboo lemur
1.3.12 Ring-tailed lemur
1.3.13 Red ruffed lemur
1.3.14 Red ruffed lemur
1.4.1 Vervet
1.4.2 Rhesus macaque
1.4.3 Olive baboon
1.5.1 Slender loris
1.5.2 Slow loris
1.5.3 Pygmy slow loris
1.5.4 Potto
1.6.1 Fat-tailed dwarf lemur
1.6.2 Gray mouse lemur
1.6.3 N. giant mouse lemur
1.7.1 South African galago
1.7.2 Greater galago
1.8.1 W.F. marmoset
1.8.2 Common marmoset
1.9.1 Aye-aye
**2 Proboscidea**
2.1.1 Asian elephant
2.1.2 Savanna elephant
**3 Lagomorpha**
3.1.1 European rabbit
3.1.2 Cotton rabbit
**4 Artiodactyla**
4.1.1 Addax
4.1.2 Impala
4.1.3 Springbok
4.1.4 Cattle
4.1.5 Domestic goat
4.1.6 White-bearded gnu
4.1.7 Thomson's Gazelle
4.1.8 Slender H. gazelle
4.1.9 Roan antelope
4.1.10 Sable antelope
4.1.11 Nile lechwe
4.1.12 Gerenuk
4.1.13 Dama gazelle
4.1.14 Grant's gazelle
4.1.15 Soemm.'s gazelle
4.1.16 Mountain goat
4.1.17 Sheep
4.1.18 Eland
4.1.19 Nyala
4.1.20 Bongo
4.1.21 Lesser kudu
4.1.22 Sitatunga
4.1.23 Greater kudu
4.2.1 Alpaca
4.3.1 Roe deer
4.3.2 Red deer
4.3.3 Indian muntjac
4.3.5 Wapity elk
4.4.1 Giraffe.hybrid
4.4.2 Okapi
4.7.1 Pig
4.7.2 WI mini. pig
4.11.1 Bowhead whale
4.12.3 Humpback whale
4.13.1 Comme.'s dolphin
4.13.2 Hector's dolphin
4.13.3 S.finn. pilot whale
4.13.4 PAC w.s. dolphin
4.13.5 Killer whale
4.13.6 Bottlenose dolphin
4.13.7 Common dolphin
4.13.9 IP. bottlenose dolphin
4.13.10 R.-toothed dolphin
4.13.11 Maui Dolphin
4.17.1 Harbor porpoise
4.19.1 Beluga whale
**5 Carnivora**
5.1.1 Dog
5.1.3 Maned wolf
5.1.5 Red fox
5.2.1 Cheetah
5.2.2 Domestic cat
5.2.4 Lion
5.2.5 Tiger
5.2.6 Florida panther
5.5.1 Spotted hyena
5.6.1 Asian s.c. otter
5.6.2 Southern sea otter
5.6.3 Ferret
5.7.1 Pacific walrus
5.8.1 Steller sea lion
5.8.2 AU sea lion
5.8.3 California sea lion
5.9.1 Harbor seal
5.9.2 Harp seal
5.11.1 Florida black bear
**6 Perissodactyla**
6.1.1 Horse
6.1.2 Grevy's zebra
6.1.3 Zebra
6.1.4 Somali wild ass
6.2.1 White rhino
6.2.2 E. black rhinoceros
6.2.3 Greater o.h. rhino
**8 Chiroptera**
8.4.1 Jamaican fruit bat
8.4.2 Seba's s.t. bat
8.4.3 Common vampire bat
8.4.4 Lesser l.n. bat
8.4.5 Pale spear-nosed bat
8.4.6 Greater s.n. bat
8.5.1 Lesser s.n. fruit bat
8.5.2 Straw-colored fruit bat
8.5.3 Indian fruit bat
8.5.4 Variable flying fox
8.5.5 Grey-headed flying fox
8.5.6 Little g.m. flying fox
8.5.7 Rodriguez flying fox
8.5.8 Large flying fox
8.5.9 Egyptian fruit bat
8.6.1 Greater horseshoe bat
8.7.1 Pallid bat
8.7.2 Big brown bat
8.7.3 Noctule
8.7.4 Evening bat
8.7.7 Brandt's bat
8.7.8 Little brown bat
8.7.9 Greater m.e. bat
8.7.10 Fish-eating bat
8.8.2 Pallas's mastiff bat
8.8.3 Mexican f.t. bat
8.17.1 Greater s.w. bat
8.17.2 Proboscis bat
**9 Rodentia**
9.1.2 Cape mole rat
9.1.3 Naked mole rat
9.1.4 Cape-dune mole rat
9.1.5 African mole rat
9.3.1 Guinea pig
9.3.2 Capybara
9.4.1 Chichillah
9.5.5 Prairie vole
9.5.9 California mouse
9.5.10 Cactus mouse
9.5.11 White-footed mouse
9.5.12 N.A. deer mouse highAlt.
9.5.13 Oldfield mouse
9.6.1 Lowland paca
9.9.1 Mouse
9.9.3 Brown rat
9.9.5 Spiny mouse
9.9.15 Long-tailed field mouse
9.10.1 Crested porcupine
9.12.1 Pouched rat
9.13.1 Yellow-bellied marmot
9.14.1 Blind mole rat
**10 Didelphimorphia**
10.1.3 Opossum
**11 Diprotodontia**
11.1.1 Agile wallaby
11.1.3 W. grey kangaroo
11.1.4 E. grey kangaroo
11.1.5 Hill wallaroo
11.1.6 Red-necked wallaby
11.1.7 Red kangaroo
11.4.1 Koala
**12 Eulipotyphla**
12.1.1 Four-toed hedgehog
12.3.10 Cinereus shrew
**13 Afrosoricida**
13.1.1 Hottentot golden mole
13.2.1 Lesser hedgehog tenrec
**14 Sirenia**
14.1.1 West Indian manatee
**16 Tubulidentata**
16.1.1 Aardvark
**17 Scandentia**
17.1.2 Slender treeshrew
17.1.3 Long-footed treeshrew
17.1.4 Large treeshrew
**18 Dasyuromorphia**
18.1.1 Tasmanian devil
**19 Hyracoidea**
19.1.1 Rock hyrax
**20 Pilosa**
20.1.1 L.'s Two-toed sloth
20.1.2 Hoff.'s two-toed sloth
**21 Monotremata**
21.1.1 Duck-billed platypus
21.2.1 Short-beaked echidna

**Figure S2**. **Basic universal clock for log-transformed age. a, b,** Chronological age (x-axis) versus DNAmAge estimated using a, leave-one-fraction-out (LOFO) and b, leave-one-species-out (LOSO) analysis. The gray and black dashed lines correspond to the diagonal line (y=x) and the regression line, respectively. Each dot (tissue sample) is labeled by the mammalian species index (legend). The species index corresponds to the taxonomic order, e.g. 1=primates, 2=elephants (Proboscidea) etc. (legend). The numbers after the first and second decimal points enumerate the taxonomic family and species, respectively. Points are colored by tissue type (Data S1.4). The heading of each panel reports the Pearson correlation (cor) across all samples. Here med.Cor (or med.MAE) denotes the median value across species that contain at least 15 samples. **c–f**, The y-axis reports the mean difference between the LOSO estimate of DNAm age and chronological age evaluated at a fixed age defined as half the maximum lifespan (denoted as Mean Delta.Age). The scatter plots depict mean delta half lifespan per species (y-axis) versus c, maximum lifespan observed in the species, d, average age at sexual maturity e, gestational time (in units of years), and f, (log-transformed) average adult body mass in units of grams.

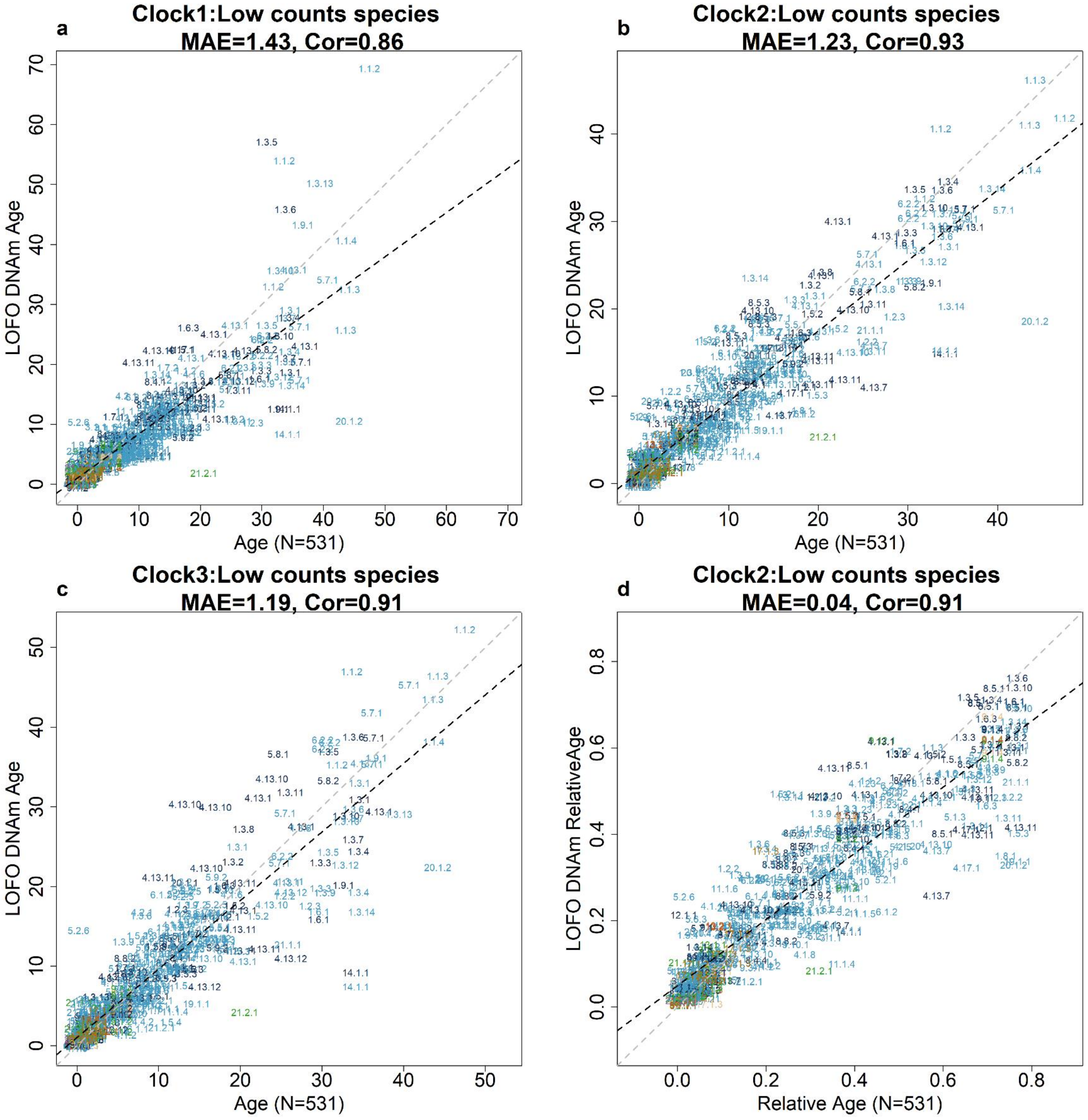


**Figure S3. Universal clocks applied to species with fewer than 15 samples**. The title of each panel lists the type of universal clock: **a,** Clock 1=basic universal clock based on log(Age+2), **b,d**, Clock 2=universal clock for relative age, **c,** Clock 3 =universal clock for log-linear age. Leave-one-fraction-out (LOFO) methylation estimates versus a–c, chronological age or d, relative age. The respective inverse transformations were applied to arrive at DNA methylation-based estimates of chronological age in years or relative age (y-axis).

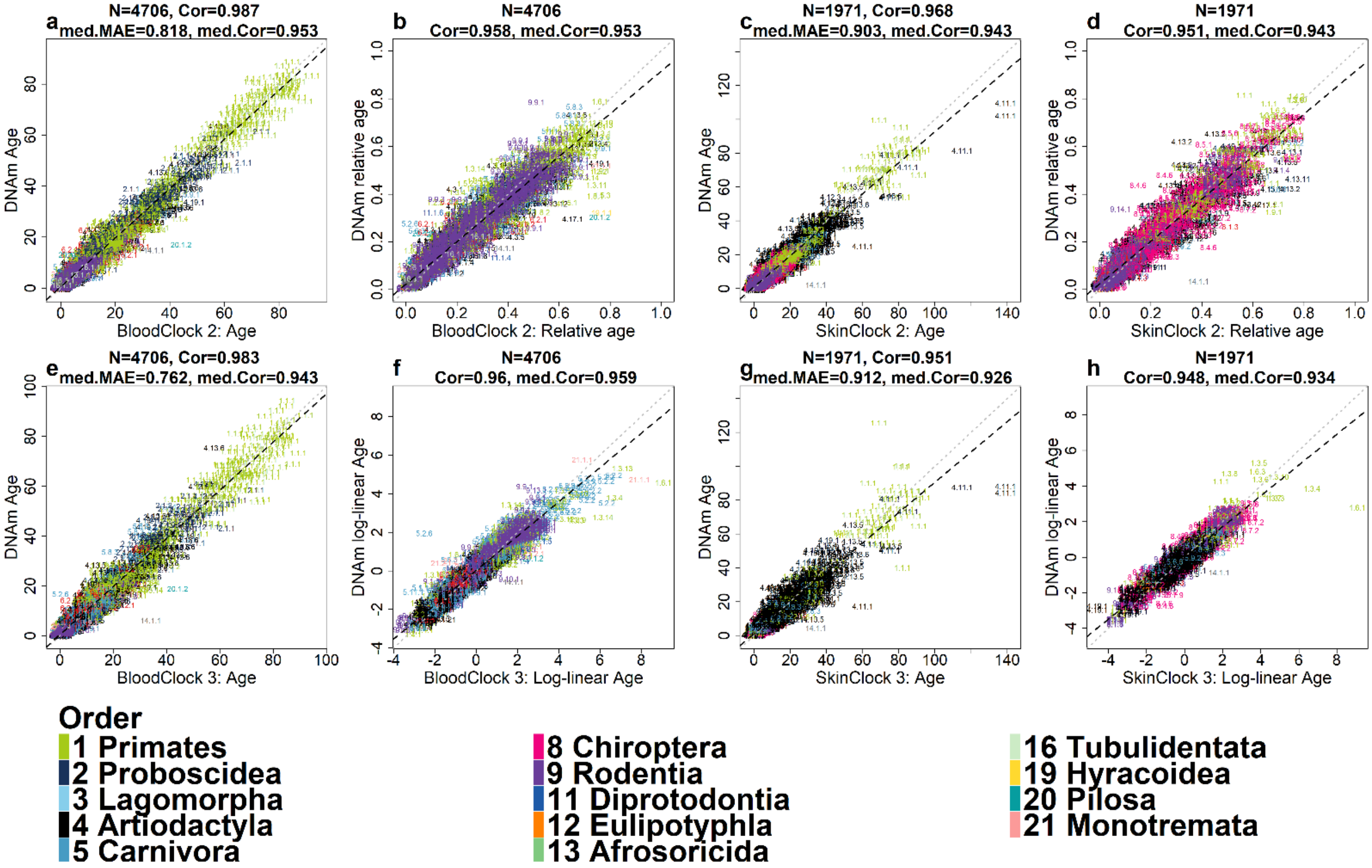


**Figure S4. Universal clocks for specific tissues (blood, skin).** These tissue-specific universal clocks were constructed in an analogous fashion to the pan-tissue clocks described in the main text. The panels show leave-one-fraction-out (LOFO) estimates (y-axis) of four clocks: blood universal clock 2 (BloodClock 2) which estimates relative age, blood universal clock 3 (BloodClock 3) which estimates log-linear transformation of age. Analogously, we defined SkinClock2 and SkinClock3. Relative age estimation incorporates maximum lifespan and gestational age, and assumes values between 0 and 1. Log-linear age is formulated with age at sexual maturity and gestational time. **a,c,e, g**, LOFO estimates of DNAm age (y-axis, in units of years) based on transforming relative age (Clock 2) or log-linear age (Clock 3). **b,f,d,h**, transformed age (x-axis) versus corresponding DNAm estimates (y-axis). The title of each panel reports the Pearson correlation coefficient across all data points and the median correlation (med.Cor) and median of median absolute error (med.MAE) across all species. Each sample is labeled by mammalian species index (explained in Figure 2) and colored by taxonomic order. The legend reports the taxonomic order and the mammalian order index as a prefix.

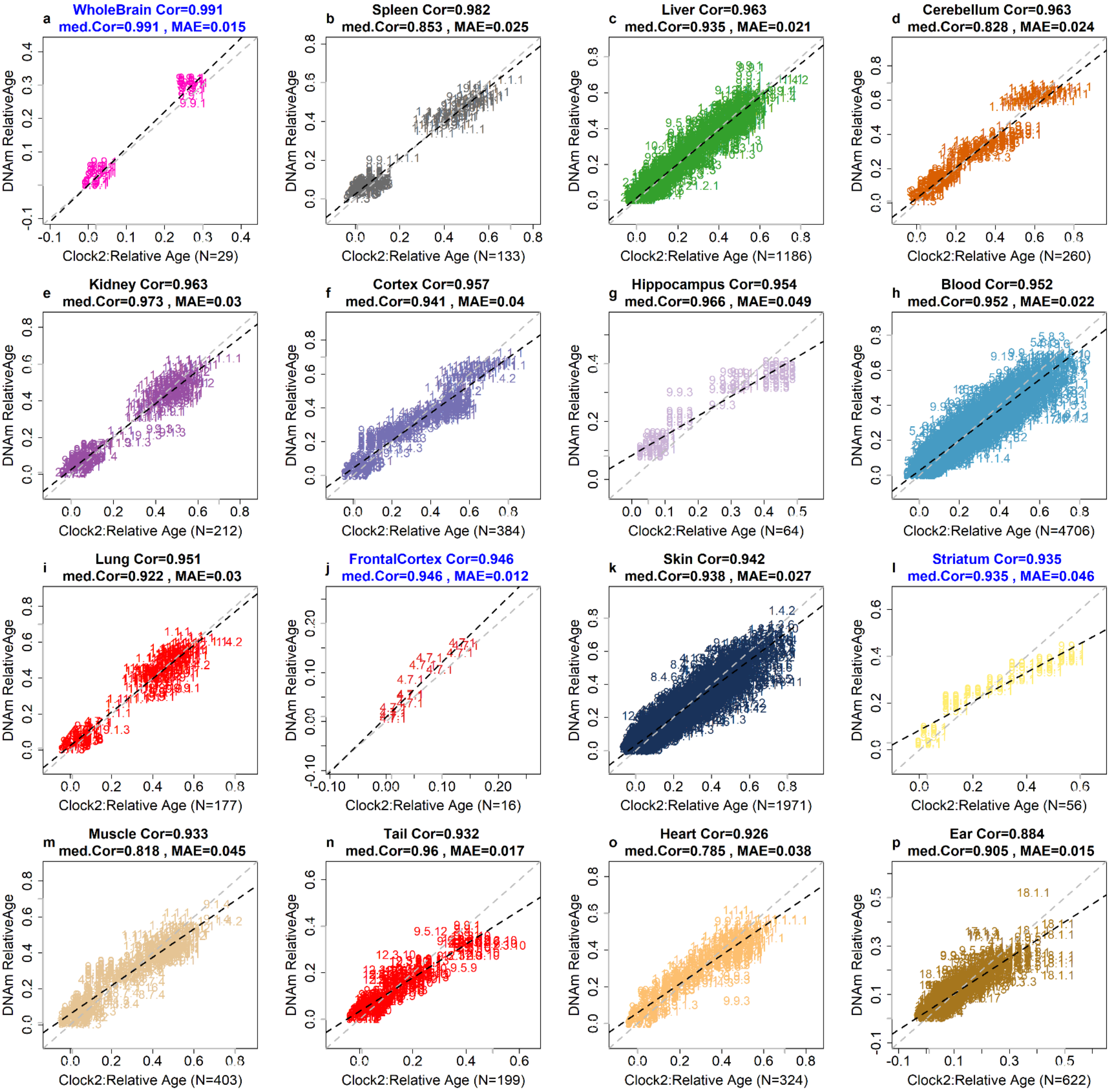

a
WholeBrain Cor=0.991
med.Cor=0.991 , MAE=0.015
DNAm RelativeAge
Clock2:Relative Age (N=29)
b
Spleen Cor=0.982
med.Cor=0.853 , MAE=0.025
DNAm RelativeAge
Clock2:Relative Age (N=133)
c
Liver Cor=0.963
med.Cor=0.935 , MAE=0.021
DNAm RelativeAge
Clock2:Relative Age (N=1186)
d
Cerebellum Cor=0.963
med.Cor=0.828 , MAE=0.024
DNAm RelativeAge
Clock2:Relative Age (N=260)
e
Kidney Cor=0.963
med.Cor=0.973 , MAE=0.03
DNAm RelativeAge
Clock2:Relative Age (N=212)
f
Cortex Cor=0.957
med.Cor=0.941 , MAE=0.04
DNAm RelativeAge
Clock2:Relative Age (N=384)
g
Hippocampus Cor=0.954
med.Cor=0.966 , MAE=0.049
DNAm RelativeAge
Clock2:Relative Age (N=64)
h
Blood Cor=0.952
med.Cor=0.952 , MAE=0.022
DNAm RelativeAge
Clock2:Relative Age (N=4706)
i
Lung Cor=0.951
med.Cor=0.922 , MAE=0.03
DNAm RelativeAge
Clock2:Relative Age (N=177)
j
FrontalCortex Cor=0.946
med.Cor=0.946 , MAE=0.012
DNAm RelativeAge
Clock2:Relative Age (N=16)
k
Skin Cor=0.942
med.Cor=0.938 , MAE=0.027
DNAm RelativeAge
Clock2:Relative Age (N=1971)
l
Striatum Cor=0.935
med.Cor=0.935 , MAE=0.046
DNAm RelativeAge
Clock2:Relative Age (N=56)
m
Muscle Cor=0.933
med.Cor=0.818 , MAE=0.045
DNAm RelativeAge
Clock2:Relative Age (N=403)
n
Tail Cor=0.932
med.Cor=0.96 , MAE=0.017
DNAm RelativeAge
Clock2:Relative Age (N=199)
o
Heart Cor=0.926
med.Cor=0.785 , MAE=0.038
DNAm RelativeAge
Clock2:Relative Age (N=324)
p
Ear Cor=0.884
med.Cor=0.905 , MAE=0.015
DNAm RelativeAge
Clock2:Relative Age (N=622)

**Figure S5. Universal clocks predict human mortality risk and tissue specific results.** DNA methylation-based estimates of relative age (y-axis) versus actual relative age (x-axis). The specific tissue or cell type is reported in the title of each panel. Each dot presents a tissue sample colored by tissue and labeled by mammalian species index (Data S1.3-1.4). The analysis is restricted to tissues that have at least 15 samples available. Leave-one-fraction-out cross-validation (LOFO) was used to arrive at unbiased estimates of predictive accuracy measures: median absolute error (MAE) and age correlation based on relative age. "Cor" denotes the Pearson correlation coefficient based on all available samples. "med.Cor" denotes the median values across all species for which at least 15 samples were available. Title is marked in blue if a tissue type was collected from a single species.

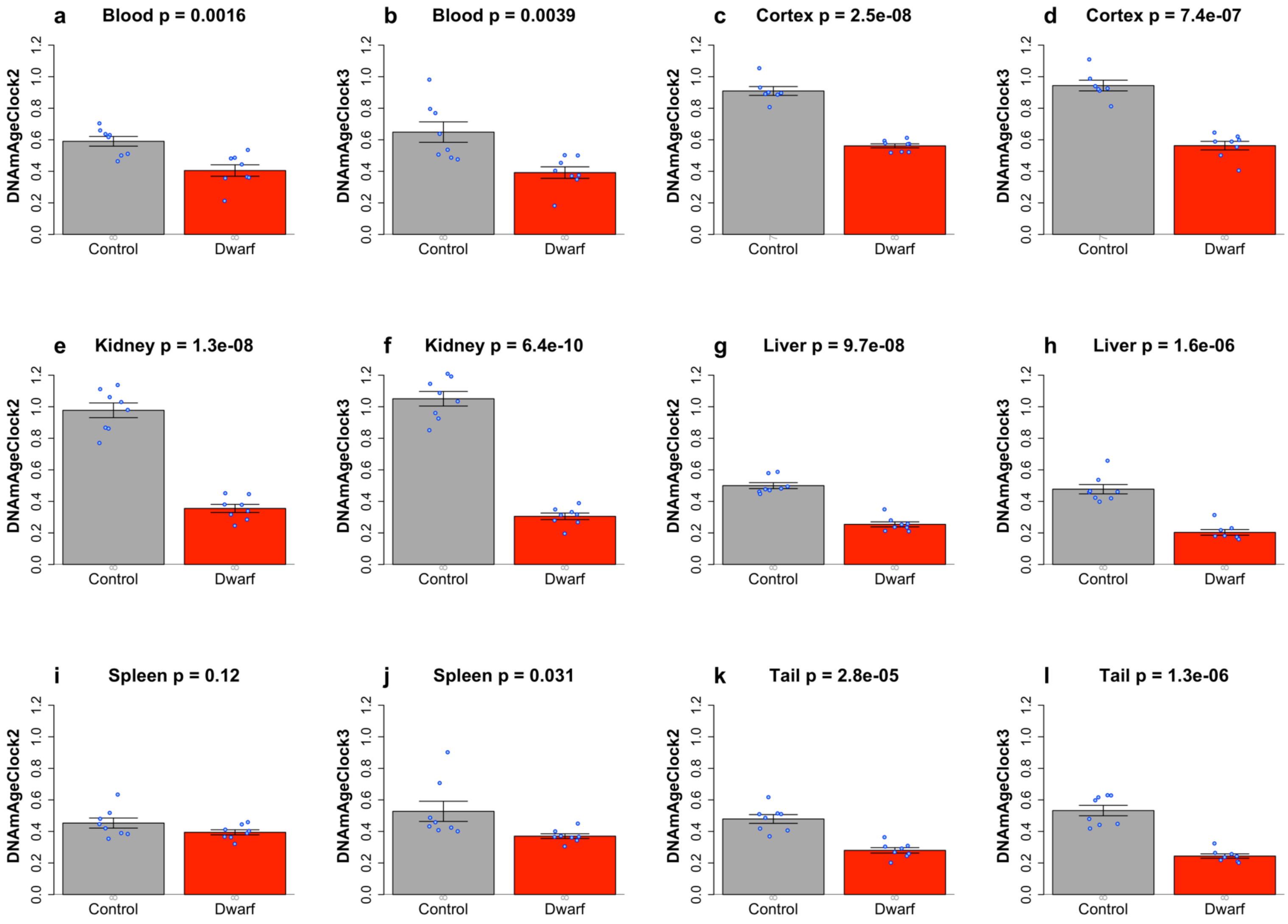


**Figure S6. Snell dwarf mice associated with slower epigenetic aging process based on universal pan-mammalian clocks.** We applied Clock 2 & Clock 3 to evaluate the anti-age intervention based on age-matched Snell dwarf mutation study: 48 normal and 47 Dwarf genotypes with all ages around 0.52 (mean±SD of age =0.52±0.01 years). Each panel presents bar plots for DNA methylation (DNAm) age versus genotype based on Clock 2 (a,c,e &g) and Clock 3 (b,d,f &h), respectively. The blue dots correspond to individual observations. The *y*-axis of the bar plots depicts the mean of one standard error. The numbers of individuals per group are reported as gray numbers underneath each bar. The heading of each panel reports tissue type and a nominal (unadjusted) two-sided *p* value based on t-test. Our analysis shows that Snell dwarf mice exhibit younger DNAm age than wild-type mice in several tissue types especially in kidney and liver samples.

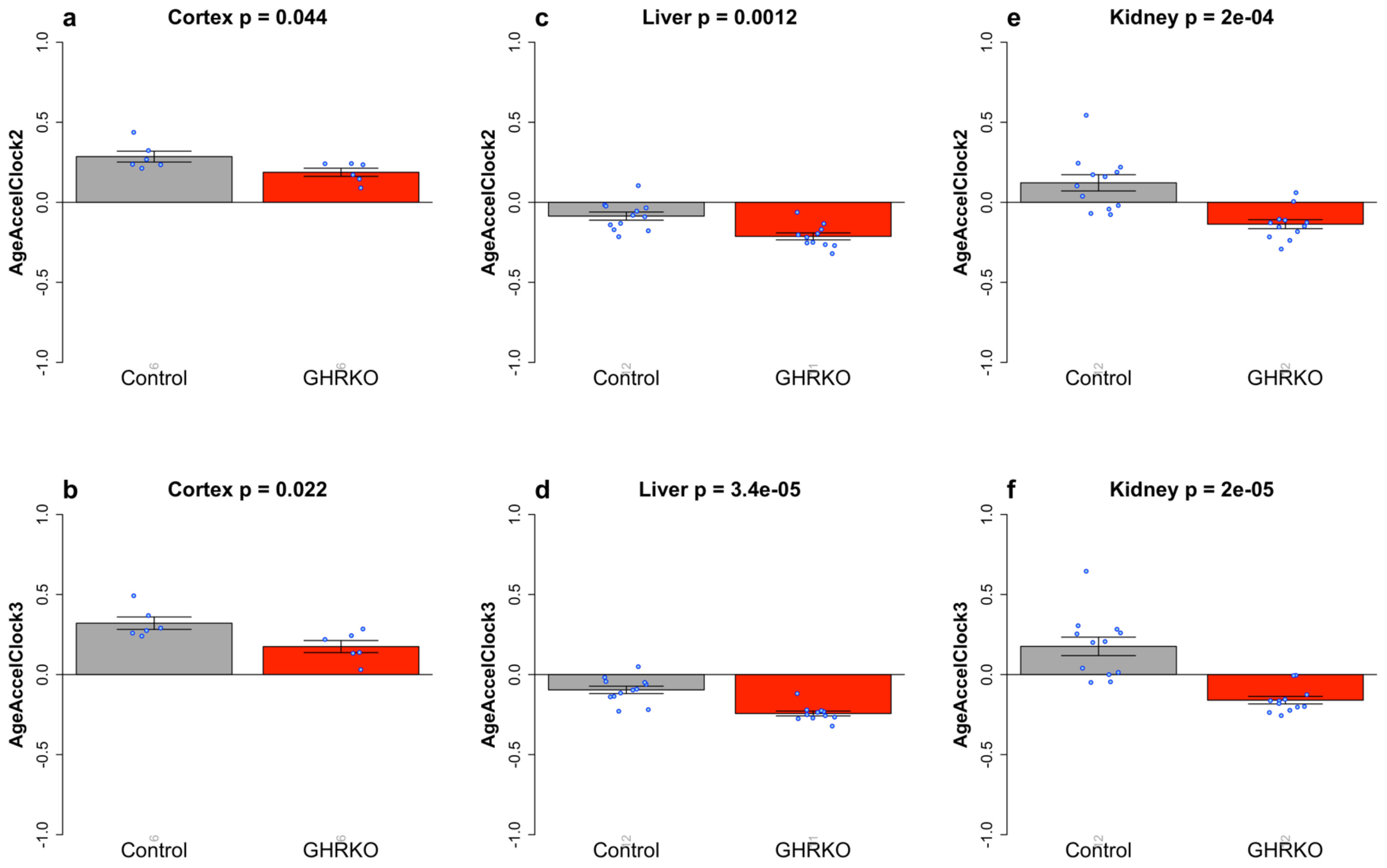


**Figure S7. Full-body growth hormone receptor knock-out mice associated with slower universal pan-mammalian clocks.** We applied Clock 2 & Clock 3 to evaluate the anti-age intervention based on age-matched growth hormone receptor knock-out (GHRKO) experiment 1: 36 normal and 35 GHRKO genotypes with mean±SD of age is 0.65±0.06 years. The comparisons were based on the measures of epigenetic age acceleration, which were defined as raw residual resulting regressing DNAm age on chronological age. By definition, the resulting measure of epigenetic age acceleration (AgeAccel) is not correlated with chronological age. Each panel presents bar plots for epigenetic age acceleration versus genotype based on Clock 2 (the top panels) and Clock 3 (the lower panels), respectively. The blue dots correspond to individual observations. The y-axis of the bar plots depicts the mean of one standard error. The numbers of individuals per group are reported as gray numbers underneath each bar. The heading of each panel reports tissue type and a nominal (unadjusted) two-sided p value based on t-test. Our analysis shows that GHRKO mice are associated with slower epigenetic aging process in several tissue types compared to wild-type mice.

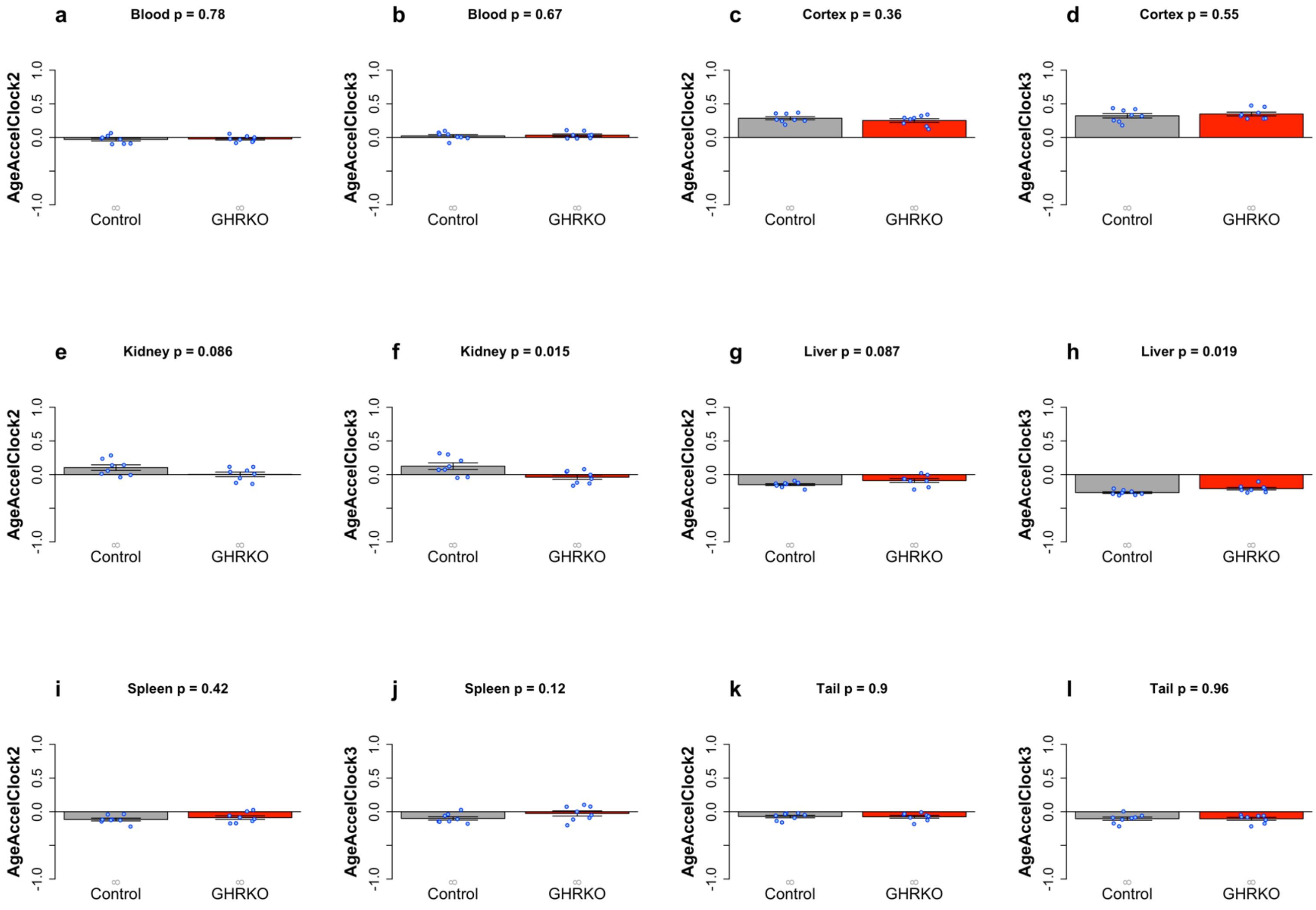


**Figure S8. Evaluation of liver-specific growth hormone receptor knock-out mice based on universal pan-mammalian clocks.** We applied Clock 2 & Clock 3 to evaluate the anti-age intervention based on age-matched GHRKO experiment 2 as a control experiment with knock-out applied on livers only: 48 normal and 48 GHRKO genotypes with mean±SD of age is 0.51±0.03 years old. The comparisons were based on the measures of epigenetic age acceleration, which were defined as raw residual resulting regressing DNAm age on chronological age. By definition, the resulting measure of epigenetic age acceleration (AgeAccel) is not correlated with chronological age. Each panel presents bar plots for epigenetic age acceleration versus genotype based on Clock 2 (a,c,e,g,i & k) and Clock 3 (b,d,f,h,j &l), respectively. The blue dots correspond to individual observations. The y-axis of the bar plots depicts the mean of one standard error. The numbers of individuals per group are reported as gray numbers underneath each bar. The heading of each panel reports tissue type and a nominal (unadjusted) two-sided p value based on t-test. Both Clocks 2 and 3 show that the liver-specific GHRKO mice are not epigenetically younger than the wild type mice with possible exceptions in liver or kidney based on Clock 3. However, the significant P values are not present by Clock 2.

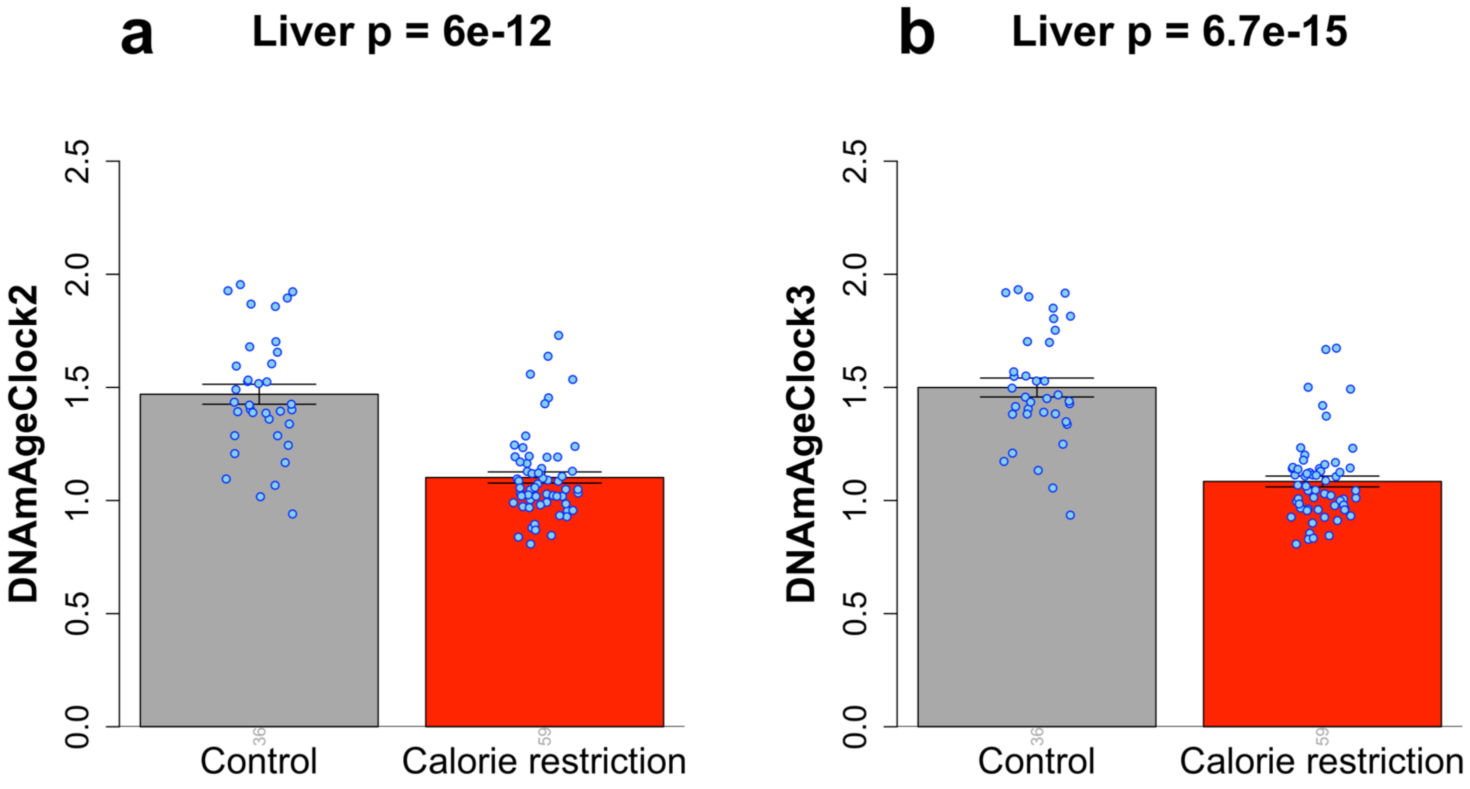


**Figure S9** Calorie restriction resulting in slower epigenetic aging process in mouse livers based on universal pan-mammalian clocks**.** We applied Clock 2 & Clock 3 to evaluate calorie restriction (CR) study in mouse livers: 59 in CR versus 36 control mice with all ages at 1.57 years old. Each panel presents bar plots for DNA methylation (DNAm) age versus experimental group based on Clock 2 (a) and Clock 3 (b). The blue dots correspond to individual observations. The *y*-axis of the bar plots depicts the mean of one standard error. The numbers of individuals per group are reported as gray numbers underneath each bar. The heading of each panel reports tissue type and a nominal (unadjusted) two-sided *p* value based on t-test. Our analysis indicates that CR reduces the rate of epigenetic aging in liver samples, supported by both Clocks 2 and 3.

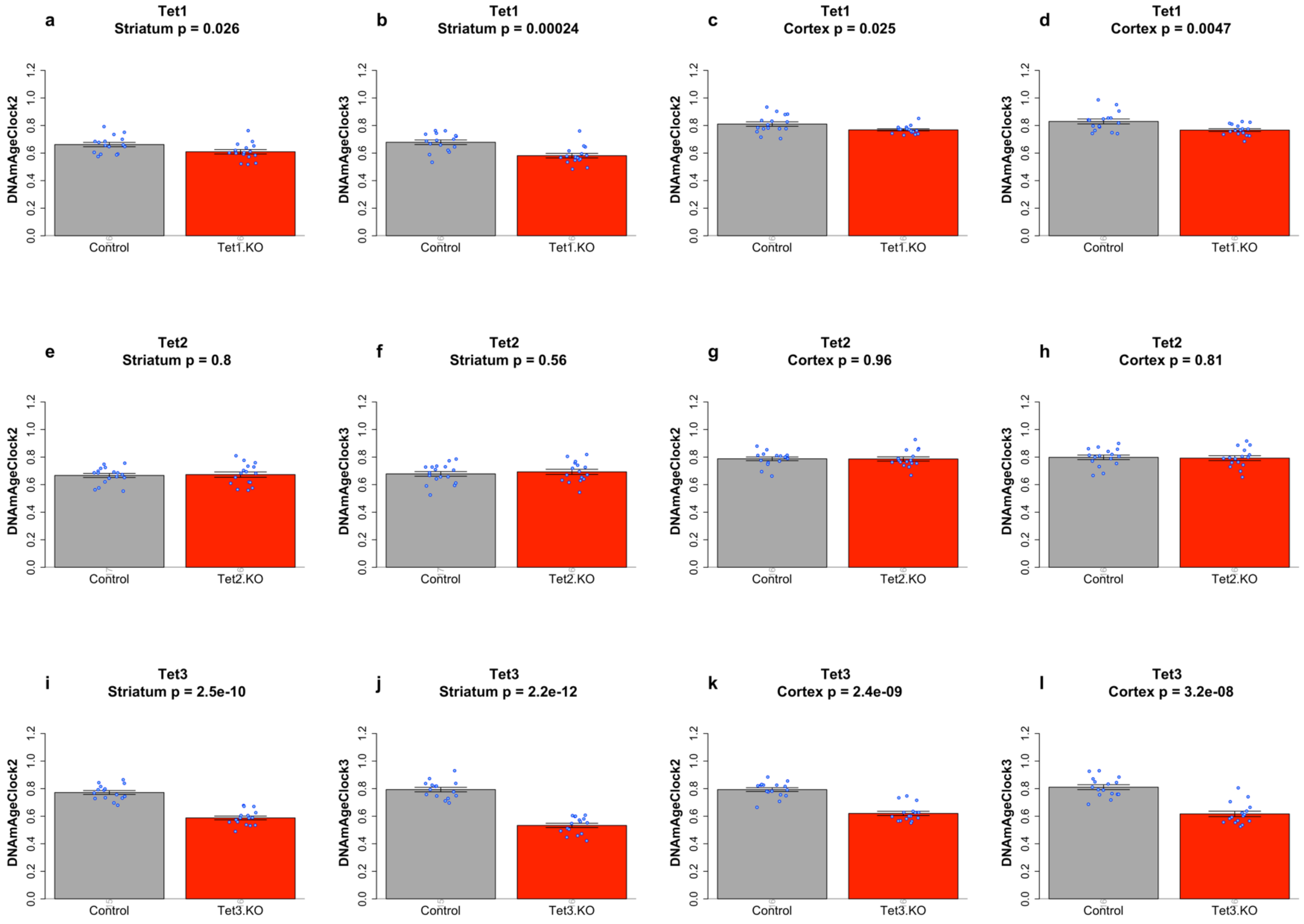


**Figure S10. Tet3 knock-out mice associated with slower epigenetic aging process in brain regions**. We applied Clock 2 & Clock 3 to evaluate Tet gene knout-out study: all mice at age 0.5 years and DNA methylation profiled in cortex and striatum on the same mouse. Tet1 (the top row): 32 controls and 32 Tet1 KO genotypes. Tet2 (the middle row): 33 controls and 32 Tet2 KO genotypes. Tet3 (the bottom row): 31 control and 32 Tet3.KO genotypes. Each panel presents bar plots for DNA methylation (DNAm) age versus genotype based on Clock 2 and Clock 3. The blue dots correspond to individual observations. The *y*-axis of the bar plots depicts the mean of one standard error. The numbers of individuals per group are reported as gray numbers underneath each bar. The heading of each panel reports tissue type and a nominal (unadjusted) two-sided *p* value based on t-test. Our analysis shows that Tet3 KO mice exhibit a reduced rate of epigenetic aging in brain regions. The significant epigenetic age reversal effects were relatively weak for Tet1 KO and barely could be observed for Tet2 KO mice.

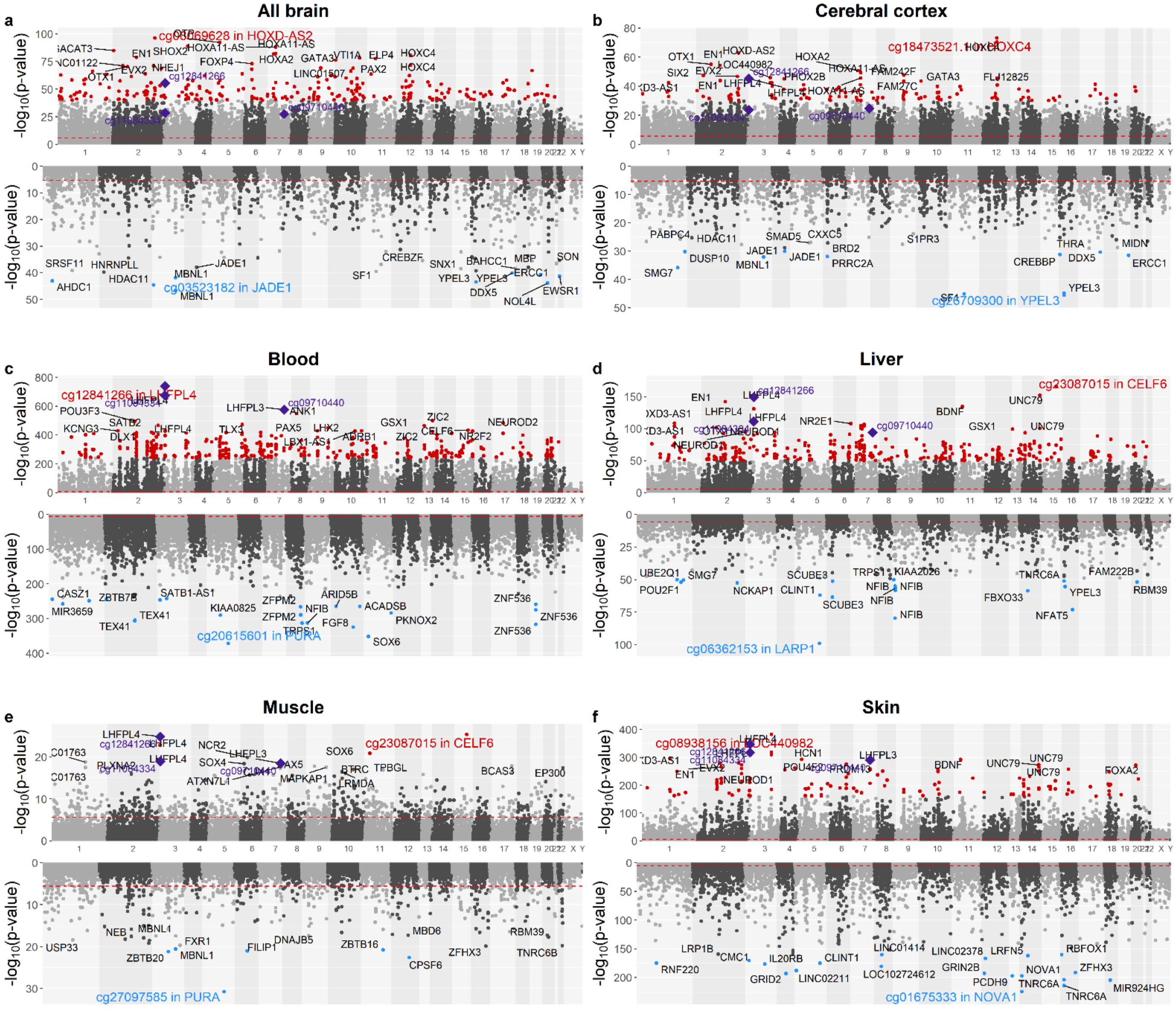


**Figure S11. Meta-analysis of chronological age in mammalian samples across specific tissue types.** Meta-analysis p-value (-log base 10 transformed) versus chromosomal location (x-axis) according to human genome assembly 38 (hg38) in **a**), brain tissues (across multiple brain regions), **b**) cerebral cortex, **c**) blood, **d**) liver, **e**) muscle and **f**) skin tissues. The upper and lower panels of the Manhattan plot depict the CpG sites that gain/lose methylation with age. In panel a, P values were calculated via two-stage meta-analysis that combined EWAS results across strata formed by species/brain-tissue (with n≥15 samples, Methods). CpGs are colored in red and blue if they exhibit highly significant positive and negative age correlations according to a meta analysis P <$1.0x10^{-40}$, $1.0x10^{-30}$, $1.0x10^{-250}$, $1.0x10^{-50}$, $1.0x10^{-20}$ and $1.0x10^{-150}$ for a–f, respectively. Red dashed horizontal lines denote Bonferroni correction. Gene names are annotated for the top 20 CpGs with positive and negative associations, respectively. CpGs are labeled by adjacent genes. Purple color and diamond shapes mark CpGs of particular interest: cg12841266 and cg11084334 in *LHFPL4* and cg09710440 in *LHFPL3.*

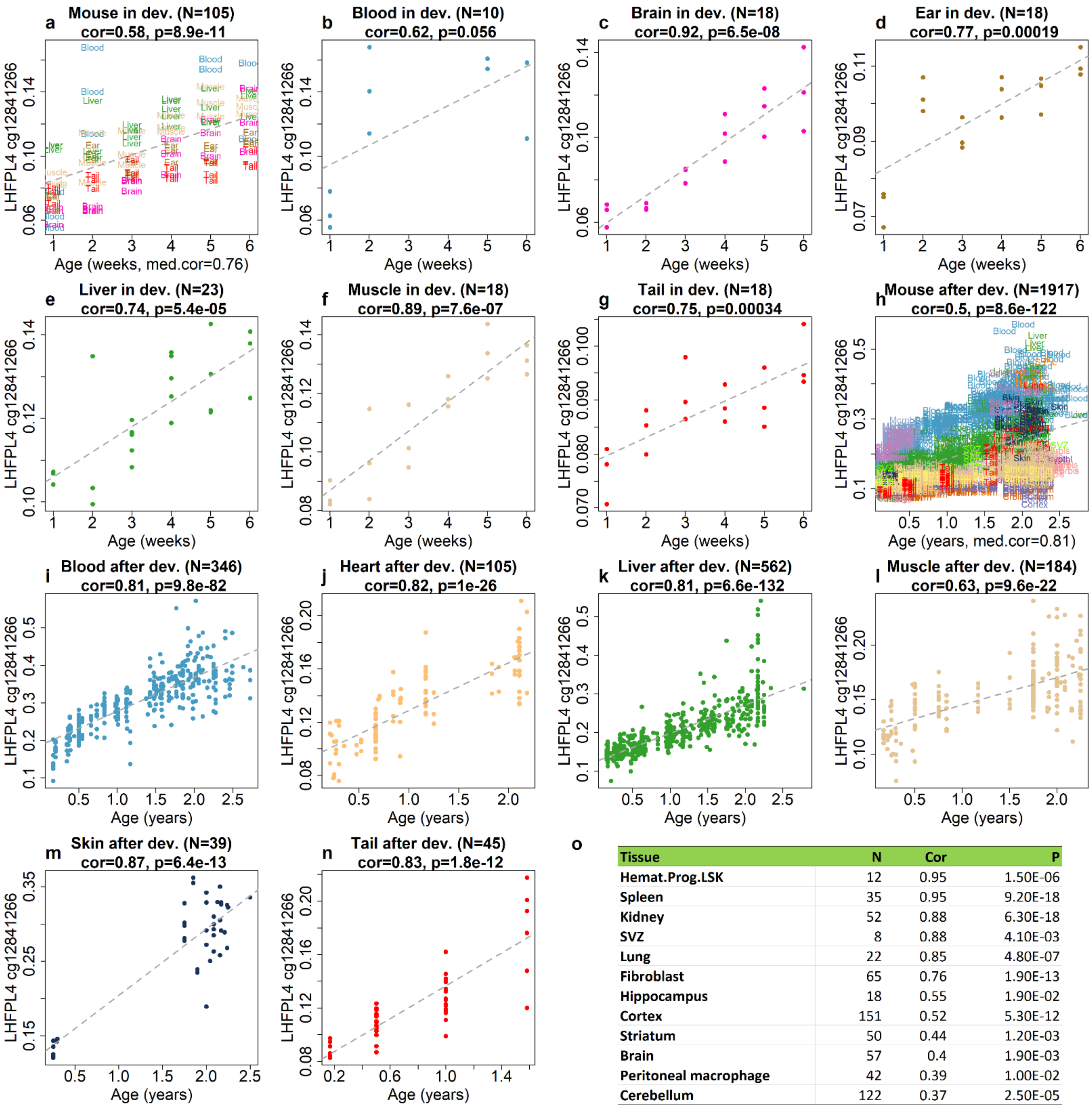


| Tissue | N | Cor | P |
|---|---|---|---|
| Hemat.Prog.LSK | 12 | 0.95 | 1.50E-06 |
| Spleen | 35 | 0.95 | 9.20E-18 |
| Kidney | 52 | 0.88 | 6.30E-18 |
| SVZ | 8 | 0.88 | 4.10E-03 |
| Lung | 22 | 0.85 | 4.80E-07 |
| Fibroblast | 65 | 0.76 | 1.90E-13 |
| Hippocampus | 18 | 0.55 | 1.90E-02 |
| Cortex | 151 | 0.52 | 5.30E-12 |
| Striatum | 50 | 0.44 | 1.20E-03 |
| Brain | 57 | 0.4 | 1.90E-03 |
| Peritoneal macrophage | 42 | 0.39 | 1.00E-02 |
| Cerebellum | 122 | 0.37 | 2.50E-05 |

**Figure S12. Methylation levels of cg12841266 (*LHFPL4)* versus chronological age (x-axis) in mouse tissues.** The panels report pairwise relationships for different tissues and age groups: **a–g**, (age between 1 week and 6 weeks) and **h–o** (older than six weeks). The mean±standard deviation [range] of chronological age is 3.5±1.7 [1.0–6.0] weeks in the developmental age group and 1.12±0.72 [0.15–2.78] years in the post-developmental group. **a, h**, all tissues combined. Each dot (sample) is colored by the tissue type. **b-g**, development and **i-o**, post development. In the panel o, we list the correlation analysis across additional different tissue types. The title reports the Pearson correlation coefficient, a corresponding nominal (unadjusted) two-sided correlation test p-value and the sample size (N).

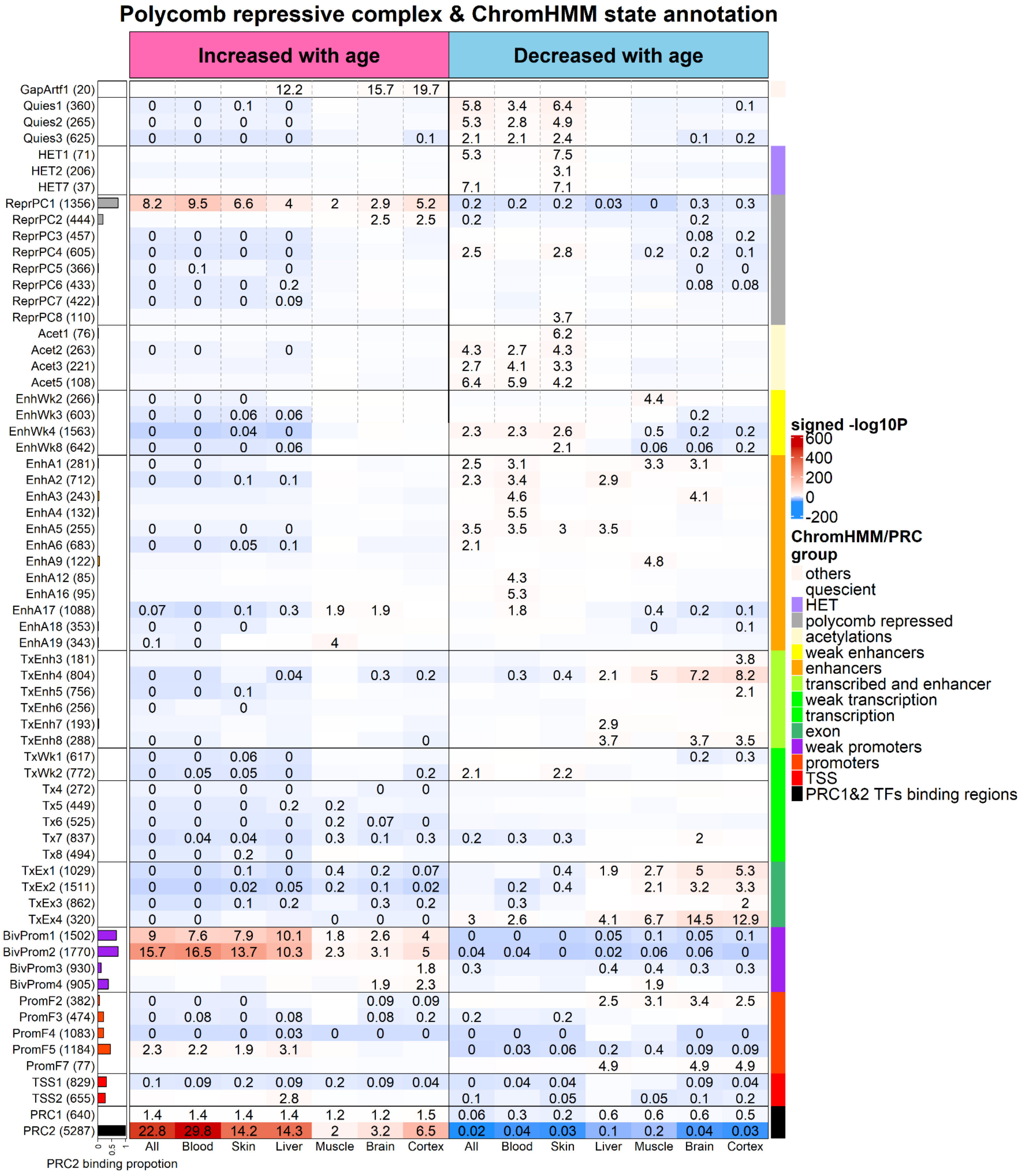

Polycomb repressive complex & ChromHMM state annotation
Increased with age
Decreased with age
GapArtf1 (20)
Quies1 (360)
Quies2 (265)
Quies3 (625)
HET1 (71)
HET2 (206)
HET7 (37)
ReprPC1 (1356)
ReprPC2 (444)
ReprPC3 (457)
ReprPC4 (605)
ReprPC5 (366)
ReprPC6 (433)
ReprPC7 (422)
ReprPC8 (110)
Acet1 (76)
Acet2 (263)
Acet3 (221)
Acet5 (108)
EnhWk2 (266)
EnhWk3 (603)
EnhWk4 (1563)
EnhWk8 (642)
EnhA1 (281)
EnhA2 (712)
EnhA3 (243)
EnhA4 (132)
EnhA5 (255)
EnhA6 (683)
EnhA9 (122)
EnhA12 (85)
EnhA16 (95)
EnhA17 (1088)
EnhA18 (353)
EnhA19 (343)
TxEnh3 (181)
TxEnh4 (804)
TxEnh5 (756)
TxEnh6 (256)
TxEnh7 (193)
TxEnh8 (288)
TxWk1 (617)
TxWk2 (772)
Tx4 (272)
Tx5 (449)
Tx6 (525)
Tx7 (837)
Tx8 (494)
TxEx1 (1029)
TxEx2 (1511)
TxEx3 (862)
TxEx4 (320)
BivProm1 (1502)
BivProm2 (1770)
BivProm3 (930)
BivProm4 (905)
PromF2 (382)
PromF3 (474)
PromF4 (1083)
PromF5 (1184)
PromF7 (77)
TSS1 (829)
TSS2 (655)
PRC1 (640)
PRC2 (5287)
0 0.5 1
PRC2 binding propotion
All Blood Skin Liver Muscle Brain Cortex All Blood Skin Liver Muscle Brain Cortex
signed -log10P
600
400
200
0
-200
ChromHMM/PRC group
others
quescient
HET
polycomb repressed
acetylations
weak enhancers
enhancers
transcribed and enhancer
weak transcription
transcription
exon
weak promoters
promoters
TSS
PRC1&2 TFs binding regions

**Figure S13. Chromatin state analysis of age-related CpGs.** The heatmap color-codes the hypergeometric overlap analysis between age-related CpGs (columns) and two groupings of CpGs (a) universal chromatin states analysis [1] and (b) binding by polycomb repressive complex 1 and 2 (PRC1, PRC2) defined based on ChipSeq datasets in ENCODE[2] , see the last two rows. The first column shows a bar plot that reports the proportion of CpGs that are known to be bounded by PRC2 that ranges from zero (RPC1) to one (PRC2). Note that chromatin states that contain a high proportion of PRC2 bound CpGs overlap significantly with the top 1,000 CpGs that increased with age across tissues and mammal species. For each row (chromatin state or PRC annotation), the table reports odds ratios (OR) from hypergeometric test results for the top 1,000 CpGs that increased/decreased with age from meta-EWAS of age across all, blood, skin, liver, muscle, brain and cerebral cortex tissues, respectively. Hypergeometric P values are listed in **Data S.5.3—S.5.9**. The heatmap color gradient is based on -log10 (unadjusted hypergeometric P value) multiplied by the sign of OR greater than one. Red colors denote OR greater than one in contrast with blue colors for OR less than one. Legend lists states based on their group category and PRC group. The y-axis lists state or PRC name and number of mammalian array CpGs inside parentheses. The left/right panel lists the results based on the top 1,000 CpGs with positive/negative age correlation. We displayed 63 universal chromatin states that show significant enrichment/depletion at $P < 0.001$ in any of the tissues.

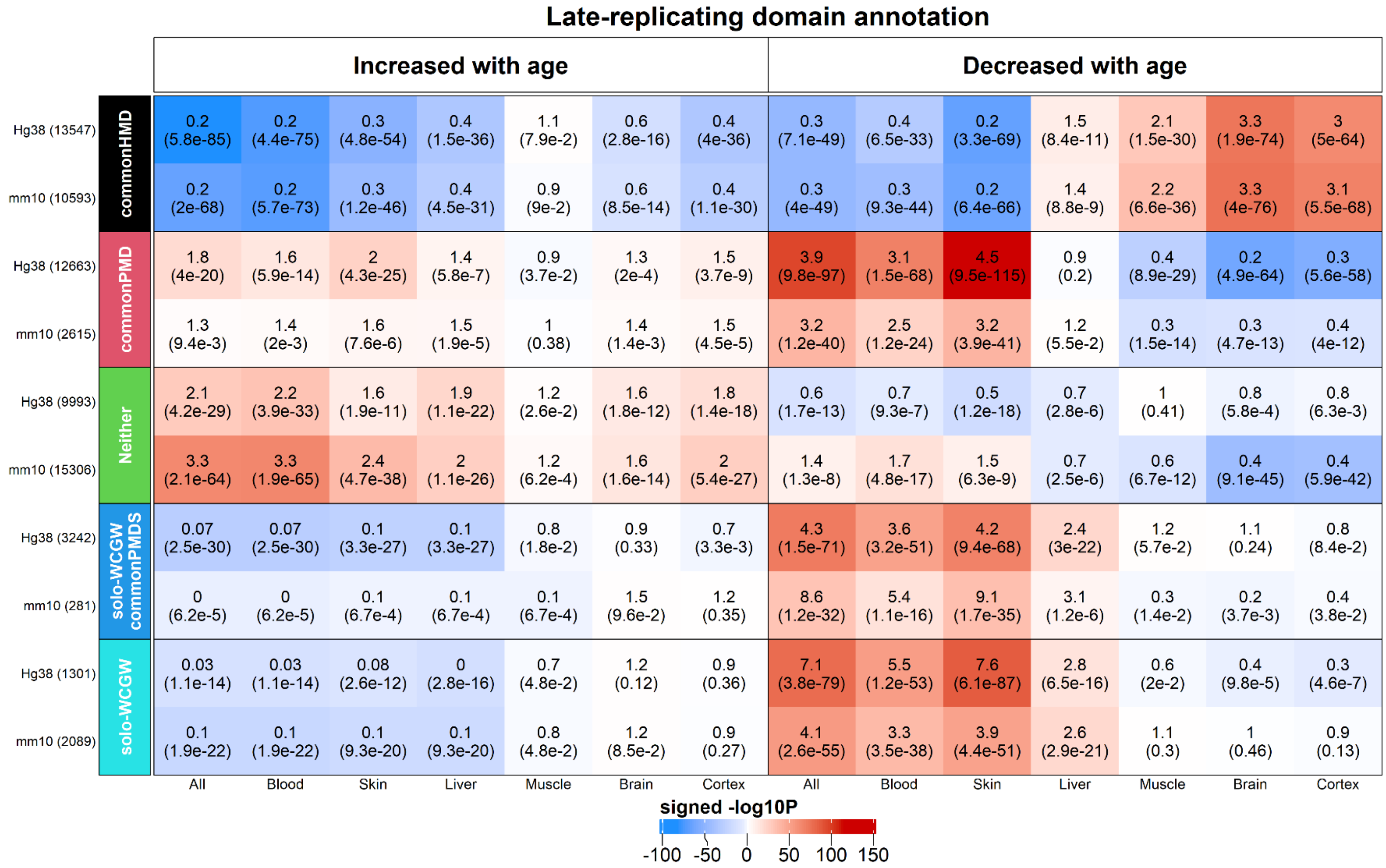


**Figure S14. Overlap with late-replicating domains.** The heatmap color-codes the hypergeometric overlap analysis between age-related CpGs (columns) and CpGs related to late-replicating domains in Hg38 and mm10 assembly[3], respectively. Two groups of late-replicating domains were analyzed (a) common PMD/HMD structures: highly methylated domains (commonHMD), partially methylated domains (commonPMD), and neither (Neither), and (b) solo-WCGW structures: genome-wide (solo-WCGW) and those in the common PMD regions (solo-WCGW commonPMDs). The y-axis lists categories of late-replicating domains and number of mammalian array CpGs inside parentheses for Hg38 and mm10 genome, respectively. For each row, the table reports odds ratios (OR) from hypergeometric test results for the top 1,000 CpGs that increased/decreased with age from meta-EWAS of age across all, blood, skin, liver, muscle, brain and cerebral cortex tissues, respectively. The heatmap color gradient is based on -log10 (unadjusted hypergeometric P value) multiplied by the sign of OR greater than one. Red colors denote OR greater than one in contrast with blue colors for OR less than one. The left/right panel lists the results based on the top 1,000 CpGs with positive/negative age correlation.

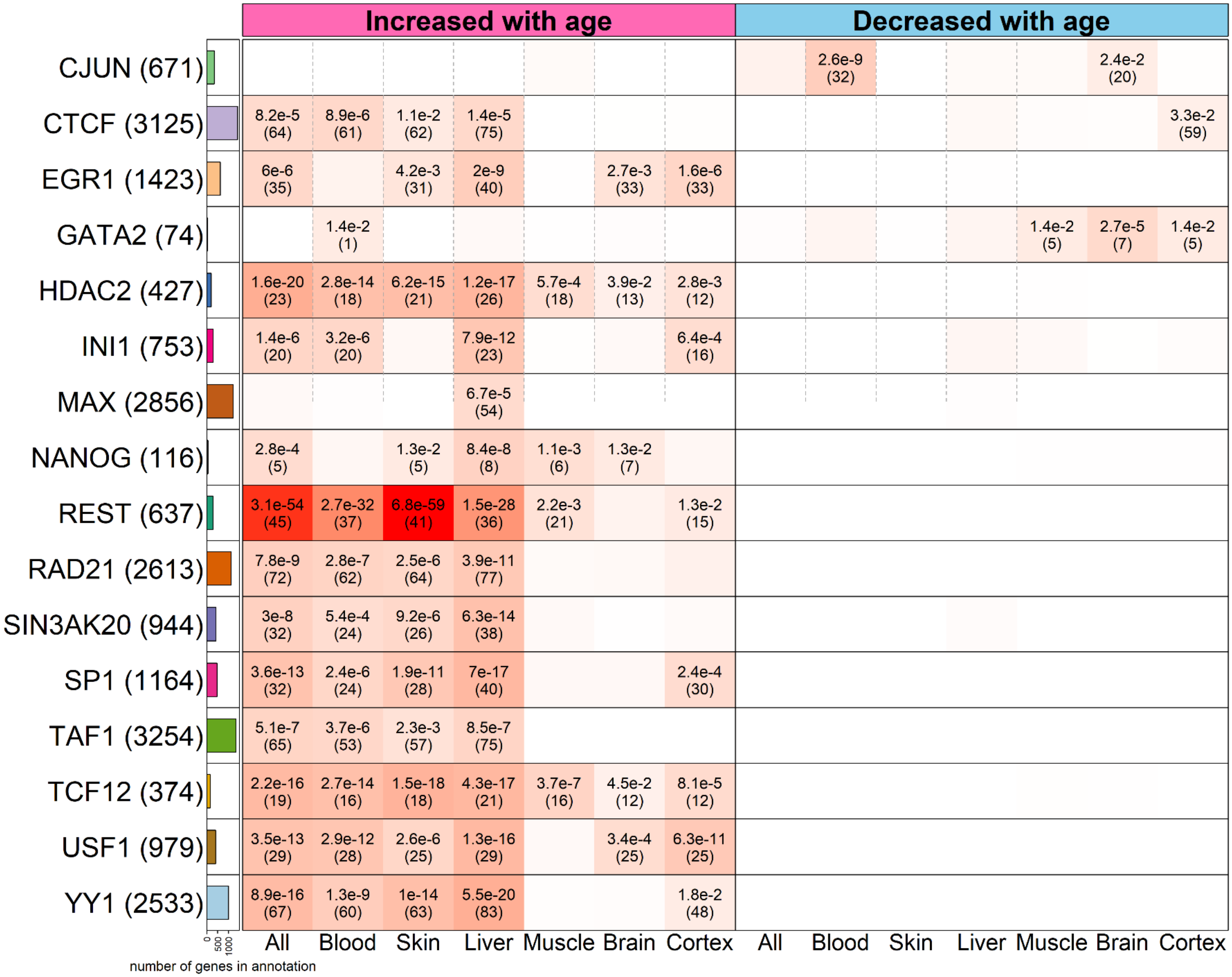


**Figure S15. Enrichment with Transcription factor binding regions**. We studied the overlapping genomic regions between (1) the CpG sites located in the binding regions of 68 transcription factors (TF) in hg19 and (2) the top 1000 CpGs that increased/decreased with age from EWAS of age across mammalian tissues. TF results (y-axis, rows) versus mammalian EWAS of age are stratified by tissue type (x-axis, columns). The left/right panels of the x-axis list the top 1000 CpGs that increased/decreased with age from meta-EWAS of age across all, blood, skin, liver, muscle, brain and cerebral cortex tissues, respectively. The y-axis lists the names of transcription factors and number of mammalian array CpGs located in the binding sites. Background in hypergeometric tests was based on the genes present in our mammalian array. The bar plots in the first column report the total number of genes at each TF according to the background. The heatmap color codes -log10 (unadjusted hypergeometric P value). Hypergeometric P values (number of overlapping genes) are listed on the heatmap provided $P<0.05$.

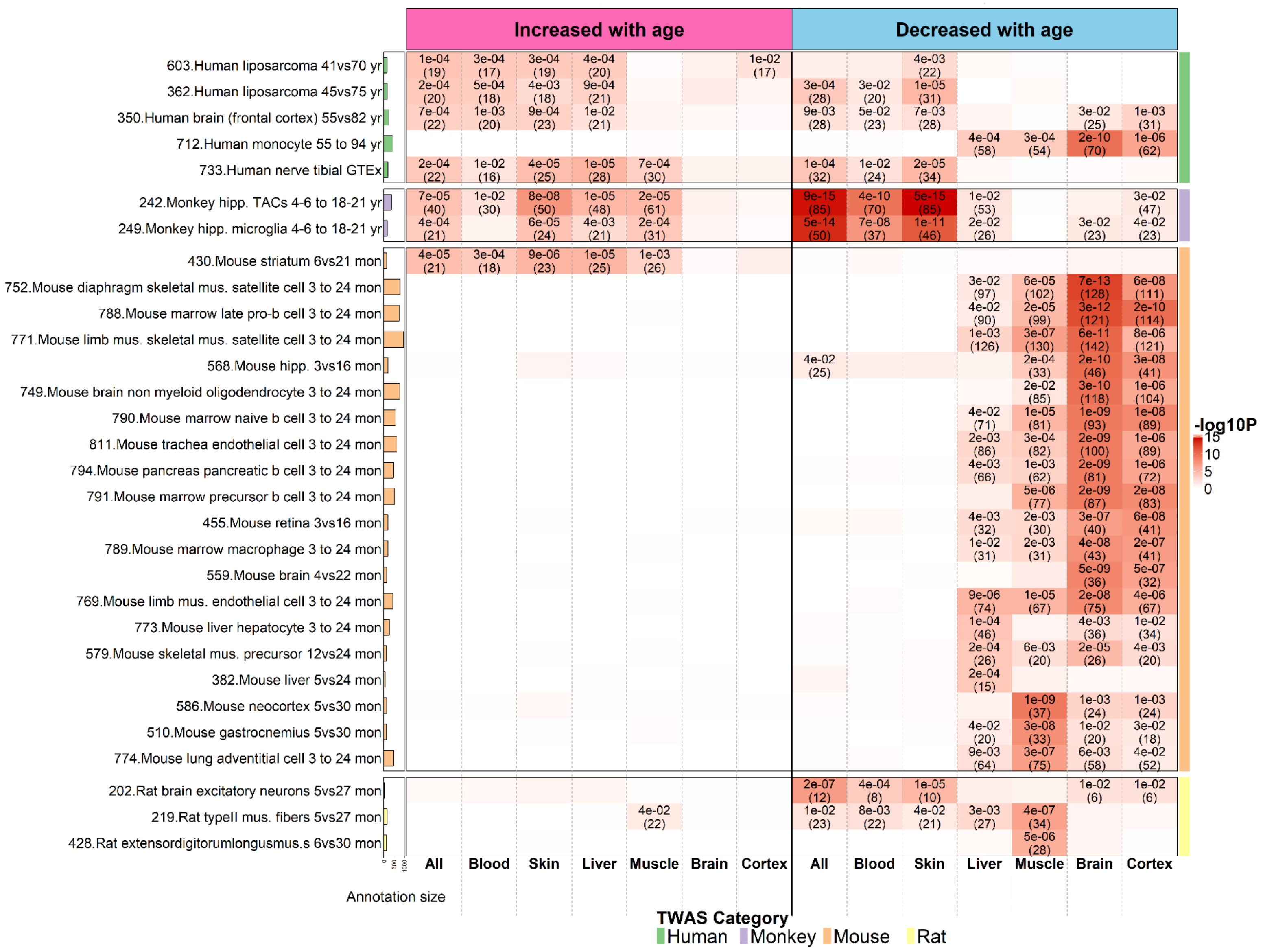


**Figure S16. Enrichment with EWAS-TWAS associated genes**. As detailed in Methods, we studied the overlapping genes between (1) the genes implicated in transcriptome-wide association studies (TWAS) in different cell types or species and (2) the top 1000 CpGs that increased/decreased with age from EWAS of age across mammalian tissues. TWAS results for complex traits (y-axis, rows) versus mammalian EWAS of age are stratified by tissue type (x-axis, columns). The left/right panels of the x-axis list the top 1000 CpGs that increased/decreased with age from meta-EWAS of age across all, blood, skin, liver, muscle, brain and cerebral cortex tissues, respectively. The y-axis lists TWAS index number (listed in Supplementary Data) followed by selected aging traits in human, monkey, mouse and rat species. Background in hypergeometric tests was based on the genes present in our mammalian array. The bar plots in the first column report the total number of genes at each TWAS according to the background. The heatmap color codes -log10 (unadjusted hypergeometric P value). The color band on the right-side annotation encodes species category in our TWAS collections. Hypergeometric P values (number of overlapping genes) are listed on the heatmap provided *P*<0.05. Abbreviations: hipp.=hippocampus, MPNST= malignant peripheral nerve sheath tumor, mus.=muscle, TACs= transiently amplifying progenitor cells.

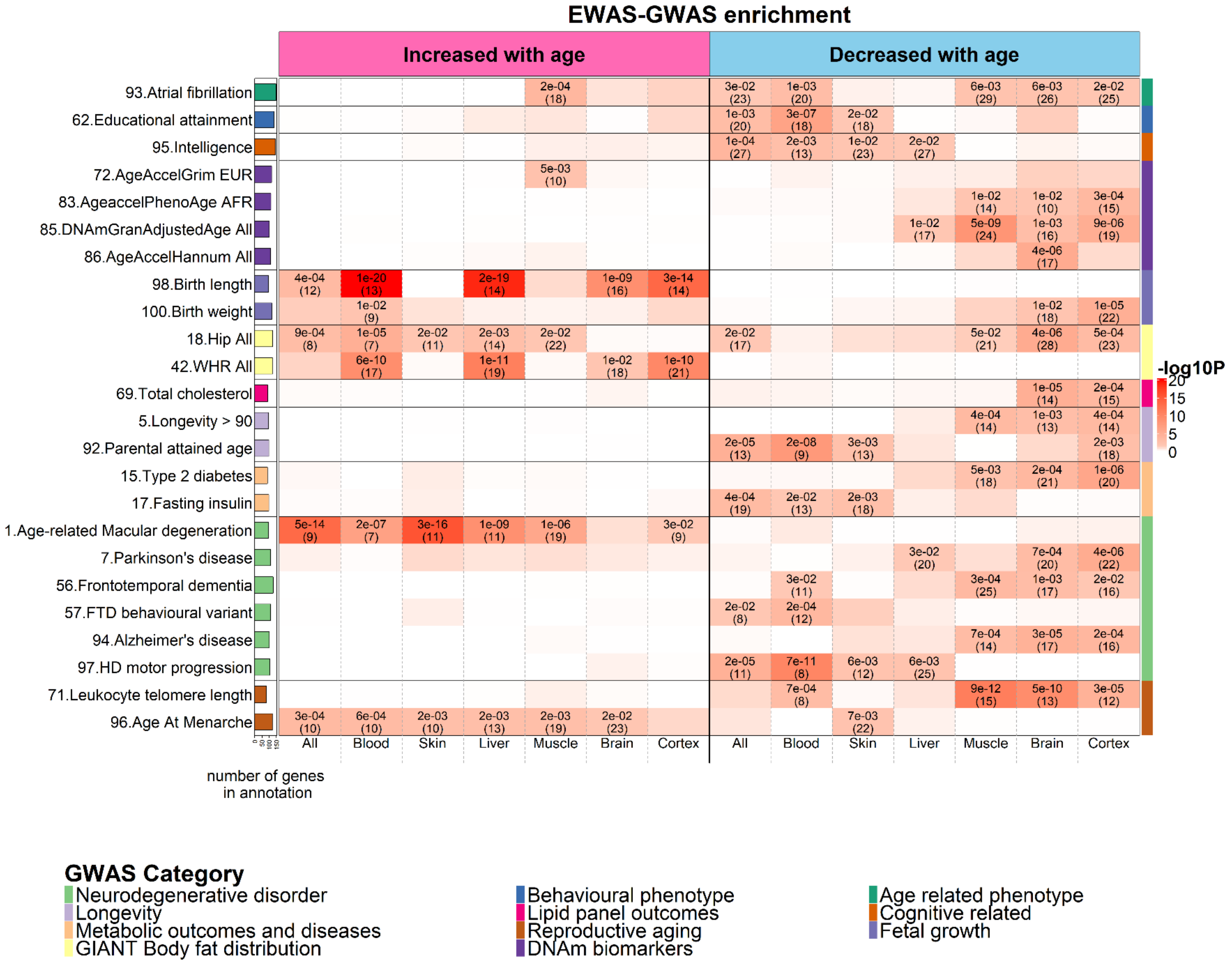


**Figure S17. Enrichment with EWAS-GWAS associated genes**. As detailed in Methods, we studied the overlapping genomic regions between (1) the top 2.5% genes implicated by genome-wide association studies (GWAS for SNP associations) of human complex traits and (2) the top 1,000 CpGs that increased/decreased with age from EWAS across mammalian tissues. GWAS results for complex traits (y-axis, rows) versus mammalian EWAS of age are stratified by tissue type (x-axis, columns). The left/right panels of the x-axis list the top 1,000 CpGs that increased/decreased with age from meta-EWAS of age across all, blood, skin, liver, muscle, brain and cerebral cortex tissues, respectively. The y-axis lists GWAS index number (listed in Supplementary Data) followed by selected complex traits. Background in hypergeometric tests was based on the genomic regions present in our mammalian array. The bar plots in the first column report the total number of the top 2.5% genes at each GWAS according to the background. The heatmap color codes -log10 (unadjusted hypergeometric P value). The color band on the right-side annotation encodes the GWAS category. Hypergeometric P values (number of the overlapping genomic regions) are listed on the heatmap provided *P*<0.05. Abbreviations: All=All ancestries, EUR=European ancestry, AFR=African American ancestry, FTD= frontotemporal dementia, WHR= waist to hip ratio.

#### Supplementary Tables

| Rank | Chr | Gene | CpG | Protein domain | Activity | Function |
|---|---|---|---|---|---|---|
| **Increased with age** | | | | | | |
| 1 | 3 | LHFPL4 | cg12841266 | Tetraspan transmembrane | Clustering of GABA receptors | Dev. of inhibitory synapse |
| 2 | 3 | ZIC1 | cg08938156 | Zinc finger | TF | Early Dev. |
| 3 | 7 | LHFPL3 | cg09710440 | Tetraspan transmembrane | Clustering of GABA receptors | Dev. of inhibitory synapse |
| 4 | 5 | TLX3 | cg26844246 | Homeobox | TF | Dev. |
| 5 | 2 | EVX2 | cg09227056 | Homeobox | TF | Limb Dev. |
| 6 | 13 | ZIC2 | cg15682828 | Zinc finger | transcriptional activator or repressor | important in the early stage of organogenesis of the CNS |
| 7 | 1 | FOXD3 | cg05551621 | Forkhead | TF | Embryonic Dev. |
| 8 | 15 | CELF6 | cg23087015 | RNA recognition motif (RRM) domains | RNA-binding | RNA processing |
| 9 | 17 | NEUROD2 | cg03679521 | bHLH | TF | Neural Dev. |
| 10 | 11 | BDNF | cg27201382 | neurotrophin family of growth factors | -- | Brain Dev. |
| 11 | 10 | PAX2 | cg01486146 | Paired box | TF | development of the urogenital tract, the eyes, and the CNS |
| 12 | 9 | PAX5 | cg20766695 | Paired box | TF | Early Dev. |
| 13 | 5 | OTP | cg24352905 | Homeobox | TF | Dev. / Cell fate specification |
| 14 | 16 | SALL1 | cg13909487 | Zinc finger | TF | Stem cell regulation and Dev. |
| 15 | 6 | NRN1 | cg16356803 | neuritin family | -- | Promotes neurite outgrowth |
| 16 | 6 | PRDM13 | cg05797975 | PR Domain Zinc Finger | chromatin binding *and* methyltra nsferase activity | transcriptional regulation |
| 17 | 6 | NR2E1 | cg27496468 | Zinc finger | Nuclear hormone receptor | Retinal Dev. |
| 18 | 2 | NEUROD1 | cg09942248 | bHLH | TF | Neural Dev. |
| 19 | 6 | TBX18 | cg09817427 | T-Box | TF | embryonic development of the sino atrial node head area |
| 20 | 20 | NKX2-2 | cg16524928 | Homeobox | TF | Spinal cord Dev. |
| 21 | 6 | TFAP2D | cg26834803 | -- | TF | Embryonic Dev. |
| 22 | 2 | SIX2 | cg03557129 | Homeobox | TF | Limb & Eye Dev. |
| 23 | 7 | TWIST1 | cg20477718 | bHLH | TF | Embryonic Dev. |

| | | | | | | |
|---|---|---|---|---|---|---|
| 24 | 4 | PHOX2B | cg13278722 | Homeobox | TF | Neural Dev. |
| 25 | 13 | OBI1-AS1 | cg13739934 | -- | Anti-sense RNA | -- |
| 26 | 11 | DBX1 | cg21452781 | Homeobox | TF | establishing the distinction of V0 and V1 neuronal fate |
| 27 | 13 | ZIC5 | cg24255409 | Zinc finger | TF | Early Dev. |
| 28 | 2 | OTX1 | cg03957108 | Homeobox | TF | Brain and sensory Dev. |
| 29 | 5 | LOC642366 | cg16744551 | -- | Anti-sense RNA | -- |
| 30 | 14 | VSX2 | cg10707128 | Homeobox | retina-specific TF | specification and morphogenesis of the sensory retina |
| 31 | 7 | HOXA13 | cg26512254 | Homeobox | TF | embryonic development |
| 32 | 7 | DLX6-AS1 | cg02898094 | -- | Anti-sense RNA | Dev. |
| 33 | 8 | EGR3 | cg02040024 | Zinc finger | Transcription Regulator | Numerous Dev. processes |
| 34 | 6 | POU3F2 | cg08921975 | Homeobox | TF | Nueron Dev. |
| 35 | 5 | IRX1 | cg02926165 | Homeobox | TF | Pattern formation |
| **decreased with age** | | | | | | |
| 1 | 5 | LARP1 | cg12880090 | regulating translation of mTORC1 downstream targets | RNA-binding | Regulation of translation |
| 2 | 21 | SON | cg24466972 | -- | RNA-binding | RNA splicing |
| 3 | 15 | SNX1 | cg27626343 | Phox (PX) | Endosomal protein | Regulates expression of EGF receptor |

**Table S1. Top age-related CpGs consistently found by EWAS in different tissues of mammals.** The table provides details on CpGs and genes that are in the intersection (Venn diagram) of EWAS of age in different tissues across different mammalian species (see **Fig. 1g**). For brain, blood, liver, muscle and skin we identified the top 1,000 *positively* age-related CpGs for each tissue type. The five sets of CpGs shared 48 CpGs located near 35 genes (increased with age). We only report the most significant CpG when several CpGs implicated the same gene. Conversely, the top 1,000 *negatively* age-related CpGs in the five tissue types implicated only three genes (decreased with age). We arrange the list of CpGs based on the EWAS meta P-value from all tissues, stratified by the sign or EWAS meta Z score. Columns report the rank, chromosome of the gene (CHR), HGNC gene symbol, the most significant CpG, protein domain, activity and function of gene expression. Abbreviations: Dev=development; TF=transcriptional factor, CNS= central nervous system.

| Universal Clock | Validation scheme | DNAm Age vs age | | | | DNAm relative age vs relative age | | |
|---|---|---|---|---|---|---|---|---|
| | | Cor | MAE | med. Cor | med. MAE | Cor | MAE | med. MAE |
| **Clock 1:** Basic, log(Age+2) | LOFO | 0.963 | 0.757 | 0.880 | 0.962 | 0.884 | 0.034 | 0.033 |
| **Clock 2:** RelativeAge | LOFO | 0.983 | 0.562 | 0.925 | 0.676 | 0.950 | 0.024 | 0.026 |
| **Clock 3:** log-linear age | LOFO | 0.980 | 0.550 | 0.918 | 0.743 | 0.942 | 0.024 | 0.029 |
| **Clock 1:** Basic, log(Age+2) | LOSO | 0.828 | 1.855 | 0.862 | 1.725 | 0.455 | 0.080 | 0.059 |
| **Clock 2:** RelativeAge | LOSO | 0.941 | 1.439 | 0.881 | 1.428 | 0.786 | 0.060 | 0.049 |
| **Clock 3:** log-linear age | LOSO | 0.915 | 1.416 | 0.880 | 1.661 | 0.677 | 0.059 | 0.056 |

**Table S2. Predictive accuracy of three universal mammalian clocks.**

The rows correspond to three universal mammalian clocks for 1) log-transformed age, log(Age+2), 2) relative age, and 3) log-linear age. The inverse age transformations were used to arrive at age estimates, DNA methylation age (in units of years). Each clock was evaluated with two validation schemes: the first three rows report results for leave-one-fraction-out cross validation (LOFO). The last three rows report results for leave-one-species-out cross validation (LOSO). Columns 3 and 4 report the Pearson correlation coefficient (Cor) and median absolute error (MAE, in units of years) across all samples, i.e. species and tissue type are being ignored. Columns 5 and 6 report the median value across species, i.e. median Cor (med.Cor) and median MAE (med.MAE), for species with at least 15 observations. Columns 7–9 repeated the analysis based on relative age. Since the med.Cor for relative age is the same as med Cor for age, we omitted the former. In Clocks 1 and 3, DNAm relative age was computed based on formula (1) with the variable Age replaced by DNAm Age.

### Supplementary Note

#### Supplementary Note 1: Data from the Mammalian Methylation Consortium

The combined data was generated by the Mammalian Methylation Consortium. DNA was extracted using the DNeasy Blood & Tissue Kit from Qiagen. Nucleic acid purity was inspected with a NanoDrop spectrophotometer, and quantified using a Qubit fluorometer dsDNA BR Assay.

Below, we list the descriptions of species.

**Primates [4]**

**Ethics**

This research complied with all relevant ethical regulations overseen by seven ethics review boards. The human skin samples were acquired with informed consent prior to collection of human skin samples with approved by the Oxford Research Ethics Committee in the UK; reference 10/H0605/1. Participants were not compensated. The secondary use of the other de-identified/coded human tissue samples (blood, postmortem tissues) is not interpreted as human subjects research under U.S. Department of Health & Human Services 45 CFR 46. Therefore, the need to obtain written, informed consent from human study participants was waived (secondary use of de-identified tissues). Human samples were covered by University of California Los Angeles IRB#18-000315. All procedures related to non-human primates were approved by different committees: baboons (UTHSCSA Animal Care and Use Committee), strepsirrhini (Duke Institutional Animal Care and Use Committee and the DLC Research Committee), rhesus macaques (Animal Care and Use Committee of the NIA Intramural Program) [5], vervet monkey (UCLA and VA Institutional Animal Care and Use Committees) [6], marmosets (IACUC of UTHSA) [7].

**Baboon care and maintenance**

All animals were given a full veterinary examination prior to recruitment to the study and no obvious cause of ill health or pathology was observed. The animals were housed in group cages at the Southwest National Primate Research Center, at Texas Biomedical Research Institute (TBRI), in San Antonio, Texas in mixed sex groups of up to 16. The remaining 4 females were housed in individual cages at the UT Health Sciences Center San Antonio (UTHSCSA).

Twenty-eight females and the ten males were fed ad libitum Purina Monkey Diet 5038 (12% energy from fat, 0.29% from glucose and 0.32% from fructose and metabolizable energy content of 3.07 kcal/g protein; Purina LabDiets, St Louis, MO, USA) (CTR). Water was continuously available to all animals. Animal health status was recorded daily.

**Necropsy**

None of the animals were euthanized for this project. Rather, we used left-over frozen tissue samples that had previously been collected as part of other projects. Necropsies were performed by either a qualified, experienced veterinarian or M.D investigator. At TRBI, baboons were pre-medicated with ketamine hydrochloride (10 mg/kg IM) and anesthetized using isoflurane (2%) resulting in general anesthesia as previously described [8]. Baboons were exsanguinated while under general anesthesia as approved by the American Veterinary Medical Association. At UTHSCSA four animals were euthanized using Pentobarbital at 390 mg/ml (Fatal- Plus Solution, Vortech, Dearborn, MI, USA) . Following cardiac asystole, respiratory failure and a lack of reflexive response to both skin pinch and eye touch stimulation, tissues (adipose, cerebellum, cerebral cortex, muscle, heart, liver) were rapidly dissected and immediately frozen in liquid nitrogen.

For the studies in which fetal tissue was obtained, all animals were housed in 20 foot × 20 foot × 15 foot metal and concrete group cages at the Texas Biomedical Research Institute. Experimental animals were obtained from appropriate groups of 16 healthy female baboons of similar pre-study body weights (10–15 kg) and morphometric features. The potential day of conception was determined based on the day of ovulation and changes in sex skin color and pregnancy was confirmed at 30 days post ovulation by using ultrasonography. Details of housing, feeding, and environmental enrichment have been published elsewhere. All procedures were approved by the University of Texas Health Science Center and Texas Biomedical Research Institute internal animal care and use committees and performed in the Association for Assessment and Accreditation of Laboratory Animal Care–approved facilities.

Prior to Cesarean section, baboons were premedicated with ketamine hydrochloride (10 mg/kg, IM). Following tracheal intubation, isoflurane (2%, 2L/min, by inhalation) was used to maintain an appropriate plane of anesthesia throughout the surgery. A cesarean section was performed at gestational day 165 (0.9 of gestation) using standard sterile techniques as previously described [9]. Following hysterotomy, the umbilical cord was identified and used for fetal exsanguination with both maternal and fetal baboon under general anesthesia as approved by the American Veterinary Medical Association Panel on Euthanasia. Postoperatively, mothers were placed in individual cages and watched until they were upright under their own power. Maternal analgesia was administered for 3 days (buprenorphine hydrochloride injection; Hospira, Inc., Lake Forest, IL, USA; 0.015 mg/kg/day) post-operatively or longer if indicated. They were returned to their group cage two weeks postoperatively.

Animals were individually fed to enable precise regulation of intake either between 7:00 am and 9:00 am or 11:00 am and 1:00 pm as described in detail elsewhere [10]. Water was continuously available in each feeding cage (Lixit, Napa, California), and the animals were fed Purina Monkey Diet 5038 (Purina, St Louis, Missouri). For this study, we selected samples representing the entire primate lifespan, from neonate to old age.

**Strepsirrhine primates**

Strepsirrhini is a suborder of primates that includes the lemuriform primates, which consist of the lemurs of Madagascar, pottos and galagos from Africa, and the lorises from Southeast Asia. Lemuroids and lorisoids together form the more ancestral sister clade to all other living primates. As such, they lend unparalleled power to any comparative study within the primate clade.

For this study, we selected a total of 91 samples from individuals representing 26 strepsirrhine species, in most cases, the entire lifespan, from immature (infant or juvenile) to senile stages: 68 samples from peripheral blood, 23 samples from skin. The strepsirrhine primates (suborders Lemuriformes and Lorisiformes) used in this study were from the Duke Lemur Center (DLC) in Durham, NC (USA). The Duke Lemur Center is certified by both the Association for Assessment and Accreditation of Laboratory Animal Care and the American Zoological Association. The animal handling and sample collection procedures in this study were performed by a veterinarian after review and approval by the Duke Institutional Animal Care and Use Committee and the DLC Research Committee. Both housing and sample collection met or exceeded all standards of the Public Health Service's “Policy on the Humane Care and Use of Laboratory Animals”. The lemurs are housed in comparable social and housing conditions, habituated to human presence, and individually identifiable. The DLC also maintains a large collection of banked tissues, deriving from routine veterinary procedures and necropsies, amassed over the Center’s 55-year history. Detailed records of life and medical history, reproduction, and social-group membership are digitally maintained [11].

Peripheral blood was collected through venipuncture with standard procedures, either during a routine veterinary procedure or at time of necropsy. Skin tissues were collected during necropsies. Whole blood was preserved in either EDTA or Lithium Heparin, and stored at -80oC. Skin tissues were either frozen directly at -80oC or were first flash frozen and then stored at -80oC.

We profiled the following species: Cheirogaleus medius (Fat-tailed dwarf lemur), Daubentonia madagascariensis (Aye-aye), Eulemur albifrons (White-headed lemur), Eulemur collaris (Collared brown lemur), Eulemur coronatus (Crowned lemur), Eulemur flavifrons (Blue-eyed black lemur), Eulemur fulvus (Brown lemur), Eulemur macaco (Black lemur), Eulemur mongoz (Mongoose lemur), Eulemur rubriventer (Red-bellied lemur), Eulemur rufus (Red-fronted lemur), Eulemur sanfordi (Sanford's brown lemur), Galago moholi(South African galago), Hapalemur griseus (Bamboo lemur), Lemur catta (Ring-tailed lemur), Loris tardigradus (Slender loris), Microcebus murinus (Gray mouse lemur), Mirza zaza (Northern giant mouse lemur), Nycticebus coucang (Slow loris), Otolemur crassicaudatus (Greater galago), Perodicticus potto (Potto), Propithecus diadema (Diademed sifaka), Propithecus tattersalli (Golden-crowned sifaka), Varecia rubra (Red ruffed lemur).

**Primates from Busch Gardens**

The blood samples from chimpanzees (Pan troglodytes, n=2), gorillas (Gorilla, n=3)

Orangutan (n=1, Pongo pygmaeus), red ruffed lemur (n=1, Varecia variegata), and White-fronted marmoset (n=1, Callithrix geoffroyi) were opportunistically collected and banked during routine health exams from these zoo-based animals located at Busch Gardens Tampa (Tampa, Florida).

**Human tissue samples**

We analyzed previously generated methylation data from n=1352 human tissue samples (adipose, blood, bone marrow, dermis, epidermis, heart, keratinocytes, fibroblasts, kidney, liver, lung, lymph node, muscle, pituitary, skin, spleen) from individuals whose ages ranged from 0 to 101 years. Out of the 1352 tissues, n=655 came from women.

The tissue samples came from four sources: tissue and organ samples from the National NeuroAIDS Tissue Consortium [12,13], Blood samples from the Cape Town Adolescent Antiretroviral Cohort study [14] and the PEG study [15], skin and other primary cells provided by Ken Raj [16]. Ethics approval (IRB#18-000315). Human postmortem samples are from previously generated methylation data (adipose, blood, bone marrow, dermis, epidermis, heart, keratinocytes, fibroblasts, kidney, liver, lung, lymph node, muscle, pituitary, skin, spleen) from individuals whose ages ranged from 0 to 93. The tissue samples came from three sources. Tissue and organ samples from the National NeuroAIDS Tissue Consortium [12,13]. Blood samples from the Cape Town Adolescent Antiretroviral Cohort study [14]. Additional blood samples from the PEG study. Skin and other primary cells provided by Kenneth Raj [16]. Ethics approval (IRB#15-001454, IRB#16-000471, IRB#18-000315, IRB#16-002028).

**Vervet monkeys [6]**

All animals used in this study were Caribbean-origin vervet monkeys (*Chlorocebus sabaeus*) from the VRC at Wake Forest School of Medicine. The VRC colony is an extended multigenerational pedigree established from 57 founders imported from the islands of St. Kitts and Nevis in the West Indies. The introduction of new animals to the pedigree ended in the mid-1980s (Jasinska et al. 2013). The colony members are socially reared in extended family groups mimicking the natural social composition of vervet monkey troops in the wild. Group sizes range from 11 to 23 animals, with one or two intact adult males included in each group. Unfamiliar males are rotated into each group every 3–5 years. The pedigree structure is genetically confirmed (Huang et al. 2015). All colony-born vervets have known chronological age accurate to 1 day.

Beyond applications in aging studies, animals from VRC are used in a wide range of research in areas such as the efficacy and enhancement of vaccines for infectious diseases, e.g., influenza and dengue (Kim et al. 2015; Holbrook et al. 2016; Briggs et al. 2014), investigations of diabetes, metabolic disease and obesity (Kavanagh et al. 2017; Kavanagh et al. 2016; Kavanagh et al. 2013); and the development of novel non-invasive biomedical imaging methodologies (Prabhakaran et al. 2017; Maldjian et al. 2014).

**Ethics statement**

The Wake Forest School of Medicine facilities are certified by the Association for Assessment and Accreditation of Laboratory Animal Care. The animal handling and sample collection procedures in this study were performed by a veterinarian after review and approval by the UCLA and VA Institutional Animal Care and Use Committees. Both housing and sample collection were in compliance with the US National Research Council Committee's Guidelines for Care and Use of Laboratory Animals (National Research Council et al. 2011) and met or exceeded all standards of the Public Health Service's "Policy on the Humane Care and Use of Laboratory Animals" (Office of Laboratory Animal Welfare n.d.).

**Vervet tissue samples**

For this study, we selected a total of 240 samples representing the entire vervet lifespan, from neonatal to senile stages: 144 samples from the peripheral blood, 48 samples from the liver, and 48 samples from the cortical brain area BA10. The brains were perfused to remove blood prior to dissection (Jasinska et al. 2017). The targeted brain area BA10 was very small, and brain samples were dissected as bulk tissues, collecting, to the extent feasible without the benefit of microscopy, the full thickness of the cortex while avoiding the underlying white matter (Jasinska et al. 2017). One outlier blood sample (202943350003_R03C01 from animal 1992020) was excluded from analysis on the basis of the DNAm profile. The remaining 143 blood samples included 14 pairs of biological replicates collected from 14 individuals at two different time points 3.9–10.93 years apart. Peripheral blood was collected through venipuncture with standard procedures. Liver and brain cortical tissues were collected during necropsies (Jasinska et al. 2017).

Genomic DNA was isolated from blood and liver samples primarily through Puregene chemistry (Qiagen). DNA from the liver was extracted manually and that from the blood was extracted with an automated Autopure LS system (Qiagen). DNA was extracted from old liver tissues and clotted blood samples manually with a QIAamp DNA Blood Midi Kit and DNeasy Tissue Kit according to the manufacturer's protocol (Qiagen, Valencia, CA). DNA from BA10 was extracted on an automated nucleic acid extraction platform AnaPrep (Biochain) with a magnetic bead based extraction method and Tissue DNA Extraction Kit (AnaPrep).

**Common marmosets [7]**

**Animal care and maintenance**

All marmosets used in this research were housed at the Barshop Institute for Longevity and Aging Studies at UT Health San Antonio (UTHSA). The Institutional Animal Care and Use Committee (IACUC) of UTHSA is responsible for monitoring housing and animal condition regularly to ensure all guidelines are met for the safety and health of the animals. This research was reviewed and approved by the UTHSA IACUC and experiments were conducted in compliance with the US Public Health Service's Policy on Humane Care and Use of Laboratory Animals and the Guide for the Care and Use of Laboratory Animals and adhered to the American Society of Primatologists (ASP) principles for the ethical treatment of non-human primates. Animals used in this study were based on age as well as a record of relatively good health as assessed by veterinary examination. Animals in this study were born either at UTHSCSA or transferred from the Southwest National Primate Research Center (SNPRC) in San Antonio TX. At UTHSA, animals were maintained using a modified specific pathogen-free barrier facility [17].

Each animal received three diet choices daily provided ad libitum: Harlan Teklad purified marmoset diet (TD99468), Mazuri Callitrichid gel diet (5MI5) and ZuPreem.

Blood draws and clinical blood counts/chemistry: All blood draws were taken during the morning hours of 08:00-11:00 from fed, non-anesthetized animals restrained in a custom assembly. Femoral vein blood collection (1.0-2.0 mL) was performed on each animal and blood was placed in to PAXgene blood collection tubes (Qiagen). Whole blood was frozen and stored at -80˚ C for shipment to UCLA for methylation analysis.

**Study samples**

For these studies we selected samples representing the entire primate lifespan, from neonate to old age. Genomic DNA was isolated from tissue samples mostly using Puregene chemistry (Qiagen). DNA from liver was extracted manually and from blood using an automated Autopure LS system (Qiagen). From old liver tissues and clotted blood samples DNA was extracted manually using QIAamp DNA Blood Midi Kit and the DNeasy Tissue Kit according to manufacturer's protocol (Qiagen, Valencia, CA). DNA from BA10 was extracted on an automated nucleic acid extraction platform Anaprep (Biochain) using a magnetic bead-based extraction method and Tissue DNA Extraction Kit (AnaPrep).

**Rhesus macaque [5]**

In total, we analyzed N=281 rhesus macaque tissue samples from 8 different sources of DNA. The rhesus monkeys have been housed continuously at the NIH Animal Center, Poolesville, MD. The animal center is fully accredited by the American Association for Accreditation of Laboratory Animal Care, and all procedures were approved by the Animal Care and Use Committee of the NIA Intramural Program. Monkeys were of a heterogenous genetic background, both Chinese and Indian origin.

Monkeys were housed individually in standard nonhuman primate caging on a 12h light/12h dark cycle, room temperature 78+/-2 degrees humidity at 60+/-20%. All monkeys had extensive visual, auditory, and olfactory but limited tactile contact with monkeys housed in the same room. Monkeys received 2 meals per day at estimated ad libitum levels throughout the study. Water was always available ad libitum. Monkeys were monitored minimally 3 times daily by trained animal care staff.

**Sample Collection**

Monkeys were fasted overnight, approximately 16-18 hours. Monkeys were anesthetized with either Ketamine, 7-10 mg/kg, IM or Telazol, 3-5 mg/kg, IM. Blood samples were obtained by venipuncture of the femoral vein using a vacutainer and EDTA tubes. Samples were immediately placed on dry ice and stored at -80 degrees. Skin samples were collected at the same time from an alcohol-wiped area of the back between the shoulder blades. Omental fat, kidney, liver, lung, skeletal muscle, and brain cortex were collected during necropsies scheduled for other study purposes. At that time, tissues were flash frozen in liquid nitrogen following collection and stored at -80 degrees. These tissues were selected for use based on having matching blood samples. None of the monkeys were sacrificed for this study.

**Prairie voles [18]**

**<u>Prairie vole colony</u>**

Male and female prairie voles (*Microtus ochrogaster*) were produced from laboratory-bred colonies at Cornell University, from breeding pairs that were offspring of wild caught animals captured in Champagne County, Illinois, USA. Voles are weaned and housed with littermates on postnatal day (PND) 21, and then housed with same-sex littermates after PND42-45. All animals received rodent chow (Laboratory Rodent Diet 5001, LabDiet, St. Louis, MO, USA) and water *ad libitum* and were maintained under standard laboratory conditions (14L:10D cycle, lights on at 08:00, 20 ± 2 °C) in transparent polycarbonate cages (29 x 18 x 13 cm) lined with Sani-chip bedding and provided nesting material. All experimental procedures were conducted and approved by the Institutional Animal Care and Use Committee (IACUC) of Cornell University (2013-0102) and were in accordance with the guidelines set forth by the National Institutes of Health.

**<u>Prairie vole tissue sample collection</u>**

Ear, liver, and brain samples from the Cornell University prairie vole colony were collected from 48 male and female prairie voles at various life stages: neonatal (<1 month old), sub-adult (2-4 months old), mature adult (4-10 months old), and middle aged/old adult (>10 months old). The pair bonded male and female prairie voles used in our study cohabitated with their partners for several months and produced at least three generations of litters. Animals were euthanized via rapid decapitation, their tissues rapidly extracted and frozen on dry ice before being stored at -80C until further processing for genomic DNA extraction. Brains were coronally sectioned and brain regions from the pair bonding circuit (PBC) were micro-dissected and pooled for each animal. The PBC brain regions included the prefrontal cortex, nucleus accumbens, lateral septum, ventral pallidum, and medial amygdala, and ventral tegmental area [19]. Genomic DNA was isolated and purified using the phenol-chloroform extraction and ethanol precipitation method. A total of 144 tissue samples were collected and processed for DNA methylation analysis. One animal was removed from the study due to a mismatch with the reported sex and our DNA methylation-based sex estimator.

**Peromyscus [20]**

Deer mice are maintained as outbred, genetically diverse closed colonies in the Peromyscus Genetic Stock center of the University of South Carolina. The study was approved by the Institutional Animal Care and Use Committee (IACUC) of the UofSC (protocol #: 2356-101506-042720) and were in accordance with the guidelines set forth by the National Institutes of Health. DNA was isolated from live animals by tail snips, or upon sacrifice from livers and brains by using DNeasy DNA isolation kit (Qiagen).

**Horses [21]**

**Ethics**

This collection protocol was approved by the UC Davis Institutional Animal Care and Use Committee (Protocol#19037). All collection protocols were approved by the UC Davis Institutional Animal Care and Use Committee (Protocols #20751 and 21455, respectively).

We generated DNA methylation data from n=42 different horse tissues collected at necropsy. The tissue atlas was generated from two Thoroughbred mares as part of the FAANG initiative [22], with the following tissues profiled: adipose (gluteal), adrenal cortex, blood (PBMCs; only n=1 mare), cartilage (only n=1 mare), cecum, cerebellum (2 samples each from lateral hemisphere and vermis), frontal cortex, duodenum, fibroblast, heart (2 samples each from the right atrium, left atrium, right ventricle, left ventricle), hypothalamus, ileum, jejunum, keratinocyte, kidney (kidney cortex and medulla), lamina, larynx (i.e. cricoarytenoideus dorsalis muscle), liver, lung, mammary gland, mitral valve of the heart, skeletal muscle (gluteal muscle and longissimus muscle), occipital cortex, ovary, parietal cortex, pituitary, sacrocaudalis dorsalis muscle, skin, spinal cord (C1 and T8), spleen, suspensory ligament, temporal cortex, tendon (deep digital flexor tendon and superficial digital flexor tendon), uterus [22]. These tissues were also used for RNAseq analyses.

Blood samples were collected via venipuncture into EDTA tubes from across 24 different horse breeds (buffy coat). Most of the samples were from the Thoroughbred (TB) (n=79) and American Quarter Horse breeds (QH, n=62). For the following breeds, we had between one and six blood samples: Andalusian, Appaloosa, Arabian, Dutch Warmblood, Hanoverian, Holsteiner, Irish Sport Horse, Lipizzaner, Lusitano, mixed breed, Oldenburg, Paint or Paint cross, Percheron, Shire, Standardbred, Warmblood and Welsh Pony. The n=49 liver samples originated from necropsy collections of horses across 19 different breeds, with most of the liver samples from QHs (n=20).

**Naked mole-rat [23]**

The NMR tissue samples were provided by two different labs: 1) Vera Gorbunova and Andrei Seluanov from the University of Rochester and 2) Chris Faulkes from the Queen Mary, University of London.

**Animals from the University of Rochester**

All animal experiments were approved and performed in accordance with guidelines set up by the University of Rochester Committee on Animal Resources with protocol number 2009-054 (naked mole-rat). Naked mole-rats were from the University of Rochester colonies. All animals in the colonies are microchipped and their ages are recorded. Housing conditions are described previously [24]. All tissues except for skin biopsies and blood were obtained from frozen tissue collection at the University of Rochester from healthy animals that were euthanized for other studies. Skin biopsies (2 mm punch) were collected from the backs of the animals under local anesthesia. Blood samples were collected from the tails. The n=3 induced pluripotent stem cells from NMR were generated as described in [25]. As control set for the iPS study, we used n=3 fibroblasts samples from animals aged 1 and 2.

Genomic DNA was extracted using Qiagen DNeasy Blood and Tissue kit and quantified using Nanodrop and Qubit.als

**Study Animals from Queen Mary University**

Naked mole-rats were maintained in the Biological Services Unit at Queen Mary University of London in accordance with UK Government Animal Testing and Research Guidance. The tissues used in this study were obtained from post-mortem specimens from animals free from disease in compliance with national (Home Office) and institutional procedures and guidelines. Because sample collection was from post-mortem material, additional local ethical approval was not required for this study. Tissue samples were snap frozen in liquid nitrogen following dissection and transferred for storage at -80ºC.

**Sheep [26]**

Sheep DNA samples for this study were derived from two distinct tissues from two strains: ear tissue from New Zealand Merino, and blood from South Australian Merino.

**<u>Sheep ear</u>**

Ear tissue was obtained from females and both intact and castrated male Merino sheep during routine on-farm ear tagging procedures in Central Otago, New Zealand. As a small piece of tissue is removed during the ear tagging process that is usually discarded by the farmer, we were able to source tissue and record the year of birth without altering animal experience, in accordance with the New Zealand Animal Welfare Act (1999) and the National Animal Ethics Advisory Committee (NAEAC) Occasional Paper No 2 [26]. The exact date of birth for each sheep is unknown, however, this was estimated to be the 18th of October each year, according to the date at which rams were put out with ewes (May 10th of each year), a predicted mean latency until mating of 1Tab2 days, and the mean gestation period from a range of sheep breeds (149 days) [27]. Castration was performed by the farmer using the rubber ring method within approximately 5-50 days from birth as per conventional farming practice [28]. Mass of yearlings was recorded by the farmer for both castrated and intact male sheep at 6.5 months of age, as a part of routine growth assessment. In total, ear tissue from 138 female sheep aged 1 month to 9.1 years and 126 male sheep (63 intact, 63 castrates) aged 6 months to 5.8 years was collected and subjected to DNA extraction.

DNA was extracted from ear punch tissue using a Bio-On-Magnetic-Beads (BOMB) protocol [29] which isolates DNA molecules using solid-phase reversible immobilisation (SPRI) beads. Approximately 3 mm punches of ear tissue were lysed in 200 µL TNES buffer (100 mM Tris, 25 mM NaCl, 10 mM EDTA, 10% w/v SDS), supplemented with 5 µL 20 mg/mL Proteinase K and 2 µL RNAse A and incubated overnight at 55 °C as per BOMB protocols. The remainder of the protocol was appropriately scaled to maximise DNA output while maintaining the necessary 2:3:4 ratio of beads:lysate:isopropanol. As such, 40 µL cell lysate, 80 µL 1.5X GITC (guanidinium thiocyanate), 40 µL TE-diluted Sera-Mag Magnetic SpeedBeads (GE Healthcare, GEHE45152105050250) and 80 µL isopropanol were combined. After allowing DNA to bind the SPRI beads, tubes were placed on a neodymium magnetic rack for ~5 minutes until the solution clarified and supernatant was removed. Beads were washed 1x with isopropanol and 2x with 70% ethanol, and then left to air dry on the magnetic rack. 25 µL of MilliQ $H_2O$ was added to resuspend beads, and tubes were removed from the rack to allow DNA elution. Tubes were once again set onto the magnets, and the clarified solution (containing DNA) was collected.

DNA was quantified using the Quant-iT PicoGreen dsDNA assay kit (ThermoFisher Scientific, cat # P11496). 1 µL DNA sample was added to 14 µL TE diluted PicoGreen in MicroAmp optical 96-well plates (ThermoFisher Scientific, cat #N8010560) as per manufacturer directions, sealed, and placed into a QuantStudio qPCR machine for analysis. Samples with DNA content greater than the target quantity of 25 ng/µL were diluted with MilliQ.

**Sheep blood**

DNA methylation was analysed in DNA extracted from the blood of 153 South Australian Merino sheep samples (80 transgenic Huntington's disease model sheep (OVT73 line) [30] and 73 age-matched controls) aged from 2.9 to 7.0 years. All protocols involving OVT73 sheep were approved by the Primary Industries and Regions South Australia (PIRSA, Approval number 19/02) Animal Ethics Committee with oversight from the University of Auckland Animal Ethics Committee. The epigenetic age of the transgenic sheep carrying the *HTT* gene was not significantly different from controls (p=0.30, Mann-Whitney U test), therefore the data derived from these animals was subsequently treated as one dataset.

300 µL thawed blood samples were treated with 2 rounds of red cell lysis buffer (300 mM Sucrose, 5 mM $MgCl_2$, 10 mM Tris pH8, 1% Triton X-100) for 10 minutes on ice, 10 minute centrifugation at 1,800 RCF, and supernatant removed between each buffer treatment. The resulting cell pellet was incubated in cell digestion

buffer (2.4 mM EDTA, 75 mM NaCl, 0.5 % SDS) and Proteinase K (500 µg/ml) at 50 °C for two hours. Phenol:Chloroform:Isoamyl alcohol (PCI, 25:24:1; pH8) was added at equal volumes, mixed by inversion, and placed in the centrifuge for 5 minutes at 14,000 RPM at room temperature (repeated if necessary). The supernatant was collected and combined with 100% ethanol at 2x volume, allowing precipitation of DNA. Ethanol was removed and evaporated, and 50 µL TE buffer (pH8) was added to resuspend genomic DNA. DNA sample concentration was initially quantified using a nanodrop, followed by Qubit.

**Pig [27]**

**<u>Porcine samples</u>**

All animal procedures were approved by the University of Illinois and University of Wisconsin Institutional Animal Care and Use Committee, and all animals received humane care according to the criteria outlined in the Guide for the Care and Use of Laboratory Animals. Porcine whole blood samples (n=146) were collected from female Large White X Landrace crossbred domestic pigs (n=84, age range 11 – 2,285 days) and Wisconsin Miniature Swine™ (n=60, age range 8 – 1,880 days) at the University of Wisconsin-Madison. Whole blood (n=16) and tissue samples (bladder, frontal cortex, kidney, liver, lung; n=16/tissue type) were collected from 16 Large White X Minnesota minipig crossbred pigs (n=9 female, n=8 male, age range 29 – 1,447 days) at the University of Illinois at Urbana-Champaign. All blood samples were collected in EDTA tubes, aliquoted, and flash frozen in liquid nitrogen within 10 minutes of collection. Tissue samples were collected and flash frozen within 10 minutes of euthanasia. All samples were stored at -80 until processing. Samples were shipped to the University of California, Los Angeles Technology Center for Genomics & Bioinformatics for DNA extraction and generation of DNA methylation data.

**Odontocete species [28-30]**

**<u>Ethics approval</u>**

The study was authorized by the management of each institution and was reviewed by their respective zoo research and animal use committees.

**Study Animals**

For model development, our study population included 293 animals from nine species of odontocetes, of which the majority were from four species including beluga (n = 66), Pacific white-sided dolphins (n = 17), killer whales (n = 37) and bottlenose dolphins (n = 137), housed at nine Association of Zoos and Aquarium (AZA), Alliance for Marine Mammal Parks and Aquariums or Japanese AZA accredited zoological institutions. Known (77.8%) or estimated (based on length at capture or rescue for stranded animals) birth dates were provided by each housing institution. In addition to zoo-based animals, we included 19 skin samples from free-ranging Norwegian killer whales to the training set because the ages of these animals could be estimated within sufficient accuracy (expected error less than 8%) on the basis of several lines of evidence including GLG counts (n = 3), whereby 1 GLG was assigned per year of age [12], length at necropsy (n = 1), juvenile at first identification (n = 2) or minimum age estimation based on length and maturity at first sighting (n = 13). The remaining 26 animals' ages could not be estimated within our accuracy parameters and skin samples from these animals were used to demonstrate model application for determining age and sex in wild animals.

**Sample collection**

Blood samples (0.5 ml min) were collected either voluntarily from the peripheral periarterial venous rete on the ventral tail fluke using an 18 to 22 gauge winged blood collection set or attached to a vacutainer collection system. Blood was collected by either the veterinary technician or veterinarian on staff and into BD Vacutainers (Becton Dickinson, Franklin Lakes, NJ) containing EDTA. Samples were inverted in the Vacutainer a minimum of 10 times and then frozen at -80°C until further testing.

Skin samples (~0.5 gm) were collected either under stimulus control or manual restraint using a sterile disposable dermal curette (Miltex, Integra Life Sciences Corp.,York, PA) from a location just posterolateral of the dorsal fin overlying the epaxial muscle. Prior to collection, a cold pack was placed on the site for several minutes prior to sampling to numb the sample site. Skin samples were placed into sterile cryovials (Nunc® Cryotubes, MilliporeSigma Corp., St. Louis, MO) and stored at -80$^{o}$C until shipment on dry ice. Skin samples from non-living animals were obtained from frozen (-80°C) specimens that had been previously collected and stored during standard necropsy procedures.

Skin samples were sectioned from previously collected killer whale biopsy samples of 45 unique individuals (photo-identified) collected in August and November 2017 and from April through July 2018 in northern Norway (Jourdain, 2017). The killer whales were biopsied using an ARTS darting system (Restech, Bodø, Norway) and 25 × 9 mm or 40 × 9 mm stainless steel tips in 2017, and with an injection gun (Pneu-Dart Inc., Williamsport, PA) and 25 × 7 mm tips in 2018 as previously described (Jourdain, 2017). In addition, tissue samples were collected from six dead, stranded killer whales, and one other individual by-caught in a herring purse-seine, in northern Norway between 2015 and 2017 (E. Jourdain, unpublished data). Skin samples were collected from the region directly posterior to the dorsal fin and stored at -20°C until analysis.

**Beluga whales [30]**

Skin tissue samples were collected from carcasses of beluga whales that were beach-cast, stranded dead, or taken during subsistence hunting between 1992 and 2015 in Cook Inlet, Alaska, USA (NMFS Research Permit 932-1905-00/MA-009526 through the Marine Mammal Health and Stranding Response Program). Skin samples were preserved in a salt and dimethyl sulfoxide (DMSO) solution and archived at NOAA's Southwest Fisheries Science Center in La Jolla, California, USA. A total of 69 individuals were selected for the clock calibration dataset, and their chronological ages were estimated by counting tooth growth layer groups. The final calibration dataset included 67 individuals due to inconsistent molecular sex data. Teeth were analyzed by at least two readers using methods validated in Lockyer et al. (2007), and a consensus age provided by NOAA was used in this study. When individuals were represented by multiple teeth in the dataset, the oldest age estimate was used to mitigate error from tooth wear (*e.g.,* the count from the tooth with the greatest number of growth layer groups).

Samples of skin tissue from living CI beluga whales were collected with a biopsy dart in 2016, 2017, and 2018 (NMFS ESA/MMPA Permit #20465; McGuire et al., 2017). Biopsy samples were frozen in the field in liquid nitrogen and later subsampled at the NOAA Alaska Fisheries Science Center in Seattle, Washington, USA. Genomic DNA was extracted from tissue samples using a standard phenol-chloroform protocol. Extracted DNA was treated with RNAse A (1 µL of 1 mg/mL added to samples of 100 µL for 30 minutes at room temperature) and then purified and concentrated using a DNA Clean and Concentrator-5 Kit (Zymo Research

Corp., USA). The concentration of genomic DNA was measured on a QUBIT 4 fluorometer (ThermoFisher Scientific, USA).

**Killer whales and bowhead whales**

**Animal Use and Ethics**

For bowhead subsistence hunts, indigenous hunters had the authorization to conduct hunts and collected samples on behalf of Fisheries and Oceans Canada. Bowhead whale biopsy samples were collected in 2019 under Fisheries and Oceans Canada (DFO) license to Fish for Scientific Purposes (LFSP) S-19/20-1007-NU and Animal Care approval (AUP) FWI-ACC-2019-14. Skin samples from eastern North Pacific killer whales were collected as previously described (Ford et al. 2018b) under NMFS General Authorization No. 781–1725, and scientific research permits 781-1824-01, 16163, 532- 1822-00, 532– 1822, 10045, 18786-03, 545-1488, 545-1761, and 15616.

**Killer whale biopsy samples & DNA extraction**

Killer whale populations in the eastern North Pacific are among the most intensively studied cetacean populations globally. The so-called 'resident' killer whale populations inhabiting the coastal waters from California to Alaska comprise individually identified whales that have been studied for over 40 years (Bigg 1982, Balcomb and Bigg 1986, Matkin et al. 1999, Ford et al. 2000, Matkin et al. 2014). These longitudinal studies and reliable identification of individual whales within populations through annual photographic census data has provided unique insight into population dynamics and demographics. The resolution provided by annual documentation of births, ages at physical maturity (males) and age at first parturition (females) provides an unparalleled opportunity to validate epigenetic models of age from whale skin.

The killer whale validation dataset is rare in both the representation across age classes and the number of known-age samples from a wild population based on direct observations (Bigg et al. 1990, Olesiuk et al. 1990, Matkin et al. 1999, Olesiuk et al. 2005, Matkin et al. 2014), providing an ideal training set for validating an epigenetic clock. Killer whale longevity is estimated to be 80 or 90 years (Olesiuk et al. 1990, Olesiuk et al. 2005) and samples in the current dataset represent individual whales ranging from age 0 (neonate) to 79 yrs. Reflecting killer whale age-related mortality patterns (Olesiuk et al. 2005), the number of individuals representing older age classes diminishes as expected with n = 11 whales estimated be > 50 yrs old based on size, physical and reproductive maturity at the time of first observation.

Ages of individual killer whales from the Southern Resident killer whale and Alaska Resident killer whale populations were determined based on the sex and size of the animal during the year that it was first documented, following Olesiuk et al. (1990). Whales born during the study (post-1974) were aged in reference to the year in which they were born. Ages for whales that were juveniles or adults when field observations began in the early 1970s were aged based on the year they reached physical maturity or, for females, the year they gave birth to their first viable offspring (Olesiuk et al. 1990). Confidence estimates (0% - 100%) were assigned to each individual sampled whale included in the dataset reflecting the certainty around age estimates, frequency of encounters with the individual and age or state of physical maturity at the time of first identification (Olesiuk et al. 1990, Olesiuk et al. 2005, Matkin et al. 2014). Genetic samples from 142 killer whales (supplementary info) were included in the dataset representing 118 different killer whales of age 0 yrs (neonate or fetus; 100% certainty) to age 75+ yrs (50% certainty).

Epidermal samples were collected from live killer whales using remote dart biopsy methods, and from dead stranded animals during routine post-mortem necropsy protocols (Barrett-Lennard et al. 1996, Parsons et al. 2003). Identities of individual whales were recorded photographically whenever possible. Total genomic DNA was extracted from skin biopsies either using a silica-membrane kit following manufacturer’s protocols (DNeasy Blood and Tissue kit, Qiagen, Valencia, CA), or following a standard proteinase K phenol/chloroform/isoamyl alcohol extraction protocol (Sambrook et al. 1989).

**Bowhead whale**

**<u>Biopsy samples & age estimates</u>**

Bowhead whales are very slow-growing and extremely long-lived baleen species that undergo periods with rapid growth (as a fetus), pauses in growth (ages 1 to about 6-7) and slow growth (age 8+) which gradually slows even more as they age (George and Thewissen 2020). For this reason, accurate age estimates of individual bowhead whales are typically limited to early life stages when multiple sources of information are available.

Estimating age for individual bowhead whales requires a combination of several different types of data (summarized in George and Thewissen (2020)) including body length, length of the longest baleen plate, body condition measurements, other morphological measurements that include frequency of scars, colour of the peduncle and chin region, and aspartic acid racemization (AAR) analyses from eye lenses. Despite multiple data sources, bowhead age can only be approximated for whales more than approximately 7 years old because the variation in size at age is many times larger than the annual growth rate of individual whales. Whales 2-7 years old cannot be accurately aged without information on the longest baleen plate, which is only available for harvested whales, although body condition provides information to assist with age estimation. Bowhead whale skin samples were collected from remote biopsy using a crossbow during photographic studies conducted in Cumberland Sound in (Young et al. 2019), and samples collected during subsistence harvest. Data available for age estimates were limited to body length, body condition, frequency of scars, and colour of the peduncle region for biopsy samples. Skin samples collected during subsistence harvest included age estimates based on body length, length of the longest baleen plate, notes on scars or peduncle colour and AAR ages. Where multiple sources of data were available, ages obtained from AAR or age-at-length estimates were adjusted to take account of the additional data as below.

Age at length was estimated using data in Figure 7 of Lubetkin et al. (2012) with estimation for gradually reduced growth rates for whales older than 60 years old as suggested by their data and those of Koski et al. (1992). Yearlings were confirmed among the smallest whales by body condition using criteria in Koski et al. (2010). Body condition was also used to adjust the age of whales 2-7 years as determined by length because the youngest of these whales (i.e., 2 years old) appeared to have the poorest body condition and body condition appeared to improve as young whales age (see Figure 3 in Koski et al., 2010).

Age based on aspartic acid racemization was determined for 11 harvested bowhead whales (4 females; 7 males). Both eyeballs were stored at –20°C immediately after dissection from harvested whales, and one eyeball per whale was subsequently used for age estimation. Dissection of eye lenses and age estimation by the aspartic acid racemization (AAR) technique was performed using methods described in (Garde et al.

2007). Estimates of individual D/L ratios were converted to age estimates as described in (George et al. 1999b, Rosa et al. 2004, Heide-Jørgensen et al. 2012, Rosa et al. 2013).

**Humpback whales**

**<u>Ethics</u>**

Skin samples were collected by the Center for Coastal Studies under research permits issued by the U.S., National Marine Fisheries Service (21485, 16325, 20465, 14245, 633-1483, 633-1778, 932-1905), the Canadian Department of Fisheries and Oceans and IACUC #NWAK-18-02.

**<u>Humpback sample collection</u>**

Skin samples were collected from live North Atlantic humpback whales by biopsy sampling techniques (Palsbøll, 1991). Sampling was performed in the Gulf of Maine between 2003 and 2020 under the authorization of nationally-issued research permits. Samples were refrigerated or frozen in the field and then archived at -80C without chemical preservative.

Year of birth can be determined precisely for humpback whales first encountered as calves during an obligatory period of maternal dependency (Clapham, 1992;Baraff). Otherwise, there are no reliable outward indicators of chronological age in this species. We therefore selected known-age samples for this study based on data from a long-term study of individual whales. The identity of each sampled individual was confirmed through photo-identification techniques (Katona, 1981) using a reference catalog of the Gulf of Maine population curated by the Center for Coastal Studies (Provincetown, MA). For individuals first seen as calves, age at sampling was the number of years between sampling and birth. However, the exact date of birth in that year was not known for any individual, and sampling was performed over a wide window (April through November) outside of the winter breeding period. We therefore used an estimated day of birth at the peak of the winter breeding season for all individuals (February 15) to refine age at the time of sampling. Photo-identification research on this population did not begin until the 1970s and individuals cataloged since time were not all seen first as calves. Thus, chronological age is unknown for many individuals, including the oldest whales in the population. We therefore selected samples from individuals with long sighting spans (as long or longer than the upper 20% of the available known-age data) to clarify epigenetic age patterns at and beyond the top of the validated age range. In these cases, we calculated a minimum age at sampling based on the fact the individual could not have been born later than the year before their first sighting. However, such whales could have been born in any earlier year and so this provided only a minimum bound on their chronological age.

The sex of sampled individuals was known independently from molecular genetic analysis (Bérubé, 1996), in some cases supplemented by observations of the genital slit (Glockner, 1983) or calving history.

**Cats [31]**

Ethics. Sample collection was approved by the Clinical Research Ethical Review Board of the RVC (URN: 2019 1947-2).

**Feline and other animal blood samples**

The DNA archive of the Royal Veterinary College (RVC) was searched for feline ethylenediaminetetraacetic acid (EDTA) blood samples that were residuals from previous routine hematology testing. Cats were selected to represent the widest age range possible based on the available samples with a uniform distribution across the entire range, available breeds and neutering status. As the samples originated from cats that were presented for veterinary investigation, cats were selected to have no or minimal abnormalities on available laboratory data (hematology, serum biochemistry, endocrinology), reviewed by a board certified veterinary clinical pathologists (BSz). The DNA samples were maintained frozen at -80 °C for various amount of time (0-11 years). Samples from guinea pigs, rabbits, ferrets, and alpacas were also residual samples from routine patients presented for veterinary care. Sample collection was approved by the Clinical Research Ethical Review Board of the RVC (URN: 2019 1947-2). Genomic DNA from cat blood was extracted using the Zymo DNA extraction kit according to the manufacturer’s instructions. DNA was eluted in water and quantified with picogreen kit according to the instructions provided.

**Non-domestic cat species**

Blood samples from cheetah (Latin name Acinonyx jubatus), lion (Panthera leo nubica), and tiger (Panthera tigris) were opportunistically collected and banked during routine health exams from these zoo-based animals located at Busch Gardens and White Oak Conservation.

**Elephants [32]**

Ethics. This study was authorized by the management of each participating zoo and, where applicable, was reviewed and approved by zoo research committees. In addition, the study received IACUC approval (#18-29) at the NZP; and endorsement from the elephant Taxon Advisory Group and Species Survival Plan.

**Study Animals**

Our study population included 140 elephants (57 African and 83 Asian) housed in 27 AZA-accredited zoos in North America (including Canada). Known or estimated birthdates were gleaned from each species' studbooks.

Uncertain surrounding age information is encoded in our variable "ConfidenceInAgeEstimate". A value of 90% indicates that the chronological age could be off by around 5 percent. Our analysis omitted animals for whom the confidence in the age estimate was less than 90 percent. For all elephants, the birthdates used in our analyses were taken from those recorded in the regional studbooks maintained through the Association of Zoos and Aquariums Taxon Advisory Group and Species Survival Plan for each species. Elephants had known birthdates if there were captive-born (i.e., they were recorded as having been born in a captive facility in the U.S. or in a range country), or were recorded as having an estimated birthdate if they were imported (i.e., capture location and date of capture was recorded in the studbook). How age was estimated for imported individuals is not described in the studbooks but was most likely done by the broker at the time of importation based on individual morphometric measurements. The majority (>70%) of the North American population is imported as captive breeding as a population management tool was not fully implemented until the late 1990s. In our sample set we had about equal proportion of captive vs imported individuals, although we did have more imported animals reflecting the management history of the population. Our samples are made up of 65 captive-born/known birthdates (21 Africans and 26 Asians), and 77 imported/estimated birthdates (36 African ad 27 Asian).

Whole blood samples from either an ear or leg vein directly into an EDTA tube were collected between 1998 and 2019 during regular veterinary examinations and shipped frozen to the genetics lab at the Smithsonian Conservation Biology Institute Center for Conservation Genomics (SCBI-CCG). The samples were stored in an ultralow freezer (-80ºC) until DNA extraction.

**Yellow-bellied marmots [33]**

Ethics. Data and samples were collected under the UCLA Institutional Animal Care and Use protocol (2001-191-01, renewed annually) and with permission from the Colorado Parks and Wildlife (TR917, renewed annually).

All samples were collected as part of a long-term study of a free-living population of yellow-bellied marmots in the Gunnison National Forest, Colorado (USA), where marmots were captured and blood samples collected biweekly during the their active season (May to August).

Individuals were monitored throughout their lives, and chronological age was calculated based on the date at which juveniles first emerged from their natal burrows. We only used female samples because precise age for most adult males is unavailable since males are typically immigrants born elsewhere. We selected 160 whole blood samples from 78 females with varying ages. From these, DNA methylation (DNAm) profiling worked well for 149 samples from 73 females with ages varying from 0.01 to 12.04 years.

**Roe deer [34]**

**<u>Study populations</u>**

We sampled roe deer living in two enclosed forests with markedly different environmental contexts: Trois-Fontaines (TF) and Chizé (CH). The Trois-Fontaines forest (1,360 ha) is located in north-eastern France (48°43’N, 4°55’E) and is characterized by a continental climate, moderately severe winters and warm and rainy summers. This site has rich soils and provides high quality habitat for roe deer (Pettorelli et al. 2006). In contrast, the Chizé forest (2,614 ha) is located in western France (46°50’N, 0°25’W) and is characterized by temperate oceanic climate with Mediterranean influences. This site has a low productivity due to poor quality soils and frequent summer droughts (Pettorelli et al. 2006), and thereby provides a quite poor habitat for roe deer in most years. Individuals from these two populations have been intensively monitored using a long-term Capture-Mark-Recapture program since 1975 and 1977 (for Trois-Fontaines and Chizé, respectively). In each site, 10-12 days of capture using drive-netting are organized every year between December and March (see Gaillard et al., 1993 for details on capture sessions), which allows capturing and measuring about half the population every year. Once a roe deer is captured, its sex and body mass (to the nearest 50g) are recorded and a basic clinical examination is performed. All individuals included in our analyses were of known age because they were either caught as newborn in spring (see Delorme et al. 1988 for further details) or as c.a. 8 months old during winter captures, when they still have their milk teeth (most often incisors and always premolars, Flerov 1952).

**Roe deer blood samples and DNA extraction**

In 2016 and 2017, we collected blood samples (up to 1mL per kg of body mass) from the jugular vein. Within 30 min of sampling, the blood was centrifuged at 3000 g for 10 min and the plasma layer was removed before washing the cells with an equivalent volume of 0.9% w/v NaCl solution. After a second centrifugation, the intermediate buffy coat layer, comprising mainly leukocytes, was collected in a 1.5-mL Eppendorf tube and immediately frozen at −80 °C in a porcos freezer (Telstar SF 8025) until further use.

We extracted genomic DNA from leucocytes using the Macherey-Nagel NucleoSpin® Blood QuickPure kit. DNA purity was assessed using a Nanodrop ND-1000 spectrophotometer (Thermo Scientific, Wilmington DE, USA). For all samples, the purity absorption range was 1.7 - 2.0 for the 260/280 nm ratio and >1.8 for the 260/230 nm ratio. We selected 96 samples by balancing the numbers of individuals among ages, and between populations and sexes. DNA concentration was determined spectrophotometrically using the Qubit assay kit. DNA samples were then diluted in ultrapure water to reach a concentration of ~70 ng.µl$^{-1}$ and displayed in a microplate to complete the DNA methylation protocol (see below). For 6 samples, the concentrations obtained after dilution were too low compared to the expected concentrations of 70 ng/µl and were excluded from the dataset. The 90 roe deer samples analysed in this study correspond to 79 individuals (i.e. 11 individuals were sampled both in 2016 and 2017) aged from 8 months to 13.5 years of age. This age range encompasses most of the roe deer lifespan as individuals older than 15 years of age are rarely observed in the wild (the oldest age ever recorded for a roe deer monitored in the wild being 17.5 years old, Gaillard et al. 1998).

**Zebras [35]**

**Ethics.**

Plains zebra samples were collected under a protocol approved by the Research Safety and Animal Welfare Administration, University of California Los Angeles: ARC # 2009-090-31, originally approved in 2009.

**Samples**

Both whole blood (96) and remote biopsy (24) samples were obtained from a captive population of zebras maintained in a semi-wild state by the Quagga Project [36] in the Western Cape of South Africa. The population was founded in 1989 with 19 wild individuals (9 from Etosha National Park in Namibia, 10 from the Kwazulu-Natal in South Africa). Since its inception, the population has undergone strong selection in an effort to reproduce the phenotype of the extinct quagga subspecies: no stripes on legs and hindquarters, and thinner and paler stripes in the head and barrel region. During sampling, individuals were uniquely identified by their stripe patterns and their ages were derived from studbook information about dates of birth, which were typically known within one month. One exception to this is a tissue sample from a founder that was sampled as a young mare and would have been at least 25 years old at sampling, but may have been slightly older. Remote biopsies were taken using an air-powered rifle affixed with a 1 mm wide by 20-25 mm deep biopsy dart and preserved in RNAlater (Qiagen). Blood samples were collected opportunistically during veterinarian visits, and preserved in EDTA tubes. Most samples were collected from different individuals, with the exception of two individuals that each were sampled twice some years apart. All samples were stored at -20 °C. After eliminating samples with low confidence for individual identity and age, we retained 76 blood samples and 20 biopsy samples, totaling 96 zebra samples.

Three zebra data sets were analyzed in our epigenetic models: (1) only blood samples, (2) only biopsy samples, and (3) blood and biopsy samples combined. The Grevy’s zebra (n=5) and Somali wild ass (n=7), are samples from zoo-based animals that were opportunistically collected and banked during routine health exams and the DNA methylation profiles from these samples have been reported previously [21].

**Rat [37]**

The rat tissues came from 4 different labs across three countries:(i) India: Nugenics Research in collaboration with School of Pharmacy SVKM's NMIMS University (K. Singh), (ii) United States: University of Tennessee Health Science Center (H. Chen) and Medical College of Wisconsin (L.C. Solberg Woods), and (iii) Argentina: University of La Plata (R. Goya).

**Rats from Tennessee and Wisconsin**

Blood samples (n=48): Male and female heterogeneous stock rats were bred at the Medical College of Wisconsin (Solberg Woods Lab) or University of Tennessee Health Science Center (Hao Chen Lab). Heterogeneous Stock (HS) populations were originally developed by breeding together eight inbred strains, followed by maintaining the colony in a manner that minimizes inbreeding, allowing fine-resolution genetic mapping of a variety of complex traits **51**. Rats were euthanized at different ages by an overdose of isoflurane (> 5%). Trunk blood was collected immediately and stored at −80 °C until processing. Blood samples were treated with streptokinase (60-80 IU/200 μl blood, overnight incubation at 37 °C) and DNA was extracted using the QiaAmp Blood Mini Kit (Qiagen Cat No./ID: 51304) following manufacturer's instructions. All procedures were approved by the Institutional Animal Care and Use Committee of the University of Tennessee Health Science Center or the Medical College of Wisconsin and followed the NIH Guide for the Care and Use of Laboratory Animals. Genomic DNA was isolated from tissue samples mostly using Puregene chemistry (Qiagen). DNA from liver was extracted manually and from blood using an automated Autopure LS system (Qiagen). From tissues and clotted blood samples DNA was extracted manually using QiaAmp DNA Blood Midi Kit and the DNeasy Tissue Kit according to manufacturer's protocol (Qiagen, Valencia, CA). DNA from BA10 was extracted on an automated nucleic acid extraction platform Anaprep (Biochain) using a magnetic bead-based extraction method and Tissue DNA Extraction Kit (AnaPrep).

**Rats from the University of La Plata (R. Goya lab)**

Multiple tissues/cell types (adipose, blood, cerebellum, hippocampus, hypothalamus, liver, neocortex, ovaries, pituitary, skin, substantia nigra): Young (3.7 mo., n=11), Late Adults (LA, 8.0 mo., n=9), Middle-Aged (M-A, 15.7 mo., n=6) and Old (25.5 mo., n=14) female Sprague-Dawley (SD) rats, raised in our Institute, were used. Animals were housed in a temperature-controlled room (22 ± 2ºC) on a 12:12 h light/dark cycle. Food and water were available *ad libitum*. All experiments with animals were performed in accordance to the Animal Welfare Guidelines of NIH (INIBIOLP's Animal Welfare Assurance No A5647-01) and approved by our Institutional IACUC (Protocol # P05-02-2017).

**Tissue sample collection**

Before sacrifice by decapitation, rats were weighed, blood was withdrawn from the tail veins with the animals under isoflurane anesthesia and collected in tubes containing 10µl EDTA 0.342 mol/l for 500µl blood. The brain was removed carefully severing the optic and trigeminal nerves and the pituitary stalk (not to tear the pituitary gland), weighed and placed on a cold plate. All brain regions were dissected by a single experimenter (see below). The skull was handed over to a second experimenter in charge of dissecting and weighing the adenohypophysis. The rest of the body was handed to other 2 or 3 experimenters who dissected and collected whole ovaries, a sample of liver tissue, adipose tissue and skin tissue from the distal portion of tails.

**Brain region dissection**

Prefrontal cortex, hippocampus, hypothalamus, substantia nigra and cerebellum were rapidly dissected on a cold platform to avoid tissue degradation. After dissection, each tissue sample was immediately placed in a 1.5ml tube and momentarily immersed in liquid nitrogen. The brain dissection protocol was as follows. First a frontal coronal cut was made to discard the olfactory bulb, then the cerebellum was detached from the brain and from the medulla oblongata using forceps. To isolate the medial basal hypothalamus (MBH), brains were placed ventral side up and a second coronal cut was made at the center of the median eminence (−3,6 mm referred to bregma). Part of the MBH was taken from the anterior block of the brain and the other part from the posterior block in both cases employing forceps. The hippocampus was dissected from cortex in both hemispheres using forceps. This procedure was also performed on the anterior and posterior blocks, alternatively placing the brain caudal side up and rostral side up. To dissect the substantia nigra, in each hemisphere a 1-mm thick section of tissue was removed from the posterior part of the brain (−4,6 mm referred to bregma) using forceps. Finally, the anterior block was placed dorsal side up, to separate prefrontal cortex. With a sharp scalpel, a cut was made 2 mm from the longitudinal fissure, and another cut was made 5 mm from it. Additionally, two perpendicular cuts were made, 3 mm and 6 mm from the most rostral point, obtaining a 9 $mm^2$ block of prefrontal cortex. This procedure was performed in both hemispheres and the two prefrontal regions collected in a code-labeled tube.

**Anterior pituitary**

Using forceps the dura matter that covers gland was removed leaving the organ free on the sella turcica. The neural lobe was carefully separated from the anterior pituitary (AP) which was then carefully lifted with fine curved tip forceps pointing upwards. It was rapidly weighed, then put in a tube and placed momentarily in liquid nitrogen.

**Ovaries**

The genital apparatus was dissected by cutting the mesentery to isolate the uterine horns, the tubular oviduct, the ovaries and the junction between the anus/rectum and the vulva/vagina, leaving the unit of the sexual organs and the urinary bladder isolated. The ovaries were carefully separated from the oviducts; the fat around the ovaries was also removed. Both gonads were placed in a single eppendorf tube and momentarily placed in liquid nitrogen.

**Liver**

Liver tissue extraction was made by cutting a piece of the median lobe (0.5 cm x 0.5 cm). Tissue was placed in a tube and momentarily stored immersed in liquid nitrogen.

**Adipose tissue**

Adipose tissue samples were obtained from the fatty tissue of the small intestine.

**Tail skin**

For skin tissue, 5 cm of a distal tail portion were cut with scissors. Skin was separated and hair removed using scalpel. Tissue was placed in a tube and stored as described for other tissues.

DNA was extracted from blood on an automated nucleic acid extraction platform called QiaSymphony (Qiagen) with a magnetic bead-based extraction kit, QIAsymphony DNA Midi Kit (Qiagen). DNA was extracted from tissue on an automated nucleic acid extraction platform called Anaprep (Biochain) with a magnetic bead-based extraction kit, Tissue DNA Extraction Kit (Biochain). DNA from brain regions was extracted using an automated nucleic acid extraction platform called QIAcube HT (Qiagen) with a column-based extraction kit, QIAamp 96 DNA QIAcube HT Kit (Qiagen).

**Rats from Nugenics Research Group**

Sprague Dawley rats of both sexes were used, from which blood, whole brain, heart and liver were harvested. Two batches or samples were prepared: the first batch was intended for training the epigenetic clock: n=42 blood samples, n=18 whole brain, n=18 heart, n=18 liver samples. The second batch of test set involved n=76 tissue samples (n=22 blood, n=18 liver, n=18 heart, n=18 hypothalamus). The test data were used to evaluate the effect of the treatment in 3 conditions: young (30-week old) and treated old samples (109 weeks old), and untreated old samples (again 109 weeks old). We evaluated 4 sources of DNA: blood (n=18), liver (n=18), heart (n=18) and hypothalamus (n=18). Ethics committee approval number - CPCSEA/IAEC/P-6/2018.

Male Sprague Dawley rats of 8 weeks (200–250 g) and 20 months (400-450g) were procured from the National Institute of Bioscience, Pune, India. Animals were housed in the animal house facility of School of Pharmacy, SVKM's NMIMS University, Mumbai during the study under standard conditions (12:12 h light: dark cycles, 55-70% of relative humidity) at 22±2°C temperature with free access to water and standard pellet feed (Nutrimix Std-1020, Nutrivet Life Sciences, India). The animals were acclimatized to laboratory environment for seven days before initiation of the study. The experimental protocol was approved by the Institutional Animal Ethics Committee. The approval number is CPCSEA/IAEC/P-75/2018. Rats were euthanized at different ages by an overdose of isoflurane (> 5%). Trunk blood was collected immediately and stored at −80 °C until processing. 100 µl blood sample was treated with 20 µl Proteinase K and then the volume was adjusted to 220 µl with Phosphate Buffer Saline (PBS) in 1.5 ml or 2 ml microcentrifuge tube. 200 µl buffer AL was mixed thoroughly to this mixture by vortexing, and incubated at 56°C for 10 min. Then 200 µl ethanol (96–100%) was added to the sample and mix thoroughly by vortexing and DNA were extracted using the Qiagen DNeasy blood and tissue kit, Qiagen Cat No./ID: 69504 following manufacturer's instructions. The study protocol was approved through Institutional Animal Ethics Committee (approval no. CPCSEA/IAEC/P-6/2018) which was formed in accordance with the norms of the Committee for the Purpose of Control and Supervision of Experiments on Animals (CPCSEA), Government of India and complied with standard guidelines on handling of experimental animals.

According to manufacturer's instructions, 20 µl Proteinase K was pipetted into a 1.5 ml or 2 ml microcentrifuge tube, to this 50–100 µl anticoagulated blood was added and the volume was adjusted to 220 µl with PBS. 200 µl Buffer AL was added and mixed thoroughly by vortexing, and incubated at 56°C for 10 min. Finally 200 µl ethanol (96–100%) was added to the sample, and mixed thoroughly by vortexing. The mixture was added to the DNeasy Mini spin column placed in a 2 ml collection tube and centrifuged at 6000 x g for 1 min. Further the DNeasy Mini spin column was placed in a new 2 ml collection tube (previous flow through and collection tube was discarded), 500 µl Buffer AW1 was added and centrifuged for 1 min at 6000 x g. Again flow-through and collection tube was discarded. The DNeasy Mini spin column was placed in a new 2 ml collection tube, 500 µl Buffer AW2 was added, and centrifuged for 3 min at 20,000 x g to dry the DNeasy membrane. Flow-through and collection tube was discarded. DNeasy Mini spin column was placed in a clean 1.5 ml or 2 ml microcentrifuge tube and 200 µl Buffer AE was pipetted directly onto the DNeasy membrane. Sample was incubated at room temperature for 1 min, and then centrifuged for 1 min at 6000 x g to elute.

**Dog [38]**

DNA samples from n = 742 dog blood samples from 93 breeds were provided by researchers at the National Human Genome Research Institute (NHGRI). The weight of individual dogs was unknown. Collection was approved by the Animal Care and Use Committee of the Intramural Program of NHGRI at the National Institutes of Health (Protocol #8329254).

**Dog lifespan and breed characteristics**

Standard breed weight (SBW), height (SBH) and lifespan were aggregated from several sources. SBW and SBH were taken from previously reported values [39,40], which were updated if AKC values differed [41]. If the AKC did not specify SBW or SBH, we used data from Atlas of Dog Breeds of the World [42]. Lifespan estimates were calculated as the average of the standard breed across sexes, compiled from numerous publications consisting primarily of multi-breed surveys of age and cause of death from veterinary clinics and large-scale breed-specific surveys, which are often conducted by purebred dog associations. Sources for lifespan are reported in Supplementary Note 1. When available, data was combined across surveys for number of dogs, minimum, maximum, mean, and median age at death. The minimums, maximums, and medians were averaged across studies to produce a representative lifespan expectation for each breed. For three breeds (American Hairless Terrier, Sloughi, and Ibizan Hound), no published survey data was available. For these breeds, the maximum age expectation was obtained from the American Kennel Club website.

**Bats [43]**

Wing tissue samples. Wing punches were taken from 778 individually marked animals that were either kept in captivity (15 species) or recaptured as part of long-term field studies (11 species). We excluded 42 samples because we did not have independent evidence to confirm minimum age estimates. For 630 samples the individual was marked shortly after birth, so age estimates were exact. For the remainder, age represented a minimum estimate because the individual was not initially banded as a juvenile. We used minimum age estimates when other evidence, such as tooth wear or time since initial capture, indicated that the minimum age estimate was likely to be close to the real age. The study was approved by the University of Maryland Institutional Animal Care and Use Committee (FR-APR-18-16).

Here, we provide additional information on when and where samples were taken from either captive or free-ranging animals for each of the 26 species of bats used in this study.

Pallid bats, *Antrozous pallidus,* were captured between 2005 and 2008 at six sites in central Oregon (44.94° N, 120.38° W) using mist nets over a water source or outside a night roost or with a handnet on an extension pole outside a day-roosting crevice. Each bat was weighed, measured and marked with a numbered band. Adults were distinguished from juveniles by closed epiphyseal gaps. Tissue samples were obtained from wing membranes using 3 mm biopsy punches and stored in 95% ethanol until DNA was extracted using a Qiagen DNeasy Tissue Kit. DNA extracts were stored frozen at -80°C. Live animal procedures conformed to the American Society of Mammalogists guidelines and were approved by the University of Maryland Institutional Animal Care and Use Committee (protocols R-05-26 and R-08-39). Bat capture and sampling was conducted with permission of the Pine Creek Conservation Area, the Oregon Department of Fish and Wildlife (permit 081-95), and the John Day Fossil Beds National Monument, National Park Service (permit JODA-2005-SCI-0003).

Wing tissue from Artibeus jamaicensis, Cynopterus brachyotis, Eidolon helvum, Pteropus giganteus, P. hypomelanus, P. poliocephalus, P. pumilus, P. rodricensis, P. vampyrus, and Rousettus aegyptiacus was taken between 2006 and 2017 from bats kept at the Lubee Bat Conservancy, an AZA (Associated Zoos and Aquariums [https://www.aza.org/current-cert]) certified facility, in Gainesville, Florida. The bats are group-housed in twelve 1068 sq. ft. enclosures with indoor temperature-controlled roosting areas and outdoor flight rooms and are fed a diet of fruit, vegetables and nutritional supplements. Wing tissue biopsies are periodically taken from individually marked animals and kept at -20°C in 95% ethanol. The majority of 243 samples from these species were taken from animals that were born in captivity. DNA was extracted with a Zymo miniprep plus kit.

Wing tissue samples were taken in 2018 from captive Carollia perspicillata housed in a tropical zoo (Papiliorama, Kerzers FR, Switzerland). Approximately 400 bats roost in an artificial cave kept on a reversed light cycle and are fed twice a night with a fruit-based diet. Since 2011, the population has been monitored by capturing individuals using a harp-trap placed at the entrance to the cave. Forearm length, body weight, reproductive status and tooth-wear are recorded from every captured individual. At first capture, individuals are marked on the forearms with a unique combination of three colored plastic rings (A.C. Hughes, UK, size XB). All captures and marking were authorized by the cantonal veterinary service (permits nb:2011_42_FR, 2013_10E_FR, 2014_59_FR). Between July and November 2018, 3mm biopsies were punched on the patagium and hermetically stored in silica gel. Based on the date of first capture and tooth-wear score, the age of each individual sampled was estimated[3]. DNA was extracted with a Zymo miniprep plus kit.

Biopsy punches (2 or 3 mm) were taken from the wing of captive common vampire bats, *Desmodus rotundus,* between 2010 and 2014. Bats were housed and fed blood[4] in flight cages (3 x 2 x 1.5 m) as a captive group at the Cranbrook Institute of Science (24-39 bats, Bloomfield Hills, MI, USA) or at the University of Maryland (7 bats, University of Maryland Institutional Animal Care and Use Committee protocol R-10–63). Age was determined based on zoo birth records. Individuals were born at the Houston Zoo, Cincinnati Zoo, Chicago Brookfield Zoo, or the Cranbrook Institute of Science. Tissue samples were stored in 95% ethanol prior to DNA extraction using a Qiagen DNeasy kit. DNA extracts were frozen for long-term storage at -80°C. Live animal procedures conformed to the American Society of Mammalogists guidelines and were approved by the University of Maryland Institutional Animal Care and Use Committee (protocol R-10-63).

Wing tissues of big brown bats (*Eptesicus fuscus*) were sampled with a 3 mm biopsy punch from the wing of known age captive bats at Northeast Ohio Medical University (NEOMED; Rootstown, Ohio) in 2018 and 2019. These animals came from a colony previously maintained by Dr. Ellen Covey at the University of Washington, which was started in 2005 with bats caught in North Carolina that were banded according to year of capture or birth. These bats underwent natural hibernation and were exclusively fed an *ab libitum* diet of fresh water and mealworms (*Tenebrio molitor*). In 2014 some of the bats were transported to NEOMED and are now housed indoors on a 12 h light/dark cycle and fed the same *ab libitum* fresh water and mealworm diet. Wing punches were taken from a second colony of *E. fuscus* by Dr. Paul Faure and Lucas Greville in February or August, 2020. This colony is kept at McMaster University in Hamilton, Ontario, Canada in an indoor/outdoor enclosure (2.5 x 8.3 x 2.7m)[7] in which the temperature fluctuates with ambient conditions, but is kept above freezing by a heater on a thermastat. Known-age animals from this colony were born in captivity from females captured in Ontario. Tissue samples from both colonies were stored in DNA Shield and kept frozen at -20°C prior to DNA extraction with a Zymo miniprep plus kit. Animal use protocols were approved by the NEOMED

Institutional Animal Care and Use Committee or the Animal Research Ethics Board of McMaster University (AUP# 20-05-20).

During July 2019, wing membrane samples were obtained from subadult or adult female and subadult male lesser long-nosed bats, *Leptonycteris yerbabuenae,* with a 4 mm biopsy punch at the entrance of the Pinacate Cave in the Reserva de la Biosfera el Pinacate y Gran Desierto de Altar (31°38'51.6" N, 113°28'53.5" W), Sonora, Mexico. Bats were captured using mist nets (Avinet models: TB02, TB06, TB012; Portland, Maine, USA) set outside caves just prior to when bats emerged to forage. Individuals were sexed, weighed and the forearm measured. To discriminate subadults from adults, age was determined by the degree of fusion of the epiphyses at the metacarpal–phalangeal joint. Tissue samples were stored in DNA/RNA Shield buffer (Zymo Scientific, Irvine, CA 92614, U.S.A.). DNA was extracted with a Zymo miniprep plus kit. Bat tissue samples were collected under permit SGPA/DGVS/06361/17 issued to R. A. Medellín by The Ministry of Environment and Natural Resources.

Samples of velvet free-tailed bats, *Molossus molossus*, come from a long-term study in Gamboa, Panama (09°07’ N 79°41’ W), where the bats roost in crevices in houses. We captured social groups with mist nets (Ecotone, Gydnia, Poland) at the entrance of roosts during evening emergence and individually marked all bats with a subcutaneous passive integrated transponder (Trovan ID-100, Euro ID, Weilerswist, Germany) at first capture. Wing tissue samples were taken with a 3 mm biopsy punch and stored in 96% ethanol until DNA extraction using a Zymo miniprep kit. Capture and handling of animals were carried out under permits SE/A-112-13, SE/A-73-14, SE/A-95-15, and SE/A-32-17 from the Autoridad Nacional del Ambiente in Panama with approval from the Institutional Animal Care and Use Committee of the Smithsonian Tropical Research Institute (2012-0505-2015).

Little brown bats, *Myotis lucifugus*, were captured as they departed from an attic maternity colony in Chestertown, Maryland (39°12’N, 76°04’W), in September 1996. Captured bats were weighed, measured and banded with individually marked bands. Young of the year were identified by their weight and absence of tooth wear. Wing membrane biopsies were taken and stored in a 5M NaCl with 20% dimethyl sulfoxide solution and kept frozen at -80°C. DNA was extracted with a Zymo miniprep plus kit. Bat capture and handling was approved by the Maryland Department of Natural Resources (permit SCO-30403).

Wing tissue samples were taken from greater mouse-eared bats, *Myotis myotis*, between 2013 and 2018 as part of a long-term mark-recapture study conducted by Bretagne Vivante in Brittany, France (47°35'N, 2°14'W"). Bats were caught using modified harp traps as they left one of five different roosts. Individuals at first capture are fitted with PIT tags to facilitate identification on subsequent recaptures. Measurements taken from each individual include sex, forearm length, weight and transponder number. Age class (juvenile or adult) is determined by examining the degree of the epiphyseal closure of the metacarpal-phalangeal joints. Wing biopsies were taken with a 3 mm biopsy punch, flash frozen and stored in liquid nitrogen prior to extraction. All procedures were conducted with full ethical approval and permission (AREC-13- 38-Teeling) awarded by the University College Dublin ethics committee and in accordance with permits issued by 'Arrêté' by the Préfet du Morbihan. DNA was extracted from wing biopsies using a Promega Wizard SV DNA extraction kit (catalog no. A2371) or the Qiagen DNeasy Blood and Tissue kit (Qiagen). Extractions carried out with the Promega kit were partially automated using a Hamilton STAR Deck liquid handling robot.

Adult and juvenile Mexican fishing bats, *Myotis vivesi*, were captured by gloved hand from roosts in talus slopes on Isla Partida Norte in the Gulf of California, Mexico (29°03'N, 113°00'W) during the day between 2015 and 2018. Individuals were measured and banded with numbered metal bands on their forearms for identification upon recapture. In 2018 wing tissue was taken with a 3 mm biopsy punch and preserved in Zymo DNA shield. Bat capture and handling were conducted under permits #7668–15, 2492–17 and #5409-18 from Dirección General de Vida Silvestre, and permits #17–16, 21–17 and 20-18 from Secretaría de Gobernación, and the University of Maryland Institutional Animal Care and Use Committee protocols FR-15-10 and FR-18-20.

Common noctules, *Nyctalus noctula*, were captured as part of a long-term study[16,17,18] at the Seeburgpark in Kreuzlingen, Switzerland (47.649928° N, 9.186123° E) where bats regularly roost in boxes. Each bat was marked with a subcutaneous pit-tag (ID100; Euro ID, Weilerswist, Germany) injected under the dorsal skin. Wing tissue samples were taken with a 3 mm biopsy punch and stored in 96% ethanol until DNA extraction using a Zymo miniprep kit. All handling and sampling of the bats in Switzerland was approved by the Veterinäramt Thurgau (permit FIBL1/12).

Wing tissue samples were taken in September 2018 from lesser spear-nosed bats, *Phyllostomus discolor*, kept in a breeding colony in the Department Biology II of the Ludwig- Maximilians-University in Munich. In this colony animals were kept under semi-natural conditions (12 h day/night cycle, 65 to 70 % relative humidity, 28°C) with free access to food and water. The license to keep and breed *P. discolor* was issued by the German Regierung von Oberbayern. Under German Law on Animal Protection a special ethical approval is not needed for wing tissue collection. Wing tissue was stored in RNAlater until DNA was extracted using QIAamp® MinElute columns following the manufacturer's instructions. The samples were eluted in 50µl of molecular grade water and concentrated to reduce their volume by approximately 50% using a Speedvac, with the following settings: duration 20 minutes, temperature in the chamber 30°C, $H_2O$ (water) mode.

Greater spear-nosed bats, *Phyllostomus hastatus,* were captured and sampled between 1990 and 2018 in Trinidad, Lesser Antilles[19,20,21,22]. Most often, harem groups, which include one adult male plus 15-20 lactating females with pups, were captured during the day from within a solution depression in the ceiling of either Tamana (10.4711°N, 61.1958°W), Caura (10.7019°N, 61.3614°W), or Guanapo cave (10.6942°N, 61.2654°W) using a bucket trap. Captured bats were sexed, measured for size, weight, and tooth wear, and individually marked with stainless steel numbered bands. Age was determined exactly for adults that were recaptured after being banded as pups. Wing biopsy punches (4 mm) were stored frozen at -80°C in either a 5M NaCl with 20% dimethyl sulfoxide solution or Zymo DNA Shield prior to DNA extraction using a Qiagen Puregene or Zymo miniprep plus kit. Frozen samples were selected to maximize the number of known-age individuals with approximately equal numbers at all ages. Animal handling methods follow guidelines by the American Society of Mammalogists and were approved by the University of Maryland Institutional Animal Care and Use Committee (protocols R-91-33, R-93-22, R-94-25, R-01-07, R-11-21, R-13-77, FR-APR-18-16) under licenses from the Forestry Division of the Ministry of Agriculture, Land and Fisheries, Trinidad and Tobago.

Wing tissue samples were taken from greater horseshoe bats, *Rhinolophus ferrumequinum*, by using 3 mm biopsy punches between 2016 and 2018 from wild female bats as part of a long-term study at a maternity colony in Gloucestershire, UK (51.7107°N, 2.2777°W). Bats were captured at the roost with hand nets, and all individuals were weighed, ringed with aluminum alloy rings and morphometric data such as forearm length recorded under licenses (Natural England Project Licences 2015-9918-SCI-SCI; 2016-23583- SCI-SCI; 2017-30137-SCI-SCI) issued to Roger Ransome. All bats studied were first marked as infants, so we could be certain of their age. Bats were aged between 1-21 years, with the 40 individuals selected in a fairly even manner across this age span. Sampling procedures were conducted under licenses (Natural England 2015-11974-SCI-SCI; 2016-25216-SCI-SCI; 2017- 31148-SCI-SCI) issued to Gareth Jones, with tissue biopsy additionally licensed under Home Office Project Licenses (PPL 30/3025 prior to 2018; P307F1428 from 2018 onwards) and Home

Tissue samples were stored in silica gel beads and then transferred to a -20C freezer for long-term storage. DNA was extracted from wing biopsies using a Promega

Wizard SV DNA extraction kit or a Qiagen DNeasy Blood and Tissue kit. Extractions carried out with the Promega kit were partially automated using a Hamilton STAR Deck liquid handling robot but otherwise followed manufacturer's instructions.

Samples of proboscis bats, *Rhynchonycteris naso,* came from a long-term study between 2005 and 2016 at La Selva Biological Station in Costa Rica (10° 25´ N, 84° 00´W)[26,27,28]. Bats were mist-netted in the vicinity of their roosts. Wing tissue was sampled with a 4 mm biopsy punch, individuals were marked with colored plastic bands, sexed, measured and age class determined (juvenile: 0-4 months, subadult: 5-10 months, or adult>10 months)[26]. Age was determined exactly for individuals that were banded as pups and recaptured as adults. Ethanol (80%) was used to preserve tissue samples, and a salt–chloroform procedure or Qiagen BioSprint 96 DNA Blood Kit was used for DNA isolation[26,28]. Research permits were granted by the MINAE (Ministerio del Ambiente y Energia) and the ACC (Área de Conservación Central). Animal treatment followed the Guide for Care and Use of Laboratory Animals of the National Institutes of Health. Animal handling complied with current Costa Rican laws and was approved by the Animal Care Review Committee of the SINAC (Sistema Nacional de Áreas de Conservación) and MINAE (permits 022-2005-OFAU, 108-2006-SINAC, 147-2007-SINAC,

Office personal licences.

183-2008-SINAC, 187-2009-SINAC, 130-2010-SINAC and 068-2011-SINAC, 115–2012- SINAC, 033–2013-SINAC, SINAC-SE-GASP-PI-R-121–2013, R-006–2015-OT-CONAGEBIO, SINAC-SE-CUS-PI-R-088–2016). Samples of greater sac-winged bats, *Saccopteryx bilineata* came from long-term studies in Costa Rica (n = 21 from La Selva Biological Station, 10° 25´ N, 84° 00´W and n = 6 from Santa Rosa National Park, 10° 53′ N, 85° 46′ W) and Panama (n = 4 from the Biological Station Barro Colorado Island (BCI) of the Smithsonian Tropical Research Institute, 9° 9′ N/79° 51′ W) between 1994 and 2016. Bats were captured with mist nets when entering or leaving their day roosts, individually banded with two coloured plastic bands on their forearms, and a wing tissue biopsy sample (4mm) preserved in 80% ethanol was taken. Age was determined exactly for individuals that were banded as pups and recaptured as adults. DNA was extracted with a salt-chloroform procedure or with the Qiagen BioSprint 96 DNA Blood Kit. The process of acquiring data and protocols for capturing and handling bats complied with the current laws of Panama and were conducted in accordance with the relevant guidelines and regulations. Our study in Panama was approved by the Smithsonian Tropical Research Institute and its Animal Care and Use Committee (ACUC, permits: IACUC 100316-0910-12, ACUC 2013-1015-2016). For research in Costa Rica, permits were granted by the MINAE (Ministerio del Ambiente y Energia), the ACC (Área de Conservación Central) and the ACG (Área de Conservación Guanacaste). Animal treatment followed the Guide for Care and Use of Laboratory Animals of the National Institutes of Health. Animal handling complied with current Costa Rican laws and was approved by the Animal Care Review Committee of the SINAC (Sistema Nacional de Áreas de Conservación) and MINAE (permits 272-2003-OFAU, 135-2004-OFAU, 022-2005-OFAU, 108-2006-SINAC, 147-2007-SINAC, 183-2008-SINAC, 187-2009-SINAC, 130-2010-SINAC and 068-2011-SINAC, 115–2012-SINAC, 033–2013-SINAC, SINAC-SE-GASP-PI-R-121– 2013, R-006–2015-OT-CONAGEBIO, SINAC-SE-CUS-PI-R-088–2016).

Mexican free-tailed bats, *Tadarida brasiliensis*, are housed at Bat World Sanctuary, a licensed non-profit bat rehabilitation facility and accredited by the Global Federation of Animal Sanctuaries (https://www.sanctuaryfederation.org/sanctuaries/bat-world/) in Weatherford, Texas. Most individuals sampled were rescued as pups, although some were rescued as adults, making their exact age unknown. Individuals are group housed in large indoor enclosures. Wing membrane biopsies (4 mm) were collected by Amanda Lollar in August 2019 and stored in Zymo DNA Shield until DNA was extracted using a Zymo miniprep plus kit.

**Cattle [44]**

**Ethical authorization and animals**

All animal procedures were carried out in accordance with the relevant guidelines at each institution. Specifically, procedures related to sample collection in Poland followed the EU Directive of the European Parliament and the Council on the protection of animals used for scientific purposes (22 September 2010; No 2010/63/EU), Polish Parliament Act on Animal Protection (21 August 1997, Dz.U. 1997 nr 111 poz. 724) with further novelization - Polish Parliament Act on the protection of animals used for scientific or educational purposes (15 January 2015, Dz.U. 2015 poz. 266). Blood and oocyte collection were approved by the Local Ethics Committee for Experiments on Animals, University of Warmia and Mazury in Olsztyn, Poland (Agreement No. LKE.065.27.2019). For animal procedures in the USA, approval from the University of Nebraska Institutional Animal Care and Use Committee was obtained (approval number is 1560). Samples were obtained from 357 female cattle (Bos Taurus). The animals were housed on the Dairy Farm of the Institute of Animal Reproduction and Food Research of Polish Academy of Sciences, (Wielki Las, Poland) and at the Eastern Nebraska Research and Extension Center at the University of Nebraska-Lincoln (Nebraska, USA). In the present study, samples were collected from Polish Red cattle and from the herd in Eastern Nebraska. In this herd, black Angus or composites of varying percentages of Simmental x Angus (black) or Red Angus where used as blood donors. Animals were free of Bovine Herpesvirus Type 1, Bovine Viral Diarrhea/Mucosal Disease, tuberculosis and Enzootic bovine leucosis.

**Blood collection and further processing**

The blood samples from both herds were collected during routine animal management activities, during the routine blood collection for disease prevention. Blood samples were taken only from cows in the luteal phase of the estrous cycle. Blood was collected into 8ml PAXgene Blood DNA Tubes (Quiagen, Cat No. 761115) and stored at -80℃ until the shipment (USA Veterinary Permission Nr 138809) to the UCLA Technology Center for Genomics & Bioinformatics (Los Angeles, USA) for further analyses.

**Oocyte Collection and further processing**

In total 80 Bovine ovaries were collected immediately *post mortem* from 40 cows which were selected for routine culling due to management reasons. All cows were in the luteal phase of the estrous cycle. Before the isolation of both ovaries from each cow, blood samples were also collected into 8ml PAXgene Blood DNA Tubes (Quiagen, Cat No. 761115) to generate both sample types from one donor. Isolated ovaries were kept on ice and immediately transported to the laboratory. Afterwards, immature bovine cumulus–oocyte complexes (COCs) were recovered by aspirating ovarian follicles in the diameter of 2–8 mm. Cumulus cells were removed from COCs by pipetting them for 5min in a Petri dish containing 500 µl of Phosphate Buffered Saline (PBS) with 0.1% hyaluronidase (Sigma-Aldrich, St. Louis, MO, USA). Denuded oocytes from every donor were pooled (10-15 immature oocytes/ovary) and then processed for genomic DNA isolation.

**Mouse data** [45]

The mouse data come from several sources listed below.

**UCLA Lab. Animal breeding and husbandry** [46]

All mice were maintained and bred under standard conditions consistent with National Institutes of Health guidelines and approved by the University of California, Los Angeles Institutional Animal Care and Use Committees. The cages were maintained on a 12:12 light/dark cycle, with food and water ad lib. Tissues were harvested from discervical mice and fresh frozen on dry ice.

The Tet3 knockout mice were generated at the Jackson Laboratory using CRISPR technology with deletion of exon 4. Tet3 heterozygous knockout mice were crossed with C57BL/6J (JAX 000664) to set up the colony. All mice were maintained and bred under standard conditions consistent with National Institutes of Health guidelines and approved by the University of California, Los Angeles Institutional Animal Care and Use Committees. The cages were maintained on a 12:12 light/dark cycle, with food and water ad lib. Mice were euthanized by cervical dislocation,  cortical and striatal tissues were then quickly dissected out and fresh frozen on dry ice until used for DNA extraction.

Bcl11b heterozygous mouse was obtained from MMRRC at University of California, Davis (Catalog # 046780-UCD) and crossed with C57BL/6J (JAX 000664) to set up the colony. All mice were maintained and bred under standard conditions consistent with National Institutes of Health guidelines and approved by the University of California, Los Angeles Institutional Animal Care and Use Committees. The cages were maintained on a 12:12 light/dark cycle, with food and water ad lib. Mice were euthanized by cervical dislocation,  cortical and tissues were then quickly dissected out and fresh frozen on dry ice until used for DNA extraction.

**BXD mice [45] University of Tennessee Health Science Center**

Samples for this study were selected from a larger colony of BXD mice that were housed in a specific pathogen-free (SPF) facility at the University of Tennessee Health Science Center (UTHSC). All animal procedures were in accordance to protocol approved by the Institutional Animal Care and Use Committee (IACUC) at the University of Tennessee Health Science Center. Detailed description of housing conditions and diet can be found in (Roy, 2020;Williams, 2020). Mice were given ad libitum access to water, and either standard laboratory chow (Harlan Teklad; 2018, 18.6% protein, 6.2% fat, 75.2% carbohydrates), or high-fat chow (Harlan Teklad 06414; 18.4% protein, 60.3% fat, 21.3% carbohydrates). Animals were first weighed within the first few days of assignment to either diets, and this was mostly but not always prior to introduction to HFD. Following this, animals were weighed periodically, and a final time (BWF) when animals were humanely euthanized (anesthetized with avertin at 0.02 ml per g of weight, followed by perfusion with phosphate-buffered saline) at specific ages for tissue collection. The present work utilizes the biobanked liver specimens that were pulverized and stored in -80 °C, and overlaps samples described in (Williams, 2020).

**Growth hormone receptor knockout from the University of Michigan**

Growth hormone receptor knockout mice (GHR -/- dwarfs) from the University of Michigan (Richard A. Miller). Full-body growth hormone receptor knockout (GHR-KO) [47].

**Calorie restricted mice from the University of Texas Southwestern Medical Center [48]**

##### *Ethics*

The Institutional Animal Care and Use Committee (IACUC) of the University of Texas Southwestern Medical Center approved the animal protocol (APN 2015-100925), which has been subsequently renewed every 3 years (2018 and 2021).

##### *Animals*

C57BL/6J male mice (6 week-old, n = 72) were obtained from the Mouse Breeding Core, Wakeland lab, UT Southwestern Medical Center, Dallas, TX, USA. After 2 weeks of acclimation, mice were individually housed in standard polycarbonate mouse cages (Fischer Scientific, Cat. Nos. 01-288-1B and 01-288-21) with stainless steel running wheel, inside isolation cabinets containing 12 cages each. Temperature and humidity levels were monitored, and the mice were housed under light/dark cycles of LD12:12 h (green LEDs, ~100 lux at the level of the cage floor). Water was provided ad libitum throughout the study.

Mice were fed with round pellets of 300 mg each containing 3.60 Kcal/g (Dustless Precision Pelletsâ, Rodent, Purified, F0075, BioServ, Flemington, NJ, USA). The food composition is similar to regular mouse chow (18.7% protein, 5.6% fat, 59.1% carbohydrates and 4.7% fiber). Food access was controlled by an automated feeder system designed in our lab with Phenome Technologies Inc., Skokie, IL, USA that precisely controls how much, when, and how often the food is dispensed (Acosta-Rodríguez et al., Cell Metabolism 2017). After six weeks of recording under ad libitum food access, 12 mice/group were randomly assigned to one of 6 feeding conditions (Acosta-Rodriguez et al., under revision): 24h access ad libitum; 70% of baseline ad libitum levels fed at the beginning of the dark (CR-night) or light (CR-day) phase which they consumed in less than 2h (classic CR protocols); or fed under CR evenly spread over 12 hours during the dark (CR-night-12h) or light (CR-day-12h) by releasing one 300 mg pellet every 90 min; or evenly spread over 24 hours (CR-spread) by releasing one 300 mg pellet every 160 min. Mice were weighed during cage change every 21 days, and bedding was checked for any evidence of food spillage. ClockLab Chamber Control Software v3.401 (Actimetrics Inc., Wilmette, IL, USA) was used to schedule and record feeding events, and ClockLab Data Acquisition System v3.209 (Actimetrics Inc., Wilmette, IL, USA) was used to record wheel-running behavior.

##### *Tissue collection and DNA extraction*

At 19 months of age, mice were released into constant darkness and samples collected every 4 hours for 48 hours (n=12 mice/condition). Tissues were snap frozen and stored at -80°C until DNA extraction.
Liver tissue was digested using 1mL of digestion buffer (100mM NaCl, 50mM Tris-HCl pH 8, 100mM EDTA pH 8 and 1% SDS) plus 20uL of protein K (20mg/mL) incubated ON at 60ºC. After a RNAse (Ambion, AM2286, 2uL) treatment for 1h at 37ºC, DNA extraction was performed using UltraPure™ Phenol:Chloroform:Isoamyl Alcohol (25:24:1, v/v, Thermo Fisher, #15593049) and phase lock gel (VWR, # 10847-802) to avoid contamination with interphase, according to manufacturer instructions.

#### South African species

##### *Ethics*

Both studies were originally approved through the Animal Ethics Committee, University of Pretoria, South Africa, under AUCC project number: EC037-15.
The Animal Use and Care Committee of the University of Pretoria evaluated and approved the experimental protocol (ethics clearance number: NAS022/2021, NAS209/2021, NAS021/2020, ec028-18, ec069–17, ec015–08) and DAFF section 20 approval (SDAH-Epi-21021810221, SDAH-Epi-20072707050, 12/11/1/1/8 (2002 LH)). Permission to capture was obtained from all landowners, and a collecting permit was obtained from the relevant nature conservation authorities (Permit number: CPF6 Western Cape- CN44-87-13780, CN44-31-2285, Northern Cape – FAUNA 0715/2020, Gauteng- CPF6-0124, Kwa-Zulu Natal- OP1545/2021).

##### *Study animals*

Tissues were from archived post mortem samples stored at -80ºC, from Fukomys damarensis collected from Winterton, Northern Cape with permission from Department of Environment and Nature Conservation,

Northern Cape Province. Georychus capensis samples were collected from Darling, Weestern Cape with permission from the Western Cape Nature Conservation.

Except for Heterocephalus glaber, Fukomys damarensis, Atelerix albiventris and Echinops telfairi, which came from captive-raised populations, all other small mammal species were wild-caught in South Africa. All terrestrial small mammal species, Crocidura cyanea and Elephantulus myurus, were caught using Sherman traps baited with oats and peanut butter. While all small subterranean mammals, Georychus capensis, Bathyergus suillus, Cryptomys hottentotus hottentotus, C. h. pretoria, C. h. mahali, C. h. natalenesis and Amblysomus hottentotus were captured using Hickman live traps, baited with a small piece of sweet potato. All traps were monitored for captures every 2-3 hours over the course of the day and left overnight, being checked first thing in the morning.

**Apodemus mice**

Apodemus mice were culled during routine (yearly) population control, via cervical dislocation confirmed by exsanguination. All animal work was conducted in accordance with the UK Home Office in compliance with the Animals (Scientific Procedures) Act 1986, was approved by the University of Edinburgh Ethical Review Committee and was carried out under the approved UK Home Office Project License PP4913586.

49

Prof Thomas Little maintains a formerly-wild, but now lab-reared wood mouse colony in standard laboratory conditions at the University of Edinburgh. The colony has been in captivity for many generations, but the wood mice are purposely outbred to maintain genetic diversity. All mice are housed individually in ventilated cages (Techniplast, 1285L) with food and water ad libitum. DNA was extracted using a phenol-chloroform method from ear punches taken from individuals of known age from the colony. We obtained DNA from 48 mice of both sexes, spanning an age range of 88 to 496 days old. This slightly unusual choice of ages arises because we utilised mice that were part of other experiments. Lifespan in wild mice is not known for certain, and many will die from extrinsic mortality (e.g. predation), but in our own field work we recapture 10-20% of tagged mice the following year (Pedersen, unpublished data), meaning wild mice may can live for hundreds of days.

**Spiny mouse**

Following anesthesia with 4% (v/v) vaporized isoflurane (Henry Schein Animal Health, Dublin, OH), full thickness ear tissue biopsies were collected from spiny mice (Acomys cahirinus) housed in breeding colonies at the University of Kentucky, Lexington, KY or Monash University, Melbourne, Australia. Biopsies were used for ear tissue samples and to produce ear pinna fibroblast cultures. Animal protocols were approved by the Institutional Animal Care and Use Committee (IACUC) at the University of Kentucky (2019-3254). Spiny mouse fetal fibroblasts were derived from embryonic skin following enzymatic digestion with trypsin. Connective tissue fibroblasts from adult spiny mice were derived from ear biopsies and grown at 37 °C in 3% oxygen and 5% CO2 as previously described (Saxena et al., 2019).

**Nova Scotia masked shrews (Sorex cinereus) [50]**

Sampling, sexing, and aging shrews

Masked shrew samples were collected from five locations: Peterborough County, Ontario and Sandy Cove, North Mountain, Long Island and Bon Portage Island (BPI) in Nova Scotia, Canada. All shrews were trapped within six weeks of each other to limit seasonal morphological changes, such as skull and body size (Lázaro et al., 2019, 2021). DNA was extracted from liver, tail, and fetus using the DNeasy Blood & Tissue Kit from QIAGEN. Shrews were left in dermestid beetle tanks for a week to remove all remaining tissue from the skulls. A Leica EZ4 microscope was used to assess age class for each shrew based on teeth wear according to Pruitt’s method(Pruitt, 1954; Rudd, 1955). Age in months was estimated based on the trapping date, the age class, and the known reproductive season of masked shrews assuming a maximum lifespan of ~17 months (Churchfield, 1990). Shrews were sexed by PCR amplification of the SRY gene (Cervantes et al., 2013; Matsubara et al., 2001) using custom primers (Table S1). Gel electrophoresis was used to identify successful amplification and thus confirm the presence of the Y chromosome (i.e. male). Age class designations were confirmed based on morphological features(Pruitt, 1954; Rudd, 1955); sex assignments were subsequently validated with the methylation assay(Wang et al., 2021).

**Shrews and small animals from Museum of Biological Diversity at The Ohio State University.**

**Ethics statement**

All wild-caught animals were collected and sacrificed in accordance with protocols approved by The Ohio State University IACUC (Institutional Animal Care and Use Committee) under protocol number 2017A00000036. All wild-caught animals were collected with scientific collecting permits issued from Ohio and Washington and according to guidelines established by the American Society of Mammalogy for the use of wild animals in research (Sikes & Animal Care and Use Committee of the American Society of Mammalogists 2016).

Tissue samples were taken from archived specimens that are in the process of being deposited in the Museum of Biological Diversity at The Ohio State University.

Tissue samples were acquired from wild animals using multiple collecting procedures, including (1) trapping live animals using Sherman live-traps or pitfalls and followed by euthanasia using isoflurane overdose, (2) sampling of roadkill animals, and (3) sampling of deceased animals received from wildlife rehabilitation centers. Abdominal and pelvic organs were removed from animals and archived in a -80C freezer.

**Marsupials [51]**

**Opossum and mouse tissue samples**

Opossum samples come from a pedigreed, breeding colony of gray short-tailed opossums (Monodelphis domestica) that was established by founder individuals purchased from the Southwest Foundation for Biomedical Research. Mouse samples come from a breeding colony of C57BL/6J mice originally purchased from the Jackson Laboratory. Both colonies are maintained by the Sears Lab at UCLA. Opossums were euthanized by CO2 inhalation to effect followed by bilateral thoracotomy. Mice were euthanized by CO2 inhalation to effect followed by cervical dislocation. These procedures are in accordance with the AVMA Guidelines for the Euthanasia of Animals 2013: https://www.avma.org/KB/Policies/Documents/euthanasia.pdf, and all animal procedures were approved by the UCLA IACUC. All tissue samples, e.g., liver, blood, tail, were taken from euthanized opossums and mice and stored at -20 C until use. DNA from samples was extracted and purified using a DNA Miniprep Plus Kit (Zymo), following manufacturer’s protocols. PicoGreen fluorescent dsDNA was used to assess the concentration of resulting DNA and concentrations adjusted to 50 to 250 ng/ul. DNA samples were submitted to the Technology Center for Genomics & Bioinformatics at UCLA for generation of DNA methylation data and further analyses.

**Tasmanian devil tissue samples [51]**

Ear samples from Tasmanian devil (Sarcophilus harrisii) were collected from individuals in the Tasmanian devil insurance metapopulation, either living as an introduced population on Maria Island (N=29), or in the zoo-based population (N=17). Samples were collected by the Save the Tasmanian Devil Program (STDP), or the respective zoos, under the STDP's Standard Operating Procedure: Trapping and handling wild Tasmanian devils and shared with the University of Sydney. The samples were collected for standard management practice, which received ethics approval over 15 years ago. DNA from these samples were made available for use in this study, no individuals were specifically sampled for the purposes of this research.

Tasmanian devils within the zoo-based insurance population are housed in a range of scenarios from intensive housing as individuals, or as pairs in breeding season; or in large group housed enclosures, ranging from 4-10 males and 4-10 females per enclosure [52]. Tasmanian devils are not known to be a social species and housing within the Tasmanian devil insurance population is under the management of the Zoo and Aquarium Association Australasia [52]. Devils are housed and fed according a variety of diet items and managed according to the ZAA Husbandry Guidelines for the species [53]. Devils living on Maria Island are wild devils and so there is no animal care or maintenance for these individuals other than twice yearly monitoring trapping trips.

Samples were selected for this study to include a range of known age individuals and sexes. DNA was extracted using either a modified phenol-chloroform protocol [54] or the MagAttract HMW DNA kit (Qiagen, Germany; cat: 67563). DNA concentration and quality were assessed using a Nanodrop 2000 Spectrophotometer (ThermoFisher Scientific) and 0.8% agarose gel electrophoresis for 30 minutes at 90V.

**Kangaroo and wallaby tissue samples [51]**

The blood samples for kangaroo and wallaby were opportunistically collected from zoo-based animals during routine health exams. We analyzed blood from the following species: Macropus rufus (red kangaroo), Macropus giganteus (Eastern grey kangaroo), Macropus fuliginosus (Western grey kangaroo), and Macropus rufogriseus (red-necked wallaby).

**Mammalian liver samples (Diego Villar Lozano and Duncan Odom)**

**Ethics**

The use of all animals in this study was approved by the Animal Welfare and Ethics Review Board, under reference number NRWF-DO-02vs, and followed the Cancer Research UK Cambridge Institute guidelines for the use of animals in experimental studies. Tissue samples from humans were obtained from the Addenbrooke's Hospital at the University of Cambridge, under license number 08-H0308-117 ''Liver specific transcriptional regulation''.

**Methods**

We obtained mammalian tissue samples from routine euthanasia procedures (e.g. macaque, marmoset, rabbit, cat, dog, horse, and opossum), commercial providers or abattoirs (e.g. cow, pig, ferret, and guinea pig), and research tissue banks from specialty conservation programmes (e.g. cetaceans, Damaraland mole-rat and tissue samples from zoo post-mortem examinations). Tissue sources of these samples have been reported in previous publications [55-57]. In most cases, tissues were obtained immediately post-mortem (typically within an hour) to maximize experimental quality. Tissues were kept on ice until processed to minimize potential loss of integrity during post-mortem time, and flash-frozen in dry ice. 20-50 mg of tissue was used for extraction of genomic DNA with standard commercial kits (Qiagen), and genomic DNA samples were plated in 96-well plates using a randomised layout obtained with the R package OSAT [58].

**Mammalian liver samples from the University of Rochester**

All experiments were performed according to procedures approved by the University of Rochester Committee on Animal Resources (UCAR). Animal protocol # 101939 / UCAR-2017-033

The tissues were from the Gorbunova and Seluanov tissue bank at the University of Rochester [59,60].

**Supplementary Note 2: List of GEO accession**

All data from the Mammalian Methylation Consortium will be posted on Gene Expression Omnibus (GEO).

Below, we list the GEO accession number (GEO), Title, and sample size (N).

| GEO | Title | N |
|---|---|---|
| GSE174758 | Methylation studies in prairie voles | 144 |
| GSE184211 | Methylation studies in multiple human tissues N16 | 661 |
| GSE184213 | Methylation studies in multiple human tissues N36 | 96 |
| GSE184215 | Methylation studies in multiple human tissues N47 | 95 |
| GSE184216 | Methylation studies in roe deer | 94 |
| GSE184218 | Methylation studies in multiple human tissues N66 | 92 |
| GSE184220 | Methylation studies in human and mouse placenta | 47 |
| GSE184221 | Methylation studies in human blood N94 N68 | 384 |
| GSE184224 | Methylation studies in human cells N76 | 17 |
| GSE190660 | Methylation studies in the vervet monkey | 240 |
| GSE190661 | Methylation studies in baboons. | 325 |
| GSE190662 | Methylation studies in strepsirrhini primates | 91 |
| GSE190663 | Methylation studies in common marmoset | 96 |
| GSE190664 | Methylation studies in rhesus macaque | 283 |
| GSE174544 | Methylation studies in yellow-bellied marmots | 159 |
| GSE190665 | In vivo partial reprogramming in 4F mice | 168 |
| GSE174767 | Methylation studies in horses | 333 |
| GSE184222 | Methylation studies in wild ass and Grevy's zebra | 12 |
| GSE184223 | Methylation studies in plains zebras | 118 |
| GSE174777 | Methylation studies in naked mole rat. Part 1 | 289 |
| GSE174778 | Methylation studies in mole rats. Part 2 | 94 |
| GSE173330 | Methylation clocks for age estimation in toothed whales and dolphins | 545 |
| GSE164127 | Genome Methylation Predicts Age and Longevity of Bats | 908 |
| GSE147002 | DNA methylation profiles from a mouse model of Huntington's disease | 72 |
| GSE147003 | DNA methylation profiles from a transgenic sheep model of Huntington's disease | 168 |
| GSE147004 | DNA Methylation Study of Huntington's Disease and Motor Progression in Three Species | 348 |

**Supplementary Note 3: Sensitivity analysis of enrichment results**

It is critical to use a suitable background when it comes to any gene/pathway enrichment study. The wrong choice of background could easily lead to erroneous but highly significant associations due to hidden biases. When it comes to the mammalian array, the choice of the proper background must reflect the following sources of bias. First, limited genome coverage provided by the 37,000 CpGs on the array. For example, the CpGs on the mammalian array cover 6,871 human and 5,659 mouse genes when each CpGs is assigned uniquely to its closest gene neighbor. Second, by design, the mammalian array is biased toward highly conserved genomic regions. To address these biases, we evaluated the GREAT analysis software tool. As illustrated below, we find that GREAT analysis effectively deals with these biases and leads to biologically meaningful insights. In the following, we will report results from two different sensitivity analyses that were inspired by our GREAT enrichment analysis of the top 1,000 age-related CpGs (EWAS of age). Our first sensitivity analysis involved a random set of 1,000 mammalian CpGs. In essence, this evaluates the null hypothesis of no relationship between chronological age and methylation. The most significant (nominal) enrichment p-value was $P$=3.9x10$^{-4}$. Note that this p-value is far less significant than the enrichment p-values for age-related CpGs in our article: top 1,000 negative CpGs lead to $P$=2.7x10$^{-8}$; top 1,000 positive age-related CpGs lead to $P$=2.7x10$^{-266}$. We repeated this analysis with several sets of random 1,000 CpGs and obtained similar results.

Second, we also evaluated the enrichment of the top 1,087 most highly conserved CpGs across 158 mammalian genomes. This sensitivity analysis addresses the concern that highly conserved CpGs could have an increased chance of correlating strongly with chronological age or, conversely, non-conserved (noise) CpGs are expected to have no signal for age and will therefore not be selected in an EWAS of age. This hidden bias would manifest itself as follows: the enrichment analysis of our meta-analysis EWAS for age would be equivalent to the EWAS of highly conserved CpGs. In the following, we provide details that demonstrate that this is not the case. This biologically meaningful set of 1,087 highly conserved CpGs led to highly significant enrichment p-values for gene sets involved in RNA processing, RNA splicing, and lipoprotein particle biosynthesis. Some of the top gene families of these conserved probes include RBM and LDLR. For example, for ontology class "MSigDB Cancer Neighborhood" we find $P$=5.2x10$^{\mathbf{-19}}$ for "Neighborhood of SMC1L1",

$P=2.67x10^{-18}$ for "Neighborhood of TDG", $P=1.57x10^{-16}$ for "Neighborhood of XRCC5". Highly significant GO Biological Processes include RNA processing ($P=1.56x10^{-17}$), RNA binding ($P=5.90x10^{-16}$), mRNA processing ($P=1.15x10^{-14}$), and RNA splicing ($P=3.9x10^{-11}$). However, these enrichments are quite distinct from those observed for the EWAS of age. RNA splicing and processing only showed a weak significance ($P=0.05–1.4x10^{-3}$) in hypomethylated age-related CpGs.. In summary, we did not observe any overlap between the top enrichment terms for the age-related CpGs with those from highly conserved regions (or those from a random set of CpGs). A detailed enrichment analysis of all the CpGs on the mammalian array can be found in Arneson et al[61].

GREAT was not explicitly designed to adjust for the issue of certain CpG's having more power to detect association based on working in more species, but it appears not to be driving categories of enrichment for age. Overall, our sensitivity analysis of the enrichment study demonstrates that GREAT analysis adjusted for potential biases arising from the design of the mammalian array and protected us against spurious associations.

**Supplementary Note 4: Enrichment analysis for overlap between EWAS of mammalian age associated genes and large-scale GWAS associated ages of complex traits**

We related our EWAS of age results in mammals with a broad category of human genome wide association studies (GWAS) studies: anthropometric traits, behavioral phenotypes, cognitive related traits, fetal growth traits, inflammatory diseases, lipid panel outcomes, metabolic outcomes and diseases, neurodegenerative and neuropsychiatric disorders, longevity, reproductive aging and other age related phenotypes including DNA methylation based biomarkers. All GWAS results are based on meta-analysis across large-scale human studies. For instance, GWAS of anthropometric traits involved more than 200k individuals from multiple ethnic groups, conducted by the GIANT consortium, https://portals.broadinstitute.org/collaboration/giant/index.php/GIANT_consortium.

We used the MAGENTA software[62] to define gene level p-values for the human GWAS results from a total of 102 large-scale GWAS results [63-81]. For each GWAS, we focused on the top 2.5% genes for downstream enrichment analysis. Other thresholds would lead to qualitatively similar results. The EWAS-GWAS enrichment analysis was based on genomic-region hypergeometric analysis as described in **Methods**. The

GWAS summary datasets were downloaded from OpenGWAS [82-84] (https://gwas.mrcieu.ac.uk/) or obtained from the corresponding study groups. Citations to the respective scientific papers are provided below.

| Index | Hg | Category | Trait | Ethnicity | Sex | PMID |
|---|---|---|---|---|---|---|
| 1 | hg19 | Neurodegenerative disorder | Age-related Macular degeneration (AMD) | EUR+ASN | All | 23455636 |
| 2 | hg19 | Neurodegenerative disorder | AMD Geographic Atrophy | EUR+ASN | All | 23455636 |
| 3 | hg19 | Neurodegenerative disorder | AMD Neovascular | EUR+ASN | All | 23455636 |
| 4 | hg19 | Neurodegenerative disorder | Alzheimer's disease | EUR | All | 24162737 |
| 5 | hg18 | Longevity | Longevity > 90 | EUR | All | 24688116 |
| 6 | hg18 | Longevity | Longevity > 85 | EUR | All | 24688116 |
| 7 | hg19 | Neurodegenerative disorder | Parkinson's disease | EUR | All | 19915575 |
| 8 | hg19 | Neuropsychiatric disorder | Schizophrenia | All | All | 25056061 |
| 9 | hg19 | Inflammatory diseases | IBD | EUR | All | 26192919 |
| 10 | hg19 | Inflammatory diseases | IBD Crohn's disease | EUR | All | 26192919 |
| 11 | hg19 | Inflammatory diseases | IBD Ulcerative colitis | EUR | All | 26192919 |
| 12 | hg18 | Neuropsychiatric disorder | Bipolar disorder | All | All | 21926972 |
| 13 | hg18 | Neuropsychiatric disorder | ADHD | All | All | 20732625 |
| 14 | hg18 | Neuropsychiatric disorder | Major depression disorder | EUR | All | 22472876 |
| 15 | hg18 | Metabolic outcomes and diseases | Type 2 diabetes | EUR | All | 22885922 |
| 16 | hg18 | Metabolic outcomes and diseases | Fasting glucose | EUR | All | 22581228 |
| 17 | hg18 | Metabolic outcomes and diseases | Fasting insulin | EUR | All | 22581228 |
| 18 | hg18 | GIANT Body fat distribution | Hip AllAncestries | ALL | M&F | 25673412 |
| 19 | hg18 | GIANT Body fat distribution | Hip EUR | EUR | M&F | 25673412 |
| 20 | hg18 | GIANT Body fat distribution | Hip AllAncestries(Males) | ALL | M | 25673412 |
| 21 | hg18 | GIANT Body fat distribution | Hip EUR (Males) | EUR | M | 25673412 |

| | | | | | | |
|---|---|---|---|---|---|---|
| 22 | hg18 | GIANT Body fat distribution | Hip AllAncestries(Females) | ALL | F | 25673412 |
| 23 | hg18 | GIANT Body fat distribution | Hip EUR (Females) | EUR | F | 25673412 |
| 30 | hg18 | GIANT Body fat distribution | Waist circumference AllAncestries | ALL | M&F | 25673412 |
| 31 | hg18 | GIANT Body fat distribution | Waist circumference EUR | EUR | M&F | 25673412 |
| 32 | hg18 | GIANT Body fat distribution | Waist circumference AllAncestries(Males) | ALL | M | 25673412 |
| 33 | hg18 | GIANT Body fat distribution | Waist circumference EUR (Males) | EUR | M | 25673412 |
| 34 | hg18 | GIANT Body fat distribution | Waist circumference AllAncestries(Females) | ALL | F | 25673412 |
| 35 | hg18 | GIANT Body fat distribution | Waist circumference EUR (Females) | EUR | F | 25673412 |
| 42 | hg18 | GIANT Body fat distribution | Waist to hip ratio AllAncestries | ALL | M&F | 25673412 |
| 43 | hg18 | GIANT Body fat distribution | Waist to hip ratio EUR | EUR | M&F | 25673412 |
| 44 | hg18 | GIANT Body fat distribution | Waist to hip ratio AllAncestries(Males) | ALL | M | 25673412 |
| 45 | hg18 | GIANT Body fat distribution | Waist to hip ratio EUR (Males) | EUR | M | 25673412 |
| 46 | hg18 | GIANT Body fat distribution | Waist to hip ratio AllAncestries(Females) | ALL | F | 25673412 |
| 47 | hg18 | GIANT Body fat distribution | Waist to hip ratio EUR (Females) | EUR | F | 25673412 |
| 54 | hg18 | GIANT BMI & Height | BMI | EUR | All | 25673413 |
| 55 | hg18 | GIANT BMI & Height | Height | EUR | All | 20881960 |
| 56 | hg19 | Neurodegenerative disorder | Frontotemporal dementia | EUR | All | 24943344 |
| 57 | hg19 | Neurodegenerative disorder | FTD Behavioral variant | EUR | All | 24943344 |
| 58 | hg19 | Neurodegenerative disorder | FTD with motor neuron disease | EUR | All | 24943344 |
| 59 | hg19 | Neurodegenerative disorder | FTD progressive non-fluent aphasia | EUR | All | 24943344 |
| 60 | hg19 | Neurodegenerative disorder | FTD semantic dementia | EUR | All | 24943344 |
| 61 | hg19 | Neurodegenerative disorder | Huntington's disease age onset | EUR | All | 26232222 |
| 62 | hg19 | Behavioral phenotype | Educational attainment | EUR | All | 27225129 |
| 63 | hg19 | Behavioral phenotype | Educational attainment (Males) | EUR | All | 27225129 |
| 64 | hg19 | Behavioral phenotype | Educational attainment (Females) | EUR | All | 27225129 |

| | | | | | | |
|---|---|---|---|---|---|---|
| 65 | hg18 | Reproductive aging | Age at menarche | EUR | All | 25231870 |
| 66 | hg18 | Reproductive aging | Age at menopause | EUR | All | 26414677 |
| 67 | hg18 | Lipid panel outcomes | HDL | | All | 24097068 |
| 68 | hg18 | Lipid panel outcomes | LDL | | All | 24097068 |
| 69 | hg18 | Lipid panel outcomes | Total cholesterol | | All | 24097068 |
| 70 | hg18 | Lipid panel outcomes | Triglyceride | | All | 24097068 |
| 71 | hg18 | Reproductive aging | Leukocyte telomere length | EUR | All | 23535734 |
| 72 | hg19 | DNAm biomarkers | AgeAccelGrim EUR | EUR | All | 34187551 |
| 73 | hg19 | DNAm biomarkers | DNAmGranAdjustedAge EUR | EUR | All | 34187551 |
| 74 | hg19 | DNAm biomarkers | AgeAccelHannum EUR | EUR | All | 34187551 |
| 75 | hg19 | DNAm biomarkers | DNAmPAI1AdjAge EUR | EUR | All | 34187551 |
| 76 | hg19 | DNAm biomarkers | IEAA EUR | EUR | All | 34187551 |
| 77 | hg19 | DNAm biomarkers | AgeaccelPhenoAge EUR | EUR | All | 34187551 |
| 78 | hg19 | DNAm biomarkers | AgeAccelGrim AFR | AFR | All | 34187551 |
| 79 | hg19 | DNAm biomarkers | DNAmGranAdjustedAge AFR | AFR | All | 34187551 |
| 80 | hg19 | DNAm biomarkers | AgeAccelHannum AFR | AFR | All | 34187551 |
| 81 | hg19 | DNAm biomarkers | DNAmPAI1AdjAge AFR | AFR | All | 34187551 |
| 82 | hg19 | DNAm biomarkers | IEAA AFR | AFR | All | 34187551 |
| 83 | hg19 | DNAm biomarkers | AgeaccelPhenoAge AFR | AFR | All | 34187551 |
| 84 | hg19 | DNAm biomarkers | AgeAccelGrim All | EUR+AFR | All | 34187551 |
| 85 | hg19 | DNAm biomarkers | DNAmGranAdjustedAge All | EUR+AFR | All | 34187551 |
| 86 | hg19 | DNAm biomarkers | AgeAccelHannum All | EUR+AFR | All | 34187551 |
| 87 | hg19 | DNAm biomarkers | DNAmPAI1AdjAge All | EUR+AFR | All | 34187551 |
| 88 | hg19 | DNAm biomarkers | IEAA All | EUR+AFR | All | 34187551 |
| 89 | hg19 | DNAm biomarkers | AgeAccelPhenoAge All | EUR+AFR | All | 34187551 |
| 90 | hg19 | Longevity | Father's attained age | EUR | All | 29227965 |
| 91 | hg19 | Longevity | Mother's attained age | EUR | All | 29227965 |
| 92 | hg19 | Longevity | Parental attained age | EUR | All | 29227965 |
| 93 | hg19 | Age related phenotype | Atrial fibrillation | EUR | All | 30061737 |
| 94 | hg19 | Neurodegenerative disorder | Alzheimer's disease | EUR | All | 30617256 |
| 95 | hg19 | Cognitive related | Intelligence | EUR | All | 29942086 |
| 96 | hg19 | Reproductive aging | AgeAtMenarche | EUR | All | 28436984 |
| 97 | hg19 | Neurodegenerative disorder | Huntington's disease motor progression | EUR | All | 28642124 |

| | | | | | | |
|---|---|---|---|---|---|---|
| 98 | hg19 | Fetal growth | Birth length | EUR | All | 34282336 |
| 99 | hg19 | Fetal growth | Infant Ponderal index | EUR | All | 34282336 |
| 100 | hg19 | Fetal growth | Birth weight | EUR | All | 34282336 |
| 101 | hg19 | Fetal growth | Birth weight fatherGenome | EUR | All | 34282336 |
| 102 | hg19 | Fetal growth | Birth weight motherGenome | EUR | All | 34282336 |
| EUR: Europeans; AFR: Africans; ASN: Asians. | | | | | | |

#### Supplementary Note 5: Human cohorts

##### Framingham Heart Study Cohort (FHS)

The FHS cohort[85] and is a large-scale longitudinal study started in 1948, initially investigating the common factors of characteristics that contribute to cardiovascular disease (CVD), https://www.framinghamheartstudy.org/index.php. The study at first enrolled participants living in the town of Framingham, Massachusetts, who were free of overt symptoms of CVD, heart attack or stroke at enrollment. In 1971, the study started FHS Offspring Cohort to enroll a second generation of the original participants' adult children and their spouses (n= 5124) for conducting similar examinations[86]. Participants from the FHS Offspring Cohort were eligible for our study if they attended eighth examination cycles and consented to having their molecular data used for study. We used the 2,544 participants with available DNA methylation profiles (measured at exam 8), from the group of Health/Medical/Biomedical (IRB, MDS) consent. The FHS data are available in dbGaP (accession number: phs000363.v16.p10 and phs000724.v2.p9). Deaths among the FHS participants that occurred prior to January 1, 2013 were ascertained using multiple strategies, including routine contact with participants for health history updates, surveillance at the local hospital and in obituaries of the local newspaper, and queries to the National Death Index. Death certificates, hospital and nursing home records prior to death, and autopsy reports were requested. When cause of death was undeterminable, the next of kin were interviewed. The date and cause of death were reviewed by an endpoint panel of 3 investigators.

**DNA methylation quantification** Peripheral blood samples were collected at the 8th examination. Genomic DNA was extracted from buffy coat using the Gentra Puregene DNA extraction kit (Qiagen) and bisulfite converted using EZ DNA Methylation kit (Zymo Research Corporation). DNA methylation quantification was conducted in two laboratory batches using the Illumina Infinium HumanMethylation450 array (Illumina).

Methylation beta values were generated using the Bioconductor *minfi* package with Noob background correction[87].

**Women's Health Initiative (WHI)**

The Women's Health Initiative is a national landmark study that enrolled postmenopausal women aged 50-79 years into the clinical trials (CT) or observational study (OS) cohorts between 1993 and 1998[88,89]. We included 2,017 WHI participants from "*Broad Agency Award 23*" (WHI BA23) with available phenotype and DNA methylation array data: 2,107 women from "*Broad Agency Award 23*" (WHI BA23). WHI BA23 focuses on identifying miRNA and genomic biomarkers of coronary heart disease (CHD), integrating the biomarkers into diagnostic and prognostic predictors of CHD and other related phenotypes. The study span three WHI sub-cohorts including GARNET, WHIMS and SHARe.

**DNA methylation quantification**

In brief, bisulfite conversion using the Zymo EZ DNA Methylation Kit (Zymo Research, Orange, CA, USA) as well as subsequent hybridization of the HumanMethylation450k Bead Chip (Illumina, San Diego, CA), and scanning (iScan, Illumina) were performed according to the manufacturers protocols by applying standard settings. DNA methylation levels (β values) were determined by calculating the ratio of intensities between methylated (signal A) and un-methylated (signal B) sites. Specifically, the β value was calculated from the intensity of the methylated (M corresponding to signal A) and un-methylated (U corresponding to signal B) sites, as the ratio of fluorescent signals $\beta = Max(M,0)/[Max(M,0)+Max(U,0)+100]$. Thus, β values range from 0 (completely un-methylated) to 1 (completely methylated).

**Supplementary Note 6: Murine anti-aging studies**

Universal clocks 2&3 were evaluated in mouse experiments: (1) Snell dwarf mice (n=95), 2) growth hormone receptor knock-out experiment 1 (GHRKO, n=71 samples), (3) GHRKO experiment 2 (n=96 samples), (4) Tet3 KO (n=63), and (5) calorie restriction (n=95). Below we summarize each experiment. More details of for experiments 2,4 and 5 can be found in **Supplementary Note 1**.

**Snell dwarf experiment**

The Snell dwarf mice for loss of function at Pit1 were generated at the University of Michigan (Richard A. Miller). Snell dwarf mice, which lack growth hormone, thyroid stimulating hormone, and prolactin, have approximately 30% - 40% longer lifespan (http://www.richmillerlab.com/long-lived-mutants). The extended lifespan related to Snell dwarf mice was previously studied by Flurkey et al. [90]

**GHRKO experiment 1**

The GHRKO mice (GHR -/- dwarfs) were generated at the University of Michigan (Richard A. Miller). The experiment was based on full-body growth hormone receptor knockout (GHR-KO) as described in details by Coschigano et al., 2003[91].

**GHRKO experiment 2**

The GHRKO mice (GHR -/- dwarfs) were generated at the University of Michigan (Richard A. Miller). The experiment only knocked out the growth hormone receptors in livers.

**Tet gene knock-out study**

*Tet1* and *Tet2* knockout mice were purchased from the Jackson Laboratory (*Tet1* KO: Strain #:017358, B6;129S4-$Tet1^{tm1.1Jae}$/J; *Tet2* KO: Strain #:023359, B6(Cg)-$Tet2^{tm1.2Rao}$/J). The *Tet3* knockout mice were generated at the Jackson Laboratory using CRISPR technology with deletion of exon 4. For each line, the heterozygous knockout mice were bred with C57BL/6J (JAX 000664) to set up the colony. All mice were maintained and bred under standard conditions consistent with National Institutes of Health guidelines and approved by the University of California, Los Angeles Institutional Animal Care and Use Committees. The cages were maintained on a 12:12 light/dark cycle, with food and water provided *ad libitum*. Mice were euthanized by cervical dislocation; cortical and striatal tissues were then quickly dissected out and flash frozen on dry ice until used for DNA extraction

**Calorie restriction**

All the mice in the calorie restriction (CR) study were male from C57BL/6J inbred strain.

C57BL/6J male mice (N=95) were obtained from the Mouse Breeding Core, Wakeland lab, UT Southwestern Medical Center, Dallas, TX, USA. All the mice have the same ages (1.57 years). DNA methylation arrays were profiled in liver samples.

**Supplementary Note 7: R code for universal pan-mammalian clocks.**

The following R code requires one input data (info in the R code) and three csv files that list three universal clocks models in csv format, which can be copied from Data3.1—Data3.3 in Supplementary Data. The info file needs to include ID variable, gestation time in years (), age at sexual maturity in years, and maximum lifespan in years. The variables names in the demo R code are SampleID, GestationTimeInYears, averagedMaturity.yrs, and maxAgeCaesar, respectively.

#
#Rcode
#

```
#

info=readRDS('mydata.Rds')#This dataset include species characters and CpGs
glmnet.csv=c('clock1.csv',
             'clock2.csv',
             'clock3.csv')#S3.1 to S3.3  tables in SupplmentaryData
beta.name=c('beta_clock1','beta_clock2','beta_clock3')
y.name=c('Y.pred1','Y.pred2','Y.pred3')
age.name=c('DNAmAgeClock1','DNAmAgeClock2','DNAmAgeClock3')

#clock2
F2_antitrans_clock2<-function(y,y.maxAge,y.gestation,const=1){
  x0=const*exp(-exp(-1*y))
  x1=x0*(y.maxAge+y.gestation)
  x=x1-y.gestation
  x
}
#
#clock3
#
F1_logli <- function(age1, m1, m2 = m1, c1=1){
  ifelse(age1 >= m1, (age1-m1)/m2 , c1*log((age1-m1)/m2/c1 +1) )
}
#RelativeAdultAge
F2_revtrsf_clock3 <- function(y.pred, m1, m2 = m1, c1=1){
  ifelse(y.pred<0, (exp(y.pred/c1)-1)*m2*c1 + m1, y.pred*m2+m1 )
}

#############
# The `loglifn` function shows how to calculate m1 for the transformation
# It is the `a_Logli` in the function

F3_loglifn = function(dat1,b1=1,max_tage = 4,
                      c1=5, c2 = 0.38, c0=0){
  n=nrow(dat1)

  age1 =
(dat1$maxAgeCaesar+dat1$GestationTimeInYears)/(dat1$averagedMaturity.yrs+dat1$GestationTimeInYea
rs)

  a1 = age1/(1+max_tage)
  dat1$a1_Logli = a1 #x/m1 in manuscript

  a2 = (dat1$GestationTimeInYears + c0)/(dat1$averagedMaturity.yrs)
  dat1$a_Logli = a_Logli = c1*a2^c2
  #m=5*(G/ASM)^0.38 from regression analysis/formula(7)


  x = dat1$Age + dat1$GestationTimeInYears
  t2 = dat1$averagedMaturity.yrs*b1 + dat1$GestationTimeInYears
  x2 = x/t2 #### log(x/t2)
  y = F1_logli(x2, a_Logli, a_Logli)

  dat1$LogliAge <- y
  return(dat1)
}

#
info$Intercept=1
MYMAX=1.3
info$HighmaxAgeCaesar=MYMAX*info$maxAgeCaesar
info$HighmaxAgeCaesar[info$SpeciesLatinName=='Homo
sapiens']=info$maxAgeCaesar[info$SpeciesLatinName=='Homo sapiens']
info$HighmaxAgeCaesar[info$SpeciesLatinName=='Mus
musculus']=info$maxAgeCaesar[info$SpeciesLatinName=='Mus musculus']
#
#predict RelativeAge
```

```
#
for(k in 1:3){
glmnet=read.csv(glmnet.csv[k])
glmnet$beta=glmnet[,beta.name[k]]
glmnet$var[1]=ifelse(glmnet$var[1]=="(Intercept)",'Intercept',glmnet$var[1])
temp=as.matrix(subset(info,select=as.character(glmnet$var)))
info[,y.name[k]]=as.numeric(as.matrix(subset(info,select=as.character(glmnet$var)))%*%glmnet$bet
a)
}
#(1) Clock 1
info[,age.name[1]]=exp(info[,y.name[k]])-2
#(2) Clock 2
info$DNAmRelativeAge=exp(-exp(-1*info[,y.name[2]]))
info[,age.name[2]]=
F2_antitrans_clock2(info[,y.name[2]],info$HighmaxAgeCaesar,info$GestationTimeInYears,const=1)
#(3) Clock 3
info=F3_loglifn(info)#to compute m estimate for tuning point in the log-linear transformation
info$m1=info$a_Logli
info$DNAmRelativeAdultAge=F2_revtrsf_clock3(info[,y.name[3]], info$m1)
info[,age.name[3]]<-
  info$DNAmRelativeAdultAge *(info$averagedMaturity.yrs + info$GestationTimeInYears) -
info$GestationTimeInYears
#
#final output
#
output=subset(info,select=c('SampleID','Age','DNAmRelativeAge','DNAmRelativeAdultAge',age.name))
```
